# A novel expectation-maximization approach to infer general diploid selection from time-series genetic data

**DOI:** 10.1101/2024.05.10.593575

**Authors:** Adam G. Fine, Matthias Steinrücken

## Abstract

Detecting and quantifying the strength of selection is a main objective in population genetics. Since selection acts over multiple generations, many approaches have been developed to detect and quantify selection using genetic data sampled at multiple points in time. Such time series genetic data is commonly analyzed using Hidden Markov Models, but in most cases, under the assumption of additive selection. However, many examples of genetic variation exhibiting non-additive mechanisms exist, making it critical to develop methods that can characterize selection in more general scenarios. Here, we extend a previously introduced expectation-maximization algorithm for the inference of additive selection coefficients to the case of general diploid selection, in which the heterozygote and homozygote fitness are parameterized independently. We furthermore introduce a framework to identify bespoke modes of diploid selection from given data, a heuristic to account for variable population size, and a procedure for aggregating data across linked loci to increase power and robustness. Using extensive simulation studies, we find that our method accurately and efficiently estimates selection coefficients for different modes of diploid selection across a wide range of scenarios; however, power to classify the mode of selection is low unless selection is very strong. We apply our method to ancient DNA samples from Great Britain in the last 4,450 years, and detect evidence for selection in six genomic regions, including the well-characterized LCT locus. Our work is the first genome-wide scan characterizing signals of general diploid selection.

**Author Summary:** Natural selection increases the likelihood that beneficial genetic variants are passed from parent to offspring and thus forms the basis of genetic adaptation to novel environments. Genomic data sampled at multiple timepoints, such as genetic material extracted from ancient remains (ancient DNA) or data from evolve and resequence experiments, can enable more precise identification of genetic variants subject to selective pressure than contemporary samples alone. However, most methods for identifying genetic variation under selection focus on additive selection, where the fitness of the heterozygote is exactly intermediate between the homozygotes. Leveraging genetic data at multiple timepoints, we develop a method to detect additive and non-additive selection as well as to infer the most likely dominance mechanism. We apply our methods to a dataset of ancient DNA from Great Britain dated less than 4,450 years before present and identify six regions with signals of recent selection, including one at the TFR2 locus that has not been previously reported as a target of selection. Our work enables more accurate quantification of non-additive selection dynamics and can be used to test more complex models of selection.

## 1 Introduction

Genetic variation that confers a fitness advantage to an organism over its peers tends to increase in frequency in the population over time until eventual fixation, if it is not lost to genetic drift. This stochastic process of selection ultimately forms the basis of adaptation. Detecting evidence of selection and quantifying its strength is thus a fundamental problem in evolutionary biology, with applications ranging from finding mutations critical to early hominid evolution (Bustamante et al. 2005) to predicting tumor growth (Bignell et al. 2010). In population genetics, many methods to detect signatures of past selective events in contemporary population genomic data have thus been developed (Nielsen 2005; Vitti et al. 2013; Lachance and Tishkoff 2013).

However, since selection acts over multiple generations, genetic data observed at several timepoints throughout allows for more accurate quantification of selective processes than present-day samples alone. Recent technological advances have enabled researchers to collect such time series genetic data genome-wide. One main source of time series genetic data is ancient DNA (aDNA), that is, genetic material extracted from deceased individuals in humans or other species (Orlando et al. 2021). Next-generation sequencing has enabled collecting genetic data from large numbers of ancient samples, particularly through the development of techniques such as hybridization enrichment (Hofreiter et al. 2015). Another major source of temporal genetic data are experimental evolution studies (Barghi et al. 2019). Contemporary experimental evolution studies use next-generation sequencing technologies on several biological replicates in evolve and resequence (E&R) experiments to obtain high-quality estimates of temporal allele frequency changes at many loci throughout the genome (Schlötterer et al. 2015). These datasets present unprecedented opportunities to detect and characterize the adaptive processes that shape genomic variation (Malaspinas 2016; Dehasque et al. 2020).

Observing the true underlying population allele frequency trajectory as it changes over time would allow for highly accurate characterization of the underlying selective processes. However, in data obtained in practice, genetic variation is often only assessed for a set of individuals sampled at a finite number of time points. Quantifying the strength of selection therefore involves modeling the action of selection, genetic drift, and other population genetic processes on the unobserved trajectory of the population allele frequency, and considering sampled data as imprecise observations of this underlying trajectory.

A commonly used framework for analysis of time series data that readily accommodates this uncertainty are Hidden Markov Models (HMMs, Bollback et al. 2008). In these HMMs, the underlying population allele frequency evolves according to a Markov process, the Wright–Fisher model, and samples are modeled as binomial observations given the underlying population allele frequency. This HMM framework has been used to estimate a variety of parameters: The additive selection coefficient *s* (Bollback et al. 2008), the time at which a beneficial mutation arose (Malaspinas et al. 2012), the effective population size *N*_*e*_ (Wang 2001), or the rate of sequencing errors (Ferrer-Admetlla et al. 2016). Within this HMM framework, Mathieson and McVean (2013) introduced an expectation-maximization (EM) approach, which can be used to estimate additive selection coefficients, as well as migration rates between sub-populations. For reviews of HMM-based approaches to estimate selection coefficients and comparisons between methods in, respectively, aDNA and E&R analyses, see Tataru et al. (2017) and Vlachos et al. (2019).

Most implementations of the aforementioned HMM approach presented in the literature to date are designed to only detect additive selection, but many examples of genetic variation exhibiting non-additive mechanisms exist. In humans, non-additive targets of selection range from the classical case of the heterozygote advantage conferred by one copy of the sickle-cell allele (Gemmell and Slate 2006; Hedrick 2012) to recent evidence of pervasive dominance in an analysis of data from the UK Biobank (Palmer et al. 2023). Additionally, stabilizing selection on complex traits, which is believed to be wide-spread in humans (Sanjak et al. 2018), manifests as underdominant selective dynamics at the loci affecting the trait (Barton 1986; de Vladar and Barton 2014; Simons et al. 2018; Koch et al. 2024).

Here, we extend the EM approach of Mathieson and McVean (2013) to estimate selection coefficients under a general diploid model, that is, when the fitness values of homozygous and heterozygous genotypes are independently parameterized. The use of an EM method to estimate the selection parameters maximizing the likelihood iteratively allows for better scaling to more than two parameters, compared to grid-search based methods such as, for example, Cheng and Steinrücken (2023). For example, the use of the EM algorithm allows us to simultaneously estimate diploid selection coefficients and the parameters characterizing the initial frequency of an allele, whereas estimating these parameters using a grid-search would be challenging.

We furthermore develop a novel framework for identifying the best-fitting mode of selection for a given temporal dataset. While other methods to estimate general diploid selection coefficients have been presented in the literature (Steinrücken et al. 2014; Foll et al. 2015; Schraiber et al. 2016; Iranmehr et al. 2017; Taus et al. 2017; Cheng and Steinrücken 2023), they do not explicitly address the statistical problem of distinguishing between different modes of selection. Furthermore, none of these methods have been applied to genome-wide data from human populations. To our knowledge, our analysis is the first that characterizes recent general diploid selection in humans from ancient DNA data on a full-genome scale.

The remainder of this article is organized as follows. In Section 2, we outline our iterative EM algorithm for efficiently estimating general diploid selection coefficients, as well as the statistical procedure for inferring the most likely mode of diploid selection. In Section 3.1, we apply our algorithm and inference framework in a wide range of simulated scenarios to assess its accuracy. We find that our method is generally well-powered to detect selection and estimate its strength, however, power to classify the mode of selection is limited. Moreover, in Section 3.1.6, we introduce a procedure for estimating a constant population size from temporal data. In Section 3.2, we then perform a genome-wide scan for recent diploid selection in the human genome, using publicly available ancient DNA data from individuals that lived in Great Britain in the last 4,450 years (Mallick et al. 2024, version 54.1), introduce a procedure to aggregate p-values across linked loci, and discuss six genomic regions that show signals of recent selection. In Section 3.3, we also apply our method to a locus involved in horse coat coloration (Ludwig et al. 2009) to demonstrate the utility of our method when exploring non-additive scenarios. Lastly, we discuss future directions in Section 4. Our method EMSel (EM algorithm for detecting Selection) and the scripts to generate the figures in this manuscript are available online at https://github.com/steinrue/EMSel.

## 2 Methods

### 2.1 Parameterizing general diploid selection

Throughout this article, we consider selection acting at a given biallelic locus in a diploid population of constant size *N*_*e*_. The dynamics of the population allele frequency at this locus can be described by the discrete Wright–Fisher model, where we denote by *A* and *a* the two alleles at the locus, and by *Y*_*t*_ ∈ [0, 1] and 1 − *Y*_*t*_ the random population-level frequency of the *A* and *a* allele in generation *t* ∈ {1, … *T*}, respectively. Suppose that the relative fitness of individuals with genotypes *AA, Aa*, and *aa* is 1 + *s*_2_, 1 + *s*_1_, and 1, respectively. Given these fitness values, if the frequency of *A* alleles in the current generation is *Y*_*t*_ = *p*, then the allele frequency in the next generation *Y*_*t*+1_ is given by 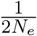 times a binomially distributed random variable with 2*N*_*e*_ draws and success probability *p*^′^ := *p* + *p*(1 − *p*) *s*_1_(1 − 2*p*) + *s*_2_*p* for small *s*_1_ and *s*_2_. This can further be approximated by a normal distribution, where 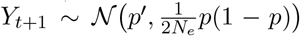, which is commonly referred to as the Wright–Fisher diffusion (Ewens 2004, Chapter 5.3).

We use the term *general diploid selection* for the case of arbitrary *s*_1_ *>* −1 and *s*_2_ *>* −1, that is, the fitness values for the homozygotes and heterozygotes are independently parametrized. Many bespoke modes of selection correspond to constrained diploid selection, where the possible values of *s*_1_ and *s*_2_ are restricted. We consider the following modes of selection:

- **Additive**: 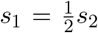. The relative fitness is proportional to the number of copies of the *A* allele. Also referred to as **haploid** or **genic** selection.
- **Dominant**: *s*_1_ = *s*_2_. Any number of copies of the *A* allele confers the full fitness effect.
- **Recessive**: *s*_1_ = 0. Only individuals homozygous for the *A* allele have a fitness effect.
- **Overdominance**: *s*_1_ *>* max{0, *s*_2_}. Heterozygous individuals have the highest fitness.
- **Underdominance**: *s*_1_ *<* max{0, *s*_2_}. Heterozygous individuals have the lowest fitness.

*Additive, dominant, recessive* selection are one-parameter modes, with *s* := *s*_2_ = 2*s*_1_, *s* := *s*_1_ = *s*_2_, and *s* := *s*_2_ (*s*_1_ = 0), respectively. *Over-* and *underdominance* are two-dimensional sub-spaces, but we frequently use the version *s* := *s*_1_ (*s*_2_ = 0) as a one-parameter selection mode that describes complete *over-* or *underdominance*. We also refer to the combination of the latter two as *heterozygote difference*. While the one-parameter modes for complete *over-* and *underdominance* are more convenient for simulations, combining them into the one-dimensional mode *heterozygote difference* when inferring parameters avoids statistical artifacts. In Figure 1, we show a diagram of the different modes of selection as sub-spaces of the full two-dimensional parameter space of general diploid selection.

**Figure 1:**
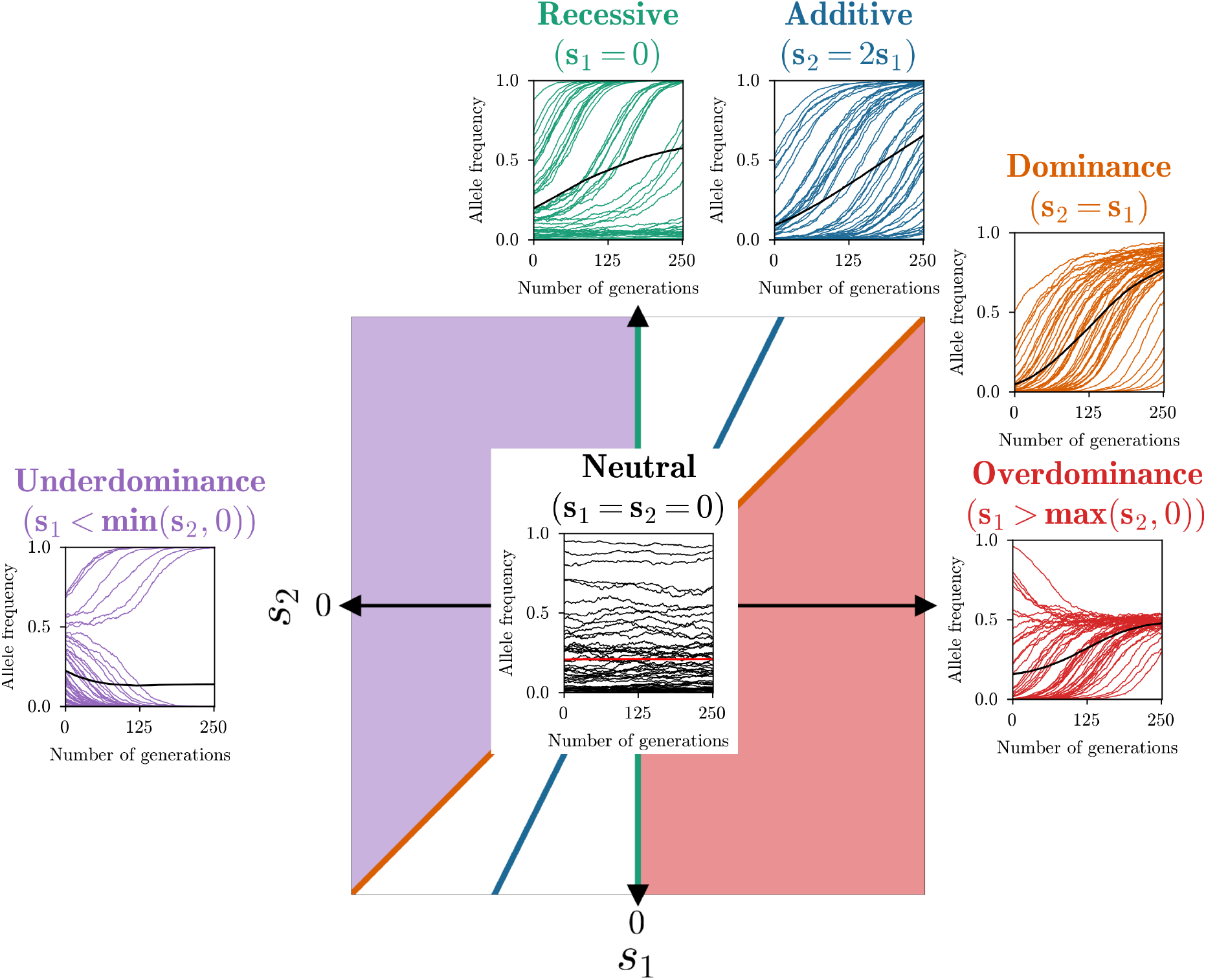
Two-dimensional space of general diploid selection and the five bespoke selection modes we consider: *Additive, dominant, recessive, over-* and *underdominance*. Colors indicate the sub-space of the respective mode. 50 replicates simulated under each mode, as well as their mean, are plotted to illustrate the characteristic shapes of the trajectories in each mode.

### 2.2 Hidden Markov Model for inferring diploid selection coefficients

#### 2.2.1 Derivation of the EM-HMM algorithm

To derive our novel method for inferring general diploid selection coefficients from sampled time series genetic data we extend the method developed by Mathieson and McVean (2013) for *additive* selection. If the exact frequency trajectory of the focal allele *p*_1_, …, *p*_*T*_ ∈ [0, 1] over *T* generations is known, then the normal approximation to the Wright-Fisher process can be used to define the likelihood of this trajectory as

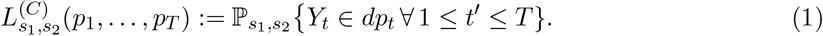

Here, we indicate the parameters of interest, the selection coefficients *s*_1_ and *s*_2_, in the subscript, and the superscript (*C*) indicates the model where the population allele frequency in each generation takes values in the continuous range [0, 1]. Without loss of generality, the focal allele is the *A* allele. Maximizing this likelihood yields a maximum likelihood estimator (MLE) of the selection coefficients. In the case of *additive* selection, this estimator has been presented by Watterson (1982). In Sections S.1.1 and S.1.2 in the Supplementary Material, we revisit Watterson’s formula for the *additive* MLE and extend the result to general diploid selection, which yields the estimators

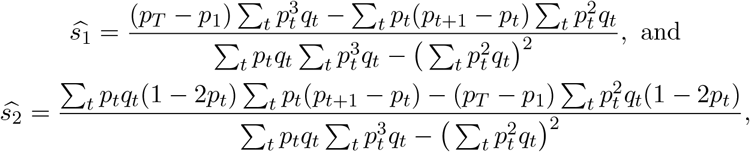

with *q*_*t*_ := 1 − *p*_*t*_.

However, as discussed in Section 1, the data often consists of a given number of individuals sampled from the population at certain points in time, and thus the full trajectory of the population allele frequency is in general not known. To efficiently integrate over this uncertainty, Bollback et al. (2008) introduced an HMM framework, where the population allele frequency at a given time is the hidden state, evolving according to the Wright-Fisher model, and the sampled genotypes are the observations. Specifically, assume that we have sampled the population at times 1 *≤ t*_1_, …, *t*_*K*_ *≤ T*. At each timepoint, the data 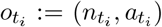 consists of the number of haplotypes sampled at this time 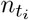, and the number of observed focal alleles 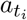. For convenience, we set *n*_*t*_ := 0 and *a*_*t*_ := 0 at times where no data is observed, and we denote the random variable associated with sampling the data at *t* by *O*_*t*_. In addition, we discretize the population allele frequency into *M* hidden states to allow efficient computation: 𝒢:= {*g*_0_ = 0, *g*_1_, …, *g*_*M*−1_ = 1}, with *g*_*i*_ ∈ [0, 1], and *g*_*i*−1_ *< g*_*i*_. The hidden state in generation *t* is then the discretized population allele frequency at *t*, which we denote by *F*_*t*_ ∈ 𝒢.

To apply the standard HMM framework (Rabiner 1989; Bishop 2006, Ch. 13.2), we must define initial probabilities 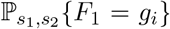, transition probabilities, and emission probabilities. For now, we assume that the initial probabilities are given. These can either be fixed or estimated, and we provide details on an estimation procedure in Section 2.2.3. For the transition probabilities from hidden state *g*_*i*_ to hidden state *g*_*j*_, we use the normal approximation to the Wright-Fisher process and define

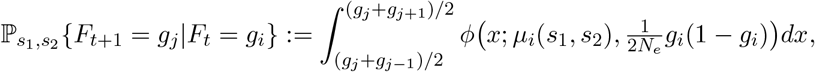

where *µ*_*i*_(*s*_1_, *s*_2_) := *g*_*i*_ + *g*_*i*_(1 − *g*_*i*_)(*s*_1_(1 − 2*g*_*i*_) + *s*_2_*g*_*i*_) for general diploid selection and *ϕ*(*x*; *µ, σ*^2^) denotes the density of a normal distribution with mean *µ* and variance *σ*. Here, we also set *g*_−1_ := −∞ and *g*_*M*_ := ∞, which assigns all the probability mass to transition outside of the interval [0, 1] to the respective boundary points. Lastly, the emission probabilities to observe *a*_*t*_ focal alleles in generation *t* are given by the binomial distribution

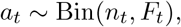

with *n*_*t*_ draws and success probability *F*_*t*_, the respective population allele frequency. Figure 2A depicts a schematic of this HMM.

**Figure 2:**
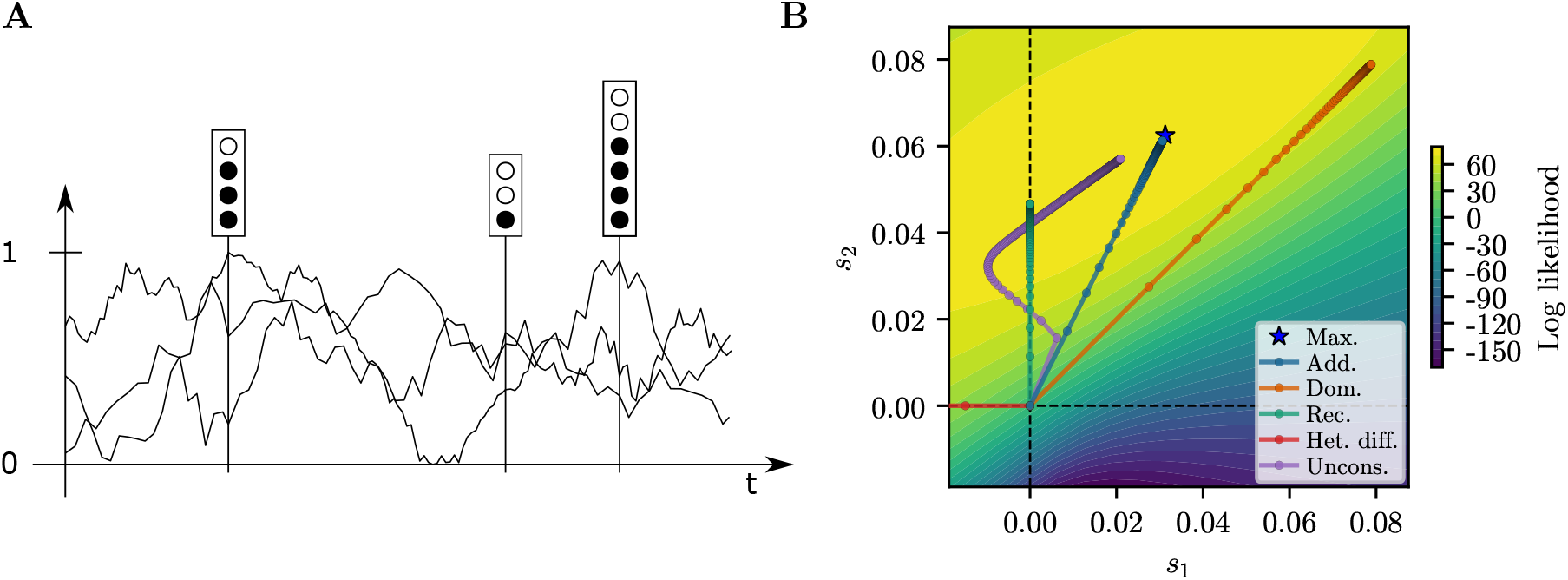
A) Schematic of an HMM. Each “stoplight” represents a haploid sample at the given time, with a certain number of focal alleles. Also plotted are three possible population allele frequency trajectories through the hidden state space. Trajectories with population frequencies closer to the frequencies in the samples are given more weight when computing the expected values in the E-step of the algorithm. B) Log-likelihood surface and path of the EM-HMM optimization under each mode for a given replicate simulated under *additive* selection.

The *forward-backward algorithm* (Rabiner 1989; Bishop 2006, Ch. 13.2) can then be used to obtain the posterior probability over the hidden states in generation *t*

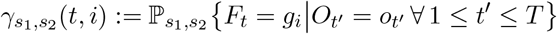

conditional on the observed data and given selection coefficients *s*_1_ and *s*_2_, and the joint posterior of the hidden states

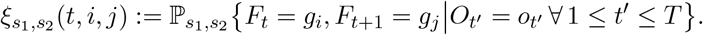

Based on these posterior distributions, posterior expectations

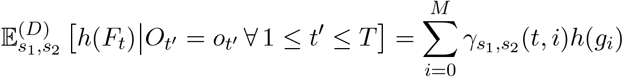

And

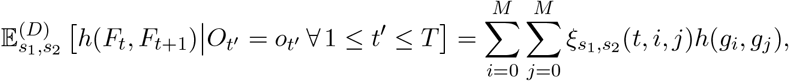

can be computed for arbitrary functions *h*(·) and *h*(·, ·) of the marginal frequency and joint marginal frequencies, respectively. Here, the superscript (*D*) indicates the use of the model with the discretized allele frequencies.

To find the maximum likelihood estimate (MLE) of the diploid selection coefficients 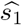 and 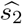 under this HMM, we use the EM algorithm (Rabiner 1989; Bishop 2006, Ch. 13.2), similar to Mathieson and McVean (2013) in the *additive* case. We refer to this algorithm as an EM-HMM algorithm. The algorithm starts with an initial parameter estimate 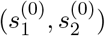. At iteration *k*, the algorithm then computes 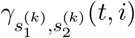 and 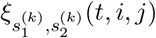 (E-step), and updates the parameter estimates by maximizing the conditional log-likelihood using

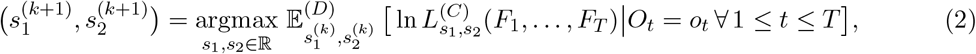

(M-step) until the estimates converge. Note that, under the binomial model, the emission probabilities are independent of *s*_1_ and *s*_2_ and do not need to be considered explicitly in this update step. Similar to the derivation of the MLE for a given trajectory from equation (1), equation (2) yields

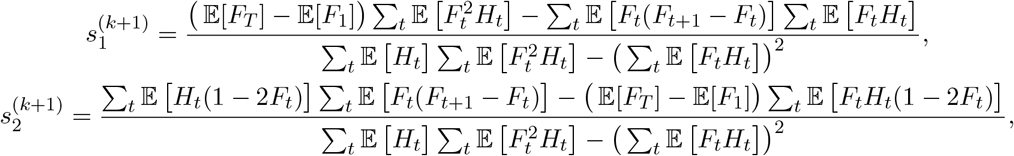

where *H*_*t*_ := *F*_*t*_(1 − *F*_*t*_), and we denote the discretized conditional expectation by

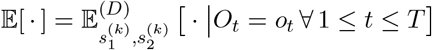

for brevity. See Section S.2.2 in the Supplementary Material for details of the derivation.

Note that this approach is not an exact EM algorithm, since it combines the conditional expectation 𝔼^(*D*)^[·] computed under the discretized model with maximization of the likelihood *L*^(*C*)^(·) in the continuous model, similar to Mathieson and McVean (2013). The reason for this hybrid approach is that while the conditional log-likelihood can be maximized in the continuous setting using the diploid generalization of Watterson’s MLE, the posterior expectations cannot readily be computed. On the other hand, in the discretized model, the posterior expectations can be computed, but the conditional log-likelihood cannot be maximized analytically. The hybrid approach, however, is computationally tractable and yields highly accurate estimates.

#### 2.2.2 Constrained optimization for bespoke selection modes

In addition to estimating unconstrained diploid selection coefficients *s*_1_ and *s*_2_, we also want to estimate selection coefficients in the one-parameter selection modes *additive, recessive, dominance*, and *heterozygote difference*. To this end, we introduce constraints on *s*_1_ and *s*_2_ using the framework of Lagrange multipliers (e.g. Hoffmann et al. 2010, Ch. 7.5). Denoting the likelihood by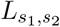, the optimization problem can be formulated as maximizing the conditional log-likelihood 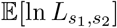 in equation (2) subject to *g*(*s*_1_, *s*_2_) = 0 for an arbitrary function *g*(·, ·), which can be solved by introducing the Lagrangian 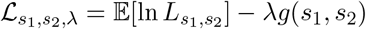 and solving 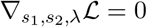.

All aforementioned one-parameter selection modes of interest can be expressed as a linear constraint, that is, *as*_1_ − *bs*_2_ = 0 for suitable *a, b* ∈ ℝ. In the Lagrange multiplier formalism, we can thus set *g*(*s*_1_, *s*_2_) = *as*_1_ − *bs*_2_ and solve the constrained optimization problem to obtain the MLE under the respective mode of selection. In Section S.2.3 of the Supplementary Material, we derive explicit expressions for the update rules for *s*_1_ and *s*_2_ for arbitrary *a, b*. As an example, Figure 2B shows the iterations of the *additive, recessive, dominance*, and unconstrained general diploid selection EM-HMM algorithms on a dataset simulated under *additive* selection.

#### 2.2.3 Estimation of the initial allele frequency distribution

The discretized distribution of the initial allele frequency can be fixed for the analysis or estimated as well. When estimating it, we fit a beta distribution to the initial frequency, as it is a flexible distribution with only two parameters, denoted by (*α, β*), which avoids potential over-parametrization. If we assume that the initial distribution does not depend on the selection coefficients, then the discretized version of the EM update rule given in equation (2) becomes

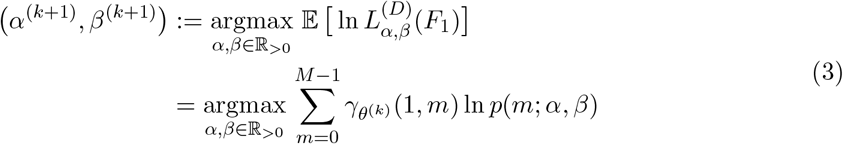

for the parameters (*α, β*). Here, 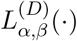 denotes the likelihood of a discretized beta distribution, the conditional expectation 𝔼 [·] is now parameterized by 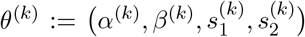, and *p*(*m*; *α, β*)is the integral of a beta distribution with parameters (*α, β*) over the *m*-th discretization interval. At each iteration *k* of the EM-HMM algorithm, we solve equation (3) numerically to update *α* and *β* alongside the selection coefficients. Since the EM update for the selection coefficients already requires computing 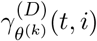, the extra step of updating the initial condition comes at minimal computational cost. We observed that estimating the initial distribution does affect the accuracy of the selection coefficient estimation, see Figure S.10 in the Supplementary Material, while providing more flexibility.

#### 2.2.4 Additional implementation details

Analyzing data using our EM-HMM algorithm requires choosing the number of hidden states or discretization intervals *M*, and where to place the discretization points 𝒢:= {*g*_0_ = 0, *g*_1_, …, *g*_*M*−1_ = 1}. A common choice for the discretization points is to space them equidistantly, that is 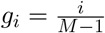. However, we observed that this discretization did not perform well when selection was strong, see Section S.5 in the Supplementary Material for details. We thus decided to use the Chebychev nodes 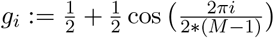 for 0 *≤ i ≤ M* − 1. These nodes are often used for numerical integration, since they mitigate Runge’s phenomenon (Mathews and Fink 2003, Ch. 4.5). Intuitively, when using equidistant spacing, the variance of the normal distribution for the transition used in the M-step becomes smaller than the size of the discretization intervals near the boundary, and consequently, the density at the discretization points does not approximate the probability mass in the interval well. Using the Chebychev nodes effectively increases the density of discretization points near the boundary and decreases it in the interior, mitigating this problem.

We analyzed simulated data using different choices of the number of hidden states *M* in Section 3.1.5. We observed that the inference does not perform well when using fewer than 250 hidden states, but is stable for higher values. To balance accuracy of the EM-HMM algorithm and computational efficiency, we thus choose *M* = 500 hidden states in all analyses. Unless stated otherwise, we initialize the selection coefficients for the EM-HMM algorithm with 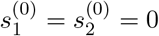 and set the starting parameters for the initial distribution to *α*^(0)^ = *β*^(0)^ = 1, which corresponds to the uniform distribution. For all analyses, we use the convergence criterion that the difference in log-likelihood between iteration *k* and iteration *k* + 1 must be less than 10^−3^. Additionally, since the EM algorithm can require many iterations to meet this convergence criterion, we tested accelerating the EM using the SQUAREM procedure (Varadhan and Roland 2008). We found that, while this approach reduced the total number of iterations required, the total runtime of the algorithm increased, because of the additional computational cost per iteration. Since we observed no noticeable increase in accuracy of the estimation, we thus proceeded with the regular EM approach.

We noticed that, especially for simulations under neutrality, the EM-HMM for many replicates would return 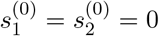 as the MLE, but the actual MLE would deviate slightly from 0, with a log-likelihood increase *<* 10^−3^. Nonetheless, this would result in many replicates reporting a log-likelihood ratio of exactly 1, and consequently, p-values around 1 were not well calibrated. To avoid potentially unwanted distortions to the distribution of p-values, we required the EM-HMM to perform at least 5 iterations before stopping, taking the maximum log-likelihood obtained in the first 5 iterations if further iterations do not increase the log-likelihood. In practice, this has minimal effect on replicates simulated under selection, since the EM-HMM requires more than 5 steps to converge in most of these cases.

### 2.3 Distinguishing between modes of selection

#### 2.3.1 P-values for a single alternative

Before describing our approach to the full problem of inferring the mode of selection with multiple alternatives, we first outline the solution to the task of rejecting neutrality in the case of a one-parameter alternative mode of selection. For a given replicate and one-parameter selection mode, we use the EM-HMM algorithm to compute the MLEs 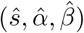, as well the log-likelihood *ll*_*s*_ for these parameters. Moreover, we compute the MLEs 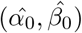 and log-likelihood *ll*_0_ under neutrality (*s* = 0). Treating the parameters of the initial distribution as nuisance parameters, we then perform standard likelihood-ratio testing using the likelihood-ratio statistic *D* = 2(*ll*_*s*_ − *ll*_0_). As the sample size goes to infinity, Wilks’ theorem (Wilks 1938) implies that *D* should be *χ*^2^ distributed with one degree of freedom under the null hypothesis *s* = 0, which we denote by *χ*^2^(1). However, each replicate consists only of one set of temporal samples, so we are not operating in the asymptotic regime of Wilks’ theorem, and have no theoretical guarantee regarding the distribution of *D*. Nonetheless, in Section 3.1.3, we show that if we compute p-values using *p* = 1 − ℙ {*χ*^2^(1) *< D*}, assuming a *χ*^2^(1) distribution, then the resulting p-values are well-calibrated, and thus the asymptotic formula provides a good approximation. P-values computed this way can thus be used accept or reject the null hypothesis of neutrality for a given dataset, under the given one-parameter alternative.

#### 2.3.2 Multiple alternatives

The mode of selection underlying a given dataset might not be known a priori, so we devised the following strategy that aims at inferring the mode of selection among multiple alternatives from the given data. We found that unconstrained estimation of the MLE 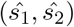 in the full two-dimensional parameter space has higher variance than inference in the bespoke constrained modes. Thus, our approach to infer the mode of selection relies solely on the constrained modes of selection. Motivated by the fact that the test statistic *D* is well-calibrated in the case of single alternative tests for all modes of selection, we propose the following procedure to establish significance and classify significant replicates:

1. For all replicates in the target dataset, obtain the MLE parameters for all constrained modes of selection: *Additive, dominance, recessive*, and *heterozygote difference*.
2. For each replicate, compute the test statistic

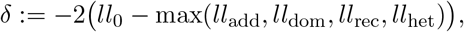

where *ll*_0_ is the likelihood under neutrality, and *ll*_*m*_ is the maximal likelihood for mode *m*.
3. For each replicate, compute its p-value as *p* = 1 − ℙ{*χ*^2^(1) *< δ*}, and reject neutrality at the desired level of significance.
4. If neutrality is rejected, classify the mode of selection according to

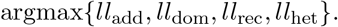

In the case that the *heterozygote difference* mode is the most likely mode, classify as *over-dominant* if *s*_1_ *>* 0 and as *underdominant* if *s*_1_ *<* 0.

We found that the p-values computed from the statistic *δ* using a *χ*^2^(1) distribution with 1 degree of freedom are not as well calibrated as as the p-values in the single alternative tests, and can lead to slightly anti-conservative p-values (see Section 3.1.3) in the tail of the distribution. This is not unexpected, as some p-values from the well-calibrated single-alternative *D* statistic are replaced by higher-likelihood estimates from additional modes of selection. However, using a *χ*^2^(2) distribution resulted in an even poorer fit.

Additionally, we explored fitting parametrized distributions to parametric bootstrap simulations of the statistic *δ*. In Figure S.14 in the Supplemental Material, we compare p-values computed using the *χ*^2^(1) and *χ*^2^(2) distribution to p-values computed using parameterized distributions, such as the generalized gamma distribution, that were fit to bootstrap simulations with 10,000 replicates in a scenario related to our data analysis in Section 3.2. While the bootstrap procedure achieved better calibration in this scenario, the procedure required extensive simulations specific to a given scenario to determine the requisite parameters. We thus recommend using the *χ*^2^(1) distribution, since it is a fast and flexible procedure that leads to reasonably well calibrated p-values, even in the tail of the distribution, and is partly motivated by theory. In practice, we do recommend simulating data in the specific scenario of interest to confirm that the p-values are well calibrated.

## 3 Results

### 3.1 Simulation Study

To assess the performance of our EM-HMM algorithm to estimate selection coefficients and infer the correct mode of selection across a variety of selection regimes, we simulated datasets under several combinations of parameters and exhibit the accuracy of the inference.

#### 3.1.1 Simulation parameters

We simulated population allele frequency trajectories under the discrete Wright-Fisher model for a given number of generations and sampled a certain number of haplotypes at given times from a binomial distribution with success probability given by the respective population allele frequency. Specifically, we simulated datasets under neutrality (*s*_1_ = *s*_2_ = 0), each one-parameter selection mode (*additive, recessive, dominant*), where we set *s* = *s*_2_, and complete *over-* or *underdominance* with *s* = *s*_1_ and *s*_2_ = 0. We simulated using selection coefficients *s* ∈ {0.005, 0.01, 0.025, 0.05} and the length of the simulated trajectory in generations *T* ∈ {101, 251, 1001}. We considered two different types of initial conditions: 1) The population allele frequency is initialized at a fixed value *p* ∈ {0.005, 0.25} for each replicate, and 2) the initial frequency of each replicate is drawn from the set 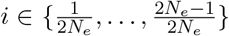 with probability proportional to 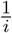. The latter is the stationary distribution for neutral segregating sites under the Poisson Random Field model (Sawyer and Hartl 1992), and thus corresponds to selection from standing variation. We conditioned the simulated data on the focal allele *A* not being lost or fixed in the samples, depending on the mode of selection. Specifically, for *additive, dominant*, and *recessive* selection, we conditioned on *A* not being fixed at the first generation and not being lost at the last generation; for *overdominance*, the simulation is conditioned on *A* segregating in the last generation; for *underdominance*, we conditioned on *A* segregating in the first generation.

For all simulations, the population size was set to *N*_*e*_ = 10,000. the sampling scheme consisted of *K* = 11 equidistantly-spaced points in time, where the first and last point align with the start and the end of the simulated trajectory, respectively. At each time, we sampled 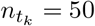 haploids. For each combination of parameters, we simulated 10,000 replicates under neutrality, and 1,000 replicates under each mode of selection and strength of selection. For each replicate, we performed inference using the same *N*_*e*_ that was used for the simulations under the four bespoke selection modes: *additive, recessive, dominant*, and *heterozygote difference*.

In Section 3.1.2, Section 3.1.3, and Section 3.1.4, we present results for the simulations with 251 generations and initial allele frequency fixed to *p* = 0.25. The results for other combinations of number of generations and initial allele frequency distribution can be found in Section S.3 of the Supplemental Material. In general, the results are similar, with the exception of extreme cases: For *recessive* or *underdominance* with low initial frequencies, the selected alleles is often lost, resulting in inaccurate estimates.

#### 3.1.2 Estimating selection coefficients

In Figure 3A, we present the distribution of the estimated selection coefficient *ŝ* across replicates simulated under the *additive, recessive, dominant, overdominant*, and *underdominant* mode. Estimates of the selection coefficients are in general accurate and unbiased. For intermediate-to-strong selection, *overdominance* has the highest variance in *ŝ* estimates. The likely explanation is that the *overdominance* replicates reach stationarity (*p* = 0.5) quickly, resulting in fewer informative samples, and the variance of the estimates is increased. In Figure S.9 of the Supplementary Material, we plot the selection coefficients estimated using the unconstrained EM for all modes of selection, that is, estimating both *s*_1_ and *s*_2_. *Additive, recessive, dominant*, and *overdominant* selection are well-estimated by the unconstrained EM, whereas the estimates for *underdominance* show more variability. We thus recommend exploring the bespoke one-dimensional modes first.

**Figure 3:**
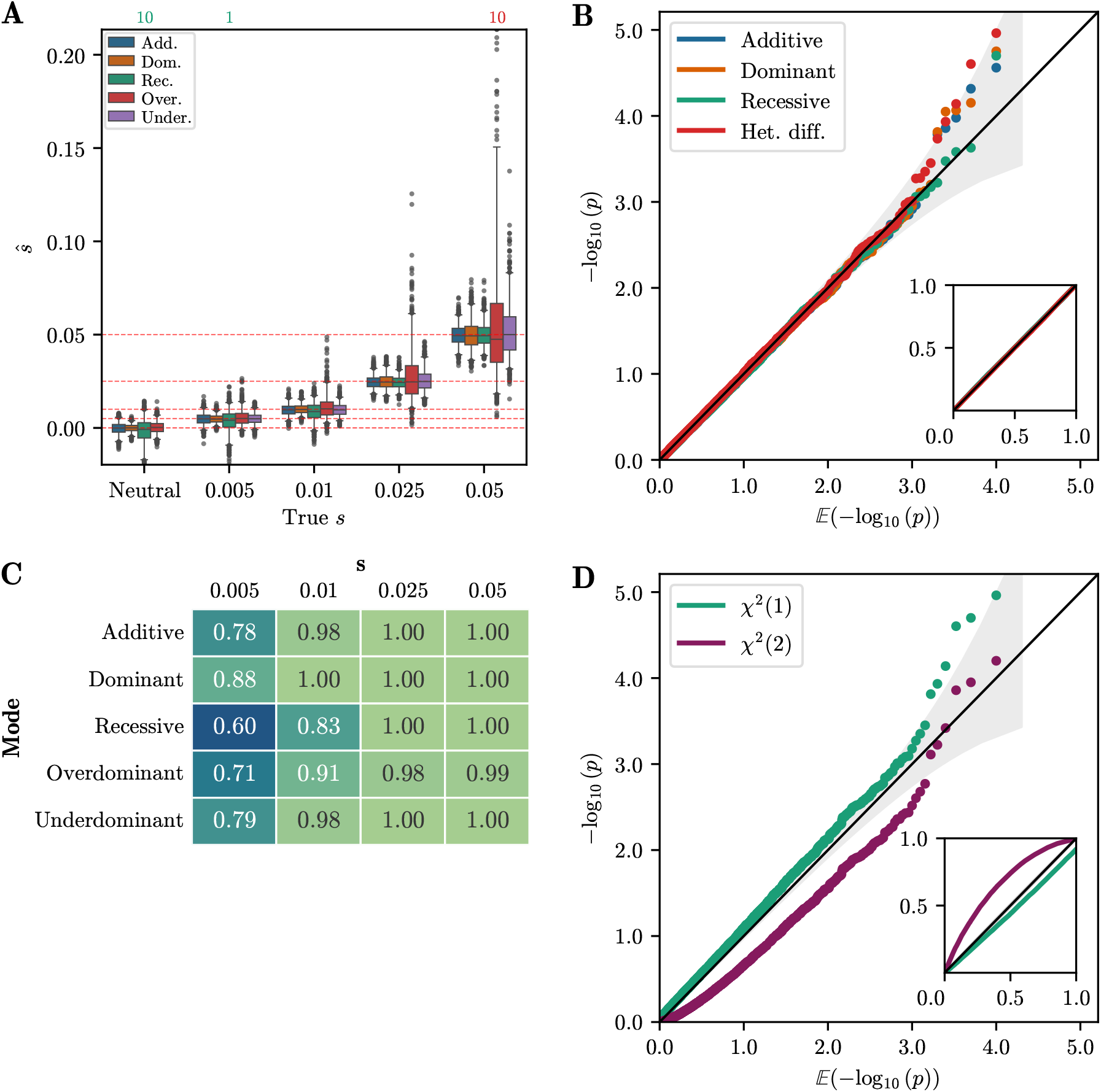
A) Boxplot of *ŝ* for 1,000 replicates simulated under each selection mode. Whiskers extend to 2.5% and 97.5%-tiles. Number of estimates outside plotting range indicated above the plot. Estimates are generally unbiased and have low variance, with the exception of *overdominance*. B) Q-Q plot of −log_10_(p) against 𝔼 [−log_10_(p)] of single-alternative tests for neutral replicates. Inset shows same plot for raw values. The p-values are well-calibrated under all modes of selection. C) Table of AUC values based on likelihood-ratios for each selection mode and selection strength simulated. For *s >* 0.01, AUC values are near 1, indicating perfect discrimination between neutral and non-neutral replicates. D) Q-Q plot of −log_10_(p) against 𝔼 [−log_10_(p)] for the *δ* statistic under the *χ*^2^(1) and *χ*^2^(2) distributions. The *χ*^2^(1) distribution is better calibrated, although it is slightly anti-conservative in the tail. For all simulations, the number of generations is *T* = 251 and the initial condition is fixed at frequency *p* = 0.25.

#### 3.1.3 Validation of single-alternative p-values

Next, we report the single-alternative p-values obtained using the *χ*^2^(1) distribution, as described in Section 2.3.1, for all four modes: *additive, recessive, dominant*, and *heterozygote difference*. Figure 3B shows a Q-Q plot, where we plot the empirical p-values against their expected value for replicates simulated under neutrality. For all selection modes, the points follow the line *y* = *x* closely, indicating that the p-values are well-calibrated, and that the distribution of the likelihoodratio statistic *D* is well-approximated by a *χ*^2^(1) distribution.

Additionally, we report the performance of the single-alternative tests in terms of the area under the curve (AUC) of a receiver-operator characteristic curve. An AUC value of 0.5 indicates no power to distinguish neutral replicates from non-neutral replicates, whereas an AUC value of 1 indicates perfect discrimination. Figure 3C reports the AUC values for each simulated selection mode over the range of selection parameters *s*. We observe that in these scenarios, the method is well-powered to reject the null hypothesis for *s* ≥ 0.01. AUC values are lower in the case of *recessive* selection, since 251 generations can be insufficient for replicates at low frequency to escape the drift-dominated regime.

#### 3.1.4 Validation of selection mode inference

We furthermore computed p-values for the simulated datasets in the case of multiple alternatives, following the procedure outlined in Section 2.3.2. A Q-Q plot of the p-values against their expected values is shown in Figure 3D. In this scenario, the *χ*^2^(1) distribution provides a good fit for the *δ* statistic, although it is slightly anti-conservative in the tail of the distribution. However, in other scenarios (see Section S.3 in the Supplementary Material), the *χ*^2^(1) distribution can be slightly *over*conservative. Since the different alternative selection modes are all embedded in a two-dimensional parameter space, Figure 3D also shows p-values computed from the *δ* statistic using a *χ*^2^(2) distribution. However, we observe that these p-values are poorly calibrated, and thus recommend using the *χ*^2^(1) distribution.

We then tested the ability of our approach to correctly identify the mode of selection. Figure 4 shows a confusion matrix resulting from inferring the mode of selection with a p-value significance threshold of 0.05. Each row of the confusion matrix represents all replicates simulated under a particular mode of selection. The numbers in the corresponding column indicate the fraction of replicates where a particular mode is inferred. Values on the diagonal reflect identification of the correct mode, whereas off-diagonal values reflect replicates where the mode is not correctly inferred. For *s* = 0, neutrality is rejected for 6.3% of replicates, indicating that the *χ*^2^(1) distribution is anticonservative. For *s* = 0.05, the correct model is inferred for a majority of replicates for all modes of selection. For weaker selection, *s* = 0.01, only *dominant* and *overdominant* selection are inferred for a majority of replicates. However, neutrality is rejected for over 50% of the replicates for all modes of selection, indicating that power to reject neutrality exists, but accuracy to correctly infer the mode of selection is more limited.

**Figure 4:**
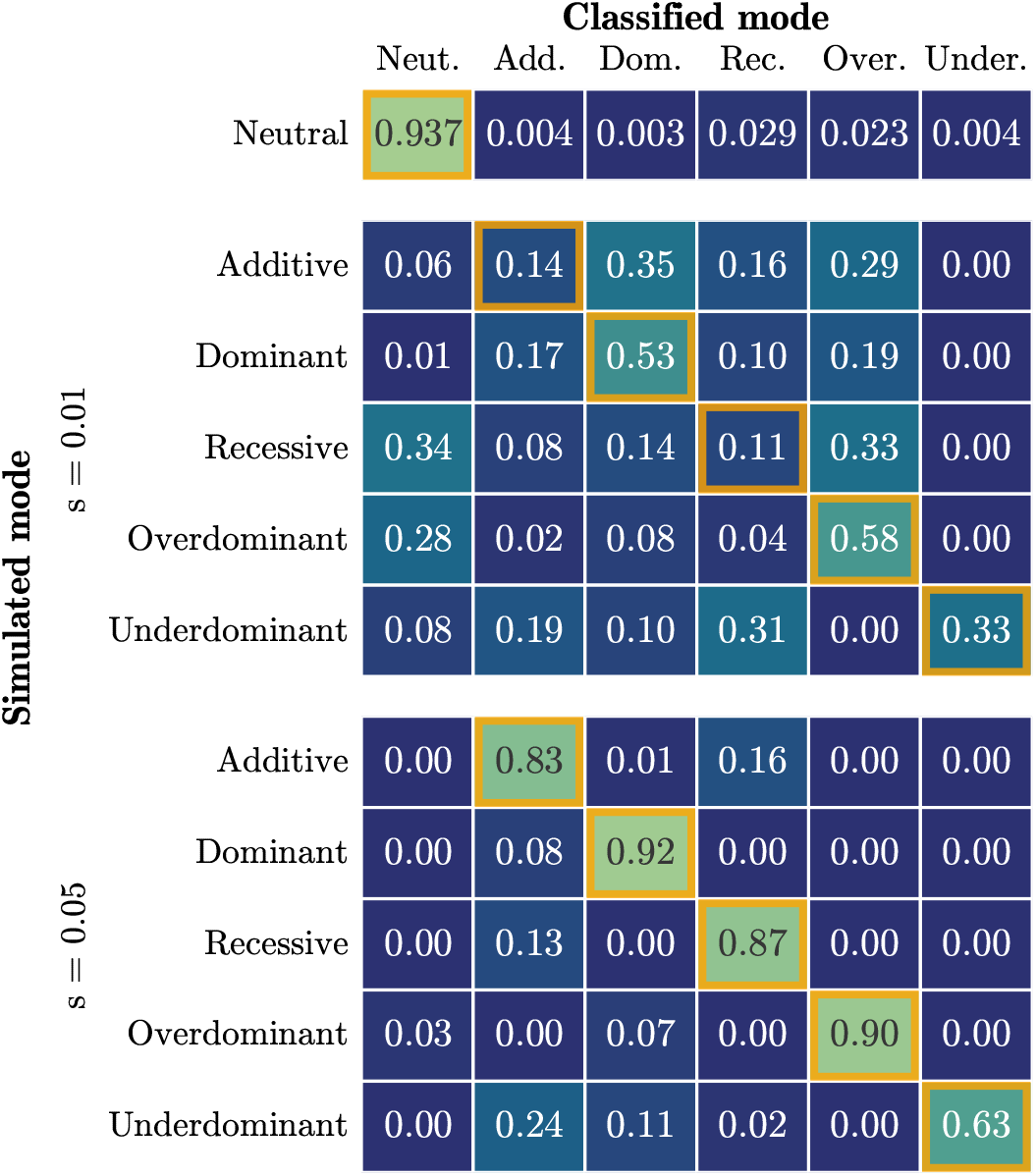
Confusion matrix of the results when inferring the mode of selection, with a p-value threshold of 0.05 to reject neutrality. Cells with an orange border represent correct classification. The correct model is inferred for the majority of replicates for *s* = 0.05 for all modes of selection. For all simulations, the number of generations is *T* = 251 and the initial condition is fixed at frequency *p* = 0.25.

Lastly, we investigate accuracy of the MLE of the selection parameters, given that the mode was correctly classified. Figure 5 shows the distribution of *ŝ* estimates for replicates that were correctly classified as *additive, dominant*, or *recessive* selection, as well as *ŝ* for neutral replicates that were classified *incorrectly*. For larger *s*, correctly classified replicates are more tightly clustered around the true value, with far fewer outliers compared to the distribution using all replicates. For lower values of *s* and neutrality, the *ŝ* estimates for replicates classified as non-neutral are biased. This is primarily due to a “winner’s curse” effect, where neutrality is only rejected for replicates with extreme MLEs.

**Figure 5:**
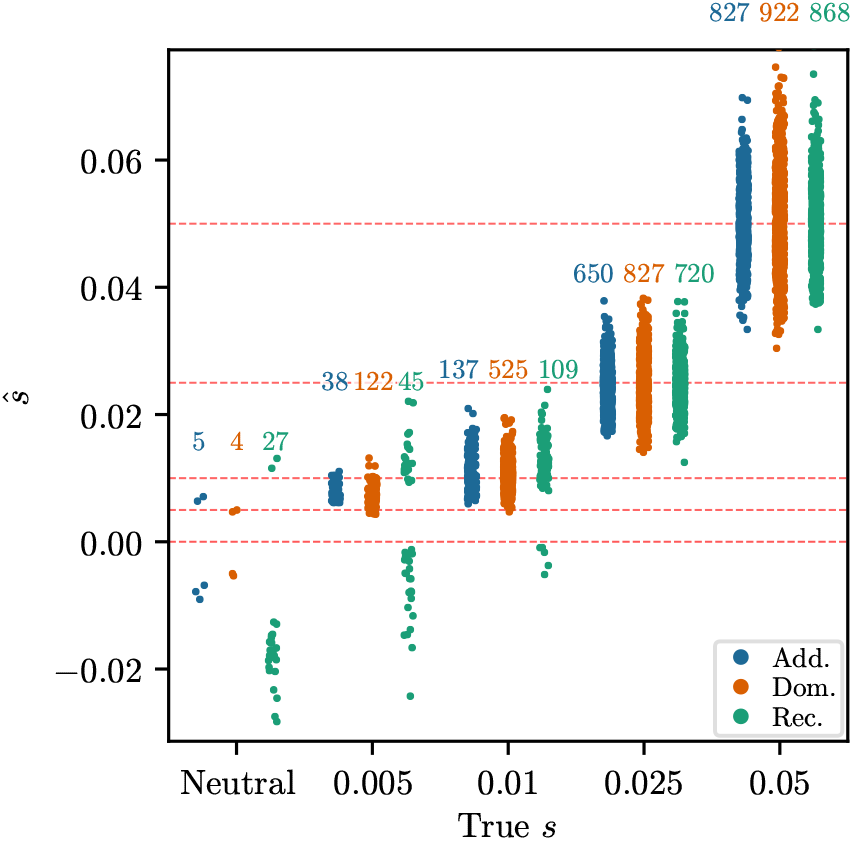
Strip plots of inferred against true *s*, conditioned on the correct model being chosen among multiple alternatives. In the neutral case, the plot shows the neutral replicates classified into certain modes. For neutrality, the strip plot is for 1,000 replicates randomly chosen from 10,000 simulated. The number over each strip indicates the number of replicates correctly classified out of the 1,000. For *s* = 0.005 and neutrality, the inferred values are strongly biased, due to a “winner’s curse” effect. For all simulations, the number of generations is *T* = 251 and the initial condition is fixed at frequency *p* = 0.25.

#### 3.1.5 Robustness of selection coefficient estimation

Next, we investigated how estimation accuracy is affected if we vary parameters that were fixed in the previous simulation study. We varied the number of times a sample is taken *K* ∈ {2, 5, 11, 35, 101}, the number of haploid samples at each timepoint 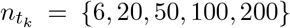, the effective population size *N*_*e*_ ∈ {10^2^, 10^3^, 10^4^, 10^5^, 10^6^}, and the number of hidden states in the HMM *M* ∈ {100, 250, 500, 1,000, 2,000}. We varied one parameter at a time, and fixed the others to 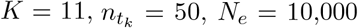, and *M* = 500. We fixed the selection coefficient *s* to 0.025 for all simulations, set the total number of generations to *T* = 251, and set the initial distribution to a fixed frequency at *p* = 0.25. Boxplots of the estimated selection coefficients for *additive, recessive*, and *dominant* selection are shown for each scenario in Figure 6. Similar to Mathieson and McVean (2013) in the *additive* case, we find that the accuracy does not depend strongly on the sampling scheme except for very low values, such as sampling 6 haplotypes per timepoint or only sampling at the beginning and end of the trajectory. However, the boxplots are tighter for larger sample sizes and more sampling times. This implies that both variance due to finite sampling and variance in the true allele frequency contribute to uncertainty of the selection coefficients estimates. Similarly, the accuracy of the estimates are comparable for 250, 500, 1,000, and 2,000 hidden states. However, for 100 hidden states, the estimates are strongly biased downwards. We thus recommend using *M* = 500 to balance accuracy with computational efficiency.

**Figure 6:**
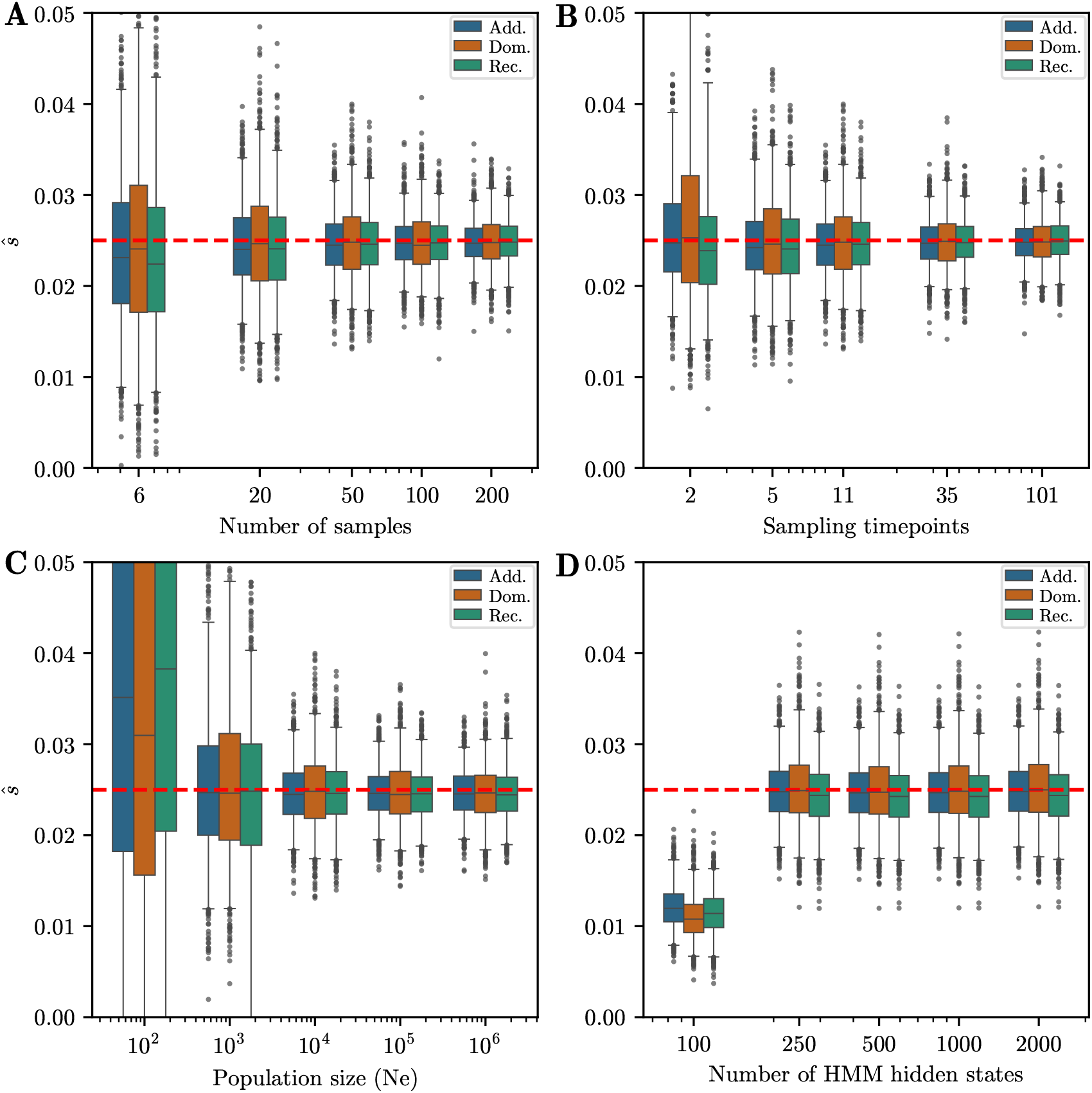
Boxplots for data simulated and inferred under *additive, dominant*, and *recessive* selection for different parameter combinations. Whiskers extend to 2.5% and 97.5%-tiles. We vary: A) Number of samples taken at each timepoint, B) Number of timepoints sampled, C) Effective population size, D) Number of hidden states in the HMM. Except for the lowest parameters, estimates of the selection coefficient are unbiased. Width of boxplots decreases as the number of samples and sampling timepoints increases. For all simulations, the number of generations is *T* = 251 and the initial condition is fixed at frequency *p* = 0.25. When not varied, parameters used are *K* = 11 sampling times, 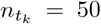 samples at each timepoint, *N*_*e*_ = 10,000, and *M* = 500 hidden states.

To investigate the accuracy of the estimates when the mode of selection is incorrect, we reanalyzed a subset of the data under scenarios in which we misspecify the mode. We consider two cases: (1) Simulating under *additive, dominant, recessive, over-*, or *underdominant* selection and analyzing under *additive* selection, and (2) simulating under *additive* selection and analyzing under *additive, dominant, recessive*, or *heterozygote difference*. Figures 7A and 7B depict boxplots of the MLEs *ŝ* under both of these misspecification scenarios.

**Figure 7:**
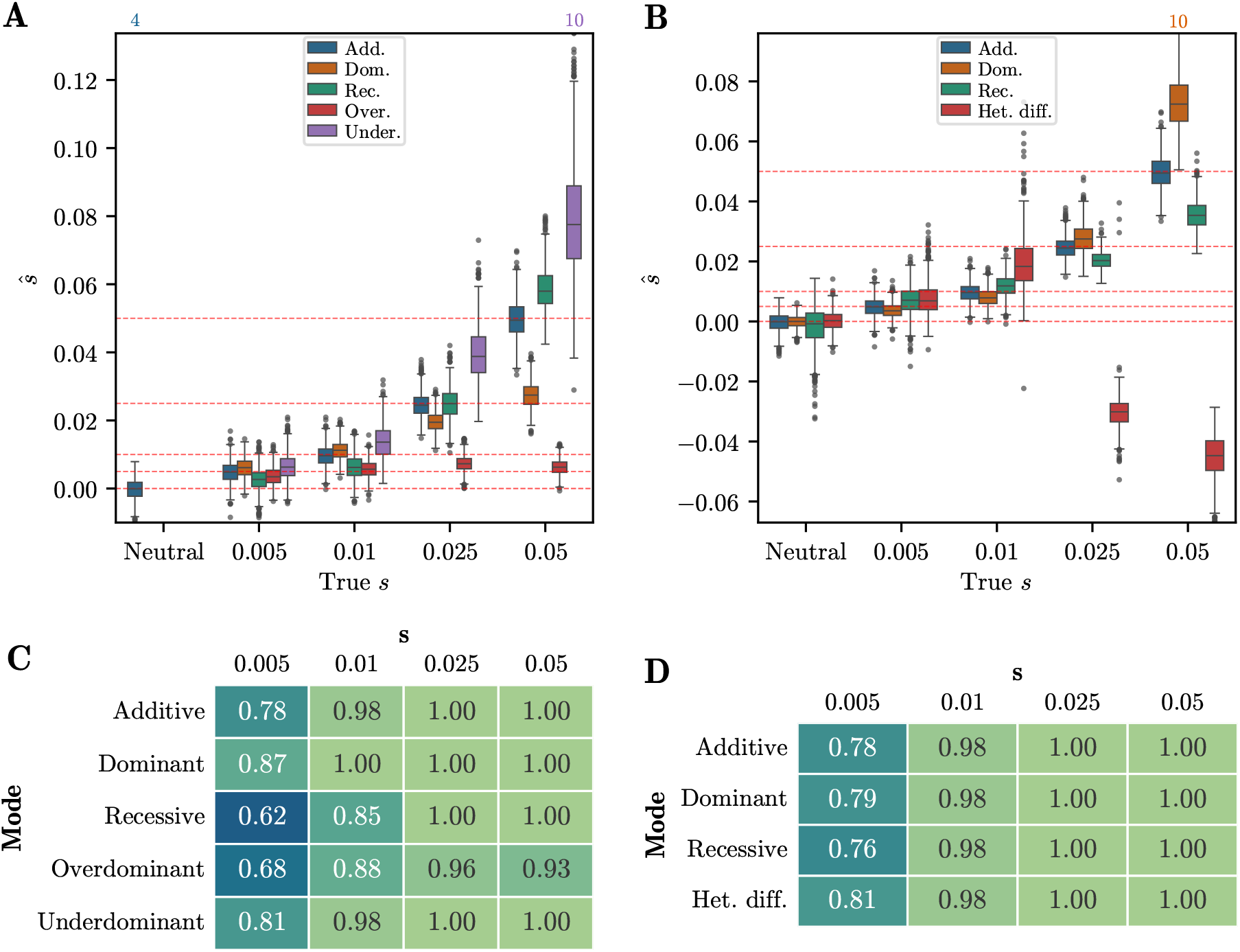
A) Boxplots of *ŝ* for 1,000 replicates simulated under each selection mode and analyzed using *additive* selection. The estimates are mostly inaccurate. B) Boxplots of *ŝ* for 1,000 replicates simulated under *additive* selection and analyzed under each selection mode. Again, the estimates are not accurate if the mode is incorrect. For both sets of boxplots, whiskers extend to 2.5% and 97.5%-tiles. C) Table of AUC values for data simulated under each selection mode using likelihood-ratios obtained assuming *additive* selection. Values are lower compared to analyzing under the correct mode of selection, but still show substantial power to reject neutrality. D) Table of AUC values for data simulated under *additive* selection using likelihood-ratios obtained assuming each possible selection mode. The values are very similar across selection modes. For all simulations, the number of generations is *T* = 251 and the initial condition is fixed at frequency *p* = 0.25.

When simulating under each mode of selection and analyzing under the *additive* EM-HMM, the estimates of *ŝ* are biased for all modes, most strongly for *overdominant* selection at high *s*. Alleles undergoing strong *overdominant* selection quickly reach a stationary frequency of *p* = 0.5. Since the alleles do not continue to increase in frequency, the *additive* EM-HMM thus underestimates the true selection strength. In the case when we simulate under *additive* selection and analyze under each selection mode, the *ŝ* estimates are not strongly biased for *recessive* and *dominant*, but reverse sign for *heterozygote difference* at high selection strengths. In this case, the allele frequency increases beyond *p* = 0.5, so the *heterozygote difference* EM-HMM changes from *overdominance* to *underdominant* selection. Overall, the biases in the estimates *ŝ* indicate that using the incorrect one-parameter EM-HMM provides inaccurate estimates of *s*.

We furthermore investigated the power to reject or accept neutrality in the single-alternative testing framework described in Section 2.3.1 under each of these misspecification scenarios. The resulting AUCs are presented in Figures 7C and 7D. For the case of analyzing other modes under *additive* selection, AUC values are slightly lower than the values in Figure 3C, but still substantially greater than 0.5. For the *additive* datasets analyzed using the incorrect selection modes, the AUC values do not differ substantially between different modes. In both misspecification scenarios, the one-parameter *additive* EM-HMM has sufficient power to accurately reject or fail to reject neutrality for moderate-to-high selection coefficients. Thus, if the goal is only to identify non-neutral evolution, it can be sufficient to analyze given data using the *additive* EM-HMM only, but for accurate characterization of the selection coefficients, the correct mode needs to be used.

#### 3.1.6 Inferring effective population size

We next explored applying our method to infer the effective population size *N*_*e*_ of the underlying population. The approach to derive the update rules for the EM-HMM algorithm provided in equation (2) and equation (3) does not readily yield an update rule for *N*_*e*_. Thus, we instead use a grid-based approach to estimate it. Specifically, we compute the likelihood of observing a given replicate under the neutral HMM with *s*_1_ = *s*_2_ = 0 and a given population size *N*_*e*_. To combine power across replicates or loci, we compute the sum of the log-likelihoods over all replicates on a grid of *N*_*e*_ values, then interpolate between these grid values and use the value of *N*_*e*_ that maximizes this composite likelihood surface as our estimate of *N*_*e*_.

We noticed that when simulating data with large values of *N*_*e*_, the resulting likelihood surface would often be very flat, making estimation challenging. To counteract this, we condition our likelihoods on observing at least one polymorphic sample at any timepoint, by dividing the likelihood by 1 −ℙ {no focal alleles observed} −ℙ {no non-focal alleles observed}. This penalizes high values of *N*_*e*_ and results in more peaked likelihood surfaces. Figure S.17 of the Supplementary Material shows several example composite likelihood surfaces. Even after this conditioning procedure, the surfaces are still fairly flat, but they do allow determining a clear maximum. Furthermore, we found that the *N*_*e*_ estimates were most accurate when the initial condition for computing the likelihood is fixed as the uniform distribution, rather than estimated as in Section 2.2.3. A possible explanation could be that the procedure to estimate the initial condition is affected by the choice of *N*_*e*_, which in turn could bias the estimates.

We simulated 25 batches of 10,000 neutral replicates under the scenario with 251 generations, *p* = 0.25 initial condition, and the sampling scheme used in Section 3.1 for *N*_*e*_ ∈ {2,500, 10,000, 40,000}. We estimated *N*_*e*_ for each batch using the above procedure. Figure 8 shows boxplots of the inferred *N*_*e*_ for each batch. The *N*_*e*_ values estimated from the grid-based procedure are tightly clustered around the true value, although slightly biased upwards in all cases. More powerful approaches to estimate *N*_*e*_ exist, for example, using IBD segments in contemporary data (Browning and Browning 2015). However, we believe that when analyzing time series genetic data, the performance of the grid-based HMM procedure is acceptable and yields the most appropriate estimate of *N*_*e*_ to use in downstream analyses.

**Figure 8:**
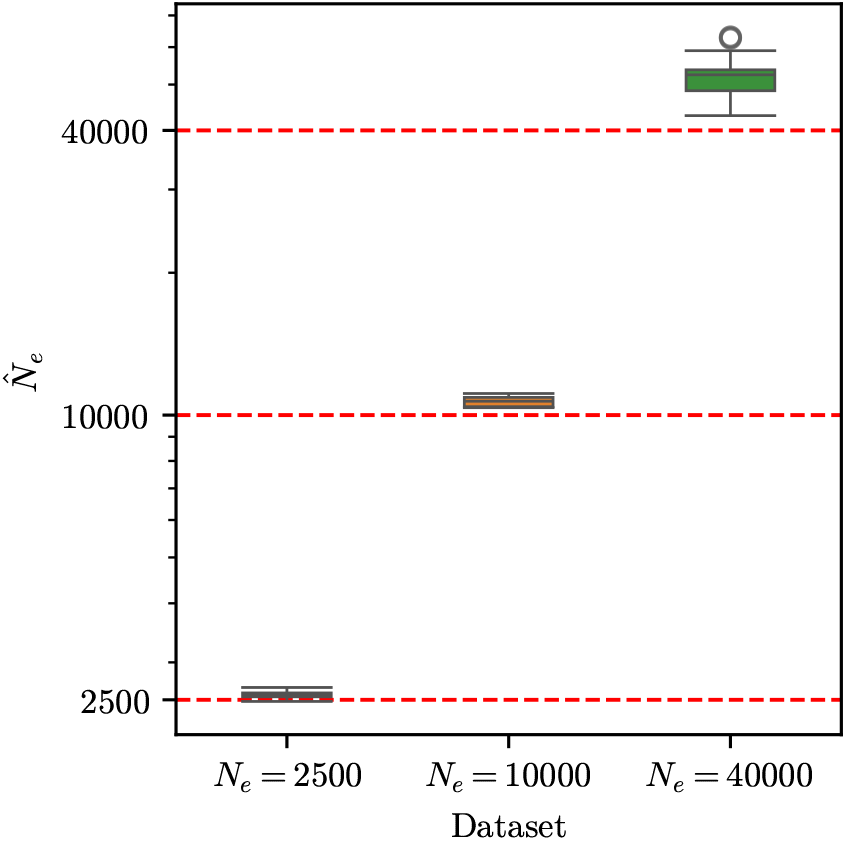
Boxplots of *N*_*e*_ estimated using the grid-based HMM procedure, for data simulated under neutrality for *N*_*e*_ ∈ {2,500, 10,000, 40,000}. For each simulated value *N*_*e*_, each of the 25 estimates shown are based on the composite likelihood of a batch of 10,000 replicates. *N*_*e*_ estimates are slightly biased upwards with low variance. For all simulations, the number of generations is *T* = 251 and the initial condition is fixed at frequency *p* = 0.25. Whiskers extend to 2.5% and 97.5%-tiles.

### 3.2 Ancient DNA dataset from Great Britain

#### 3.2.1 Description of GB aDNA dataset

Having demonstrated the ability of our approach to accurately characterize selection in simulated data, we applied our method to time series genetic data obtained from human ancient DNA. To this end, we extracted the genotype information from a subset of the individuals in the Allen Ancient DNA Resource (AADR), version v54.1, which is a frequently updated repository that aims at comprising most currently published ancient DNA datasets (Mallick et al. 2024).

Our method assumes that the data originates from a single panmictic population. We thus follow a rationale similar to Mathieson and Terhorst (2022) and restrict our analyses to samples from Great Britain in the last 4,450 years. We manually removed samples that were not from the mainland. Restricting to this geographic area and time window, it is unlikely that the data we analyze substantially violates the assumption of a single panmictic population, since the last major admixture event into Great Britain is estimated to have occurred before 4,450 years ago (Olalde et al. 2018; Chintalapati et al. 2022), although evidence for some more recent gene flow has been presented (Patterson et al. 2022; Gretzinger et al. 2022). Figure S.18 in the Supplementary Material shows a map with sample locations and times. Moreover, Figure S.19 in the Supplementary Material depicts two PCA plots, demonstrating that the samples cluster with modern European individuals on a global scale, but do not exhibit evidence of strong structure on a local scale.

When analyzing samples that were genotyped using the 1240K capture assay and samples genotyped using whole genome sequencing together, we noticed spurious signals of selection, see Figure S.22 in the Supplementary Material. As a conservative approach, we thus only analyzed samples that were genotyped using 1240K capture, and excluded the 174 whole genome samples in this geographic area and timeframe from our dataset. We note that this conservative approach also removed the present-day samples. Our resulting dataset, henceforth referred to as the GB aDNA dataset, thus comprises 504 ancient pseudo-haploid samples genotyped using the 1240K assay and spanning 125 generations, when using a generation time of 28.1 (Moorjani et al. 2016). Samples in the same generation are binned together to form the final dataset used for analysis. These individuals are a subset of the individuals published by Olalde et al. (2018), Patterson et al. (2022), and Gretzinger et al. (2022).

We furthermore applied three filters to each SNP in the dataset. First, each SNP must have genotyped samples at two or more timepoints. Second, each SNP must have more than 50 (10% of 504) samples genotyped in total. Third, the minor allele frequency when pooling all samples at a given SNP across time must be greater than 0.05. We expect only SNPs that pass these filters to yield reliable signals of selection. In total, out of the 1,150,638 SNPs available, 743,417 (64.6%) pass these three filters and were used in the final analysis.

#### 3.2.2 Data-matched simulations

To assess the accuracy of our method in the specific context of the GB aDNA dataset, we simulated two datasets matching the sampling scheme and timeframe of the GB aDNA dataset: In the first, which we refer to as the IBDNe dataset, we simulated the data using a history of effective population sizes that vary over time, using a population size history for the British population previously inferred from the UK10K dataset (Browning and Browning 2015). In the second, which we refer to as the const-Ne dataset, we simulate under a single constant *N*_*e*_ estimated from the GB aDNA dataset (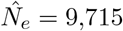, see Section 3.2.3). We show results for the IBDNe dataset in this section and show results for the dataset with constant *N*_*e*_ in Section S.6 of the Supplementary material.

We simulated allele frequency trajectories under a particular mode of selection for *T* = 125 generations using selection coefficients *s* ∈ {0, 0.005, 0.01, 0.025, 0.05}, and sampled haplotypes given each trajectory according to the sampling times and sizes of the GB aDNA dataset. For the IBDNe dataset, we used the graphreader tool (https://www.graphreader.com) to extract values of *N*_*e*_ at each generation from Figure 4A given by Browning and Browning (2015) for the period spanned by the samples in the GB aDNA dataset. These values were then used as time-varying *N*_*e*_ in the Wright–Fisher simulations. Figure S.15 of the Supplementary Material shows the extracted *N*_*e*_ values. In addition, we randomly omitted sampled haplotypes with probability equal to the fraction of missingness at a randomly selected SNP in the GB aDNA dataset, to emulate the same degree of missing data. We provide a histogram of the fraction of sampled haplotypes missing for each SNP in the GB aDNA dataset in Figure S.20 in the Supplementary Material. To further ensure that the simulated replicates match the GB aDNA dataset, we apply the same three SNP-based filtering criteria, and only keep replicates that pass all filters. We simulated 10,000 replicates under neutrality and for each *s* under each of the five one-parameter selection modes.

To generate the initial frequency for each simulated replicate, we first estimated the parameters *α* and *β* for the beta distribution of the initial frequency under neutrality (*s* = 0) at each SNP in the GB aDNA dataset that passes our filters, as described in Section 2.2.3. Figure S.21 in the Supplementary Material shows a histogram of the mean values *α/*(*α* + *β*) of the initial distributions estimated for each SNP. For each simulated replicate, we then chose one SNP uniformly at random, and set the initial frequency of the replicate equal to the mean of the chosen SNP. This procedure ensures that the initial frequency distribution of the simulated data matches the GB aDNA dataset closely, and captures any potential biases, for example, due to ascertainment of the 1240K SNP set.

To heuristically account for the variable population size history in the simulated data, we follow a strategy similar to Foll et al. (2014): We use the neutral replicates to estimate a shared constant 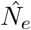 using the procedure described in Section 3.1.6, and use the inferred 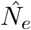 when estimating the selection coefficients for each replicate. We then computed the MLEs for the selection coefficients using our EM-HMM for each replicate, with the mode of selection in the analysis matched to the simulated mode, and show the distribution of the MLEs *ŝ* in Figure 9A. As for the simulated datasets in Section 3.1.1, the estimates of the selection coefficients are largely unbiased. However, unlike in the simulated datasets, for strong selection, the data-matched replicates have a slight downward bias. This is likely due to the fact that the data-matched simulation comprise half the number of generations as the simulation study presented in Figure 3A. The simulated datasets with *T* = 101 generations show a similar, though less pronounced, downward bias at high *s* – see Figure S.3A in the Supplementary Material. In addition, the variance in the estimate of *ŝ* for *underdominance* is lower than for the simulated datasets in Section 3.1.1 despite the shorter time horizon.

**Figure 9:**
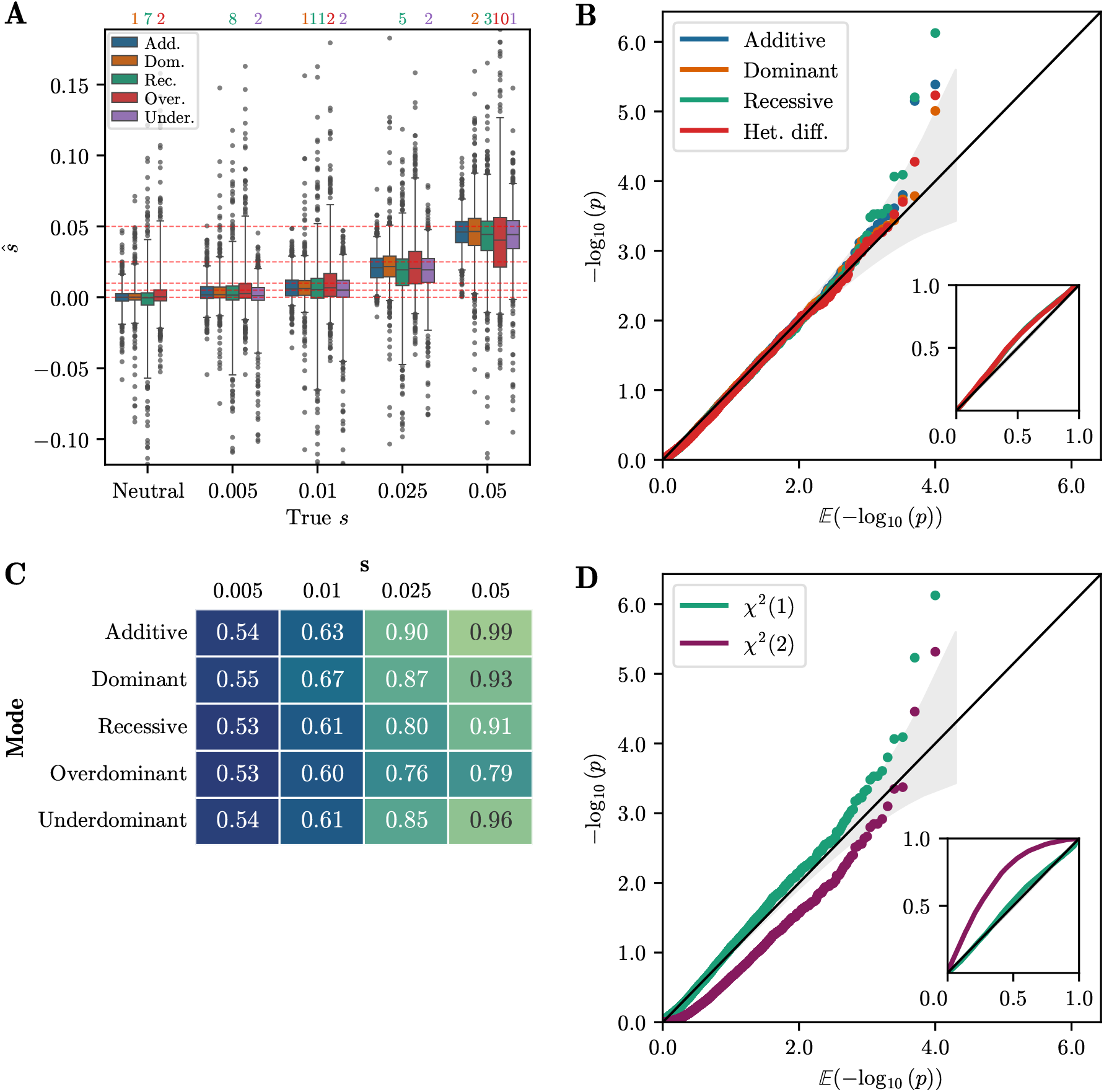
A) Boxplots of MLEs *ŝ* for all one-parameter selection modes. Each boxplot shows 1,000 random replicates of the 10,000 simulated. Whiskers extend to 2.5% and 97.5%-tiles. Estimates are largely unbiased for small *s*, and slightly biased downward for large *s*. B) Q-Q plot of −log_10_(p-value) against 𝔼 [−log_10_(p-value)] for single-alternative tests obtained using the *χ*^2^(1) distribution. Inset shows raw p-value against expected p-value. As for the simulated datasets, p-values are well calibrated. C) Table of AUC values for data-matched simulations using likelihood-ratios for each one-parameter selection mode. D) Q-Q plot of −log_10_(p-value) against 𝔼 [−log_10_(p-value)] for multiple-alternative likelihood ratio statistic *δ* using the *χ*^2^(1) and *χ*^2^(2) distributions. The *χ*^2^(1) distribution provides a good fit that is slightly anti-conservative in the tail. For all simulated replicates, the number of generations is *T* = 125, the value of *N*_*e*_ in each generation is derived from Browning and Browning (2015), and the initial frequencies match the GB aDNA dataset.

Next, we applied our procedure to test for a single alternative and our procedure to infer the mode of selection to the IBDNe simulations. For the test of a single alternative, Figure 9B shows a Q-Q plot of the p-values computed from the likelihood-ratio statistic *D* assuming a *χ*^2^(1) distribution against their expected value. We again observe that these p-values are well-calibrated. As in Section 3.1.3, we computed AUC values to assess power of the single-alternative tests on theIBDNe simulations, which are shown in Figure 9C. For *s* ≥ 0.025, the AUC values are between 0.8 and 1, with the exception of *overdominant* selection. In general, the AUC values are lower than in Figure 3C; for example, for *s* = 0.05 in the case of *overdominance*, the AUC is 0.81 for the IBDNe simulations, compared to 0.99 for the simulations in Section 3.1.1. This is likely due to the initial distribution that we used for the IBDNe matched simulations, which has more weight close to *p* = 0, combined with the reduced number of generations. Both of these properties result in smaller cumulative changes in allele frequency, and thus power to distinguish from neutral replicates is reduced.

We also applied our multiple alternatives framework introduced in Section 2.3.2. Figure 9D shows the p-values for the neutral replicates from the IBDNe dataset, computed using the statistic *δ* under *χ*^2^(1) and *χ*^2^(2) distributions, plotted against the expected value. The p-values are in general well-calibrated when using the *χ*^2^(1) distribution, although, as with the simulations in Section 3.1.4, the p-values are slightly anti-conservative in the tail of the distribution.

Figure 10 summarizes the inference of the selection mode for all replicates of the IBDNe simulations using the procedure described in Section 2.3.2; again using a p-value threshold of 0.05 to reject neutrality. For *s* = 0.05, we successfully reject neutrality for a large fraction of replicates, up to 95% of cases when simulating under *additive* selection, but the correct model is only inferred for 30% to 55% of the replicates. For the lower selection strength *s* = 0.01, neutrality is only rejected in a small percentage of the simulations. As in the simulation study presented in Section 3.1.4, we find that we can detect non-neutral evolution, but power to infer the correct mode of selection is limited. Note that we fail to reject 94% of neutral replicates, not 95%, due to the fact that p-values are slightly anti-conservative under the *χ*^2^(1) distribution.

**Figure 10:**
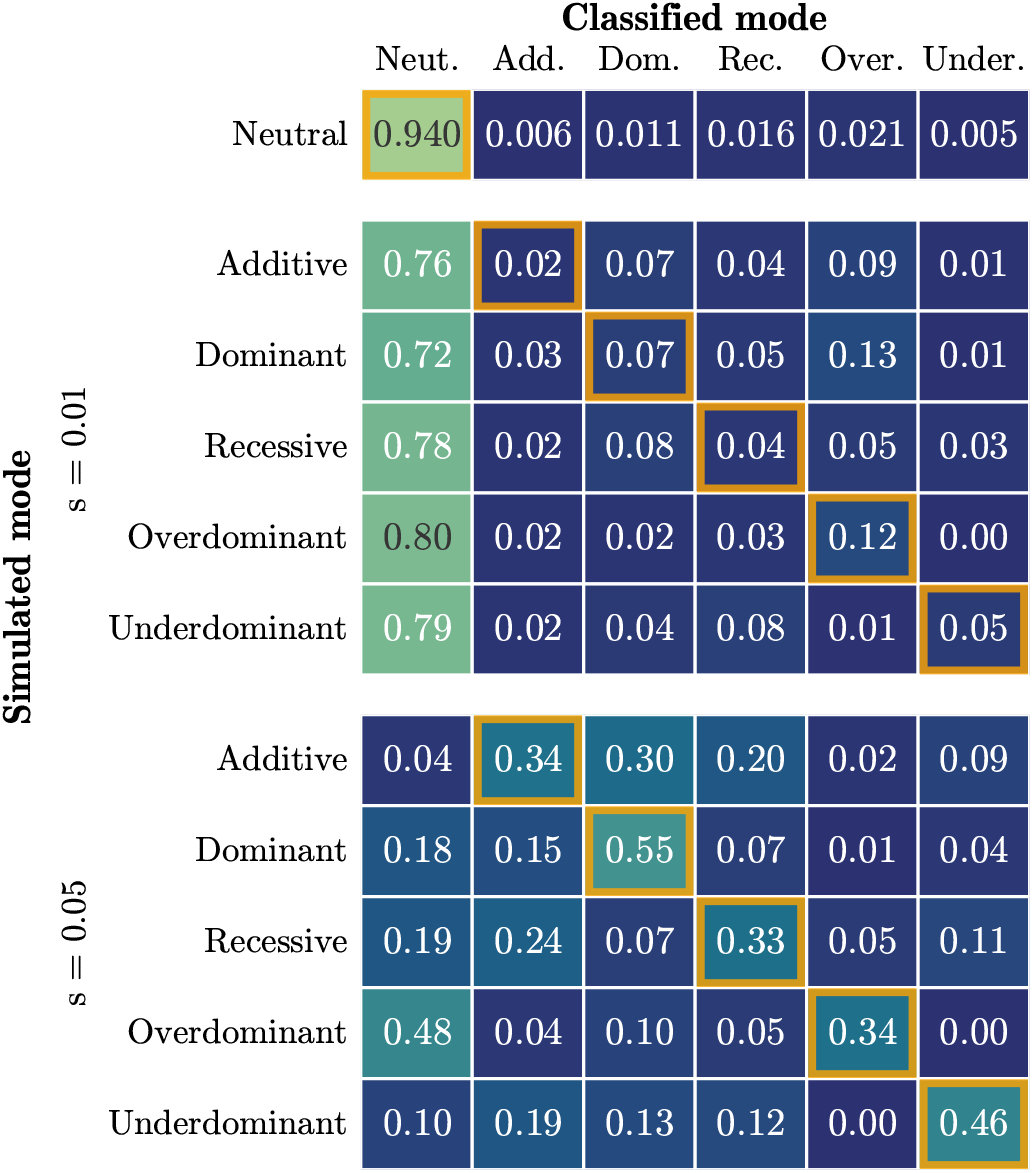
Confusion matrix for the procedure to infer selection mode applied to data-matched simulations, using a p-value threshold of 0.05. Each cell represents the fraction of replicates that were classified as a particular mode. Performance is worse than for the simulations in Section 3.1.1 – the correct model is only inferred for the plurality of replicates for *s* = 0.05. For all simulations, 10,000 replicates are simulated under a given selection mode and strength, the number of generations is *T* = 125, the value of *N*_*e*_ in each generation is derived from Browning and Browning (2015), and the initial frequencies match the GB aDNA dataset.

Lastly, Figure 11 shows the distribution of the inferred selection coefficient, conditional on inferring the correct mode of selection among multiple alternatives, as well as the distribution of the inferred selection coefficient for neutral replicates classified as non-neutral. For *s >* 0.01, estimated selection coefficients are similar to the unconditional estimates, indicating that our model inference procedure does not strongly bias the estimates in this parameter regime. However, for *s* = 0.005 and *s* = 0.01, most inferred coefficients are higher than the true value. As with the simulated data in Section 3.1.4, we observe a “winner’s curse” phenomenon for lower selection coefficients, where only replicates with extreme allele frequency changes are classified as non-neutral, and consequently *ŝ* is also large.

**Figure 11:**
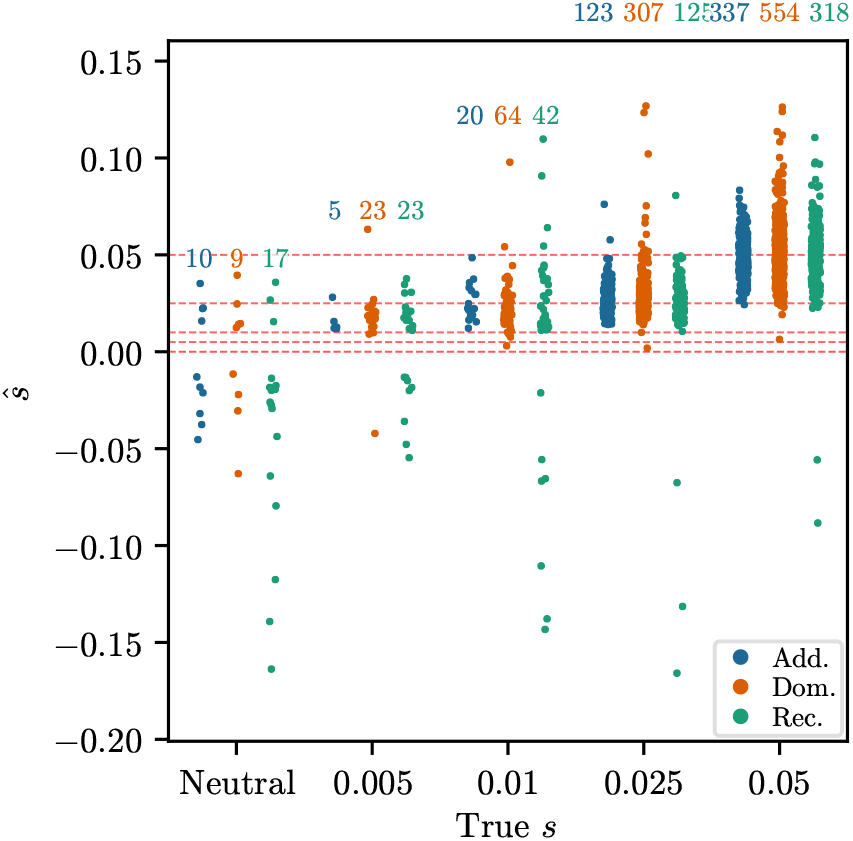
Strip plots of *ŝ* against true *s*, conditioned on inferring non-neutrality for the neutral replicates and on inferring the correct selection mode among multiple alternatives for non-neutral replicates. Each strip plot is for 1,000 replicates randomly chosen from the 10,000 simulated. Number above each strip indicates the number of replicates correctly classified. For *s* = 0.005 and *s* = 0.01, the inferred values are biased upwards, due to the “winner’s curse”. For all simulations, the number of generations is *T* = 125, the value of *N*_*e*_ in each generation is derived from Browning and Browning (2015), and the initial frequencies match the GB aDNA dataset.

#### 3.2.3 Significant signals of selection in the GB aDNA dataset

We then applied our EM-HMM algorithm and procedure to infer the mode of selection to all 743,417 SNPs in the GB aDNA dataset that pass our filters. As shown, for example, by Browning and Browning (2015), the history of effective population sizes in Great Britain varies over time. To account for this heuristically, we again mirror the strategy of Foll et al. (2014) and first estimated a single constant effective population size 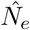 shared across SNPs, using the procedure described in Section 3.1.6, resulting in 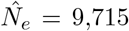. For each SNP, we then estimated the MLE using our EM-HMM for all one-parameter modes of selection, fixing the inferred 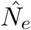 as the constant effective population size. As shown in Section 3.2.2, this heuristic to account for time-varying *N*_*e*_ yields accurate estimates of selection coefficients and well-calibrated p-values. Here, we primarily describe the results under the *additive* mode as well as the results of the procedure to infer the mode of selection. Results for the other one-parameter modes, as well as Q-Q plots of the p-values for all modes, can be found in Section S.9 of the Supplementary Material.

Figure 12 shows a Manhattan plot of the p-values for the single-alternative likelihood-ratio test computed using the *additive* EM-HMM. We also indicate the significance threshold obtained from applying the Benjamini–Hochberg (BH) procedure (Benjamini and Hochberg 1995) to correct for multiple testing at a false discovery rate (FDR) of *α* = 0.05. In Section S.10 of the Supplementary Material, we compute the same p-values when permuting the sampling times in the dataset, demonstrating that the enrichment of low p-values we observe is a reliable signal in the data. Furthermore, we observe clusters of low p-values on Chromosomes 2, 5, and 6. This clustering of p-values is expected, since due to genetic hitchhiking (Smith and Haigh 1974), SNPs that are in proximity to an actual target of selection, and thus in linkage disequilibrium (LD) with the target, will also exhibit non-neutral dynamics.

**Figure 12:**
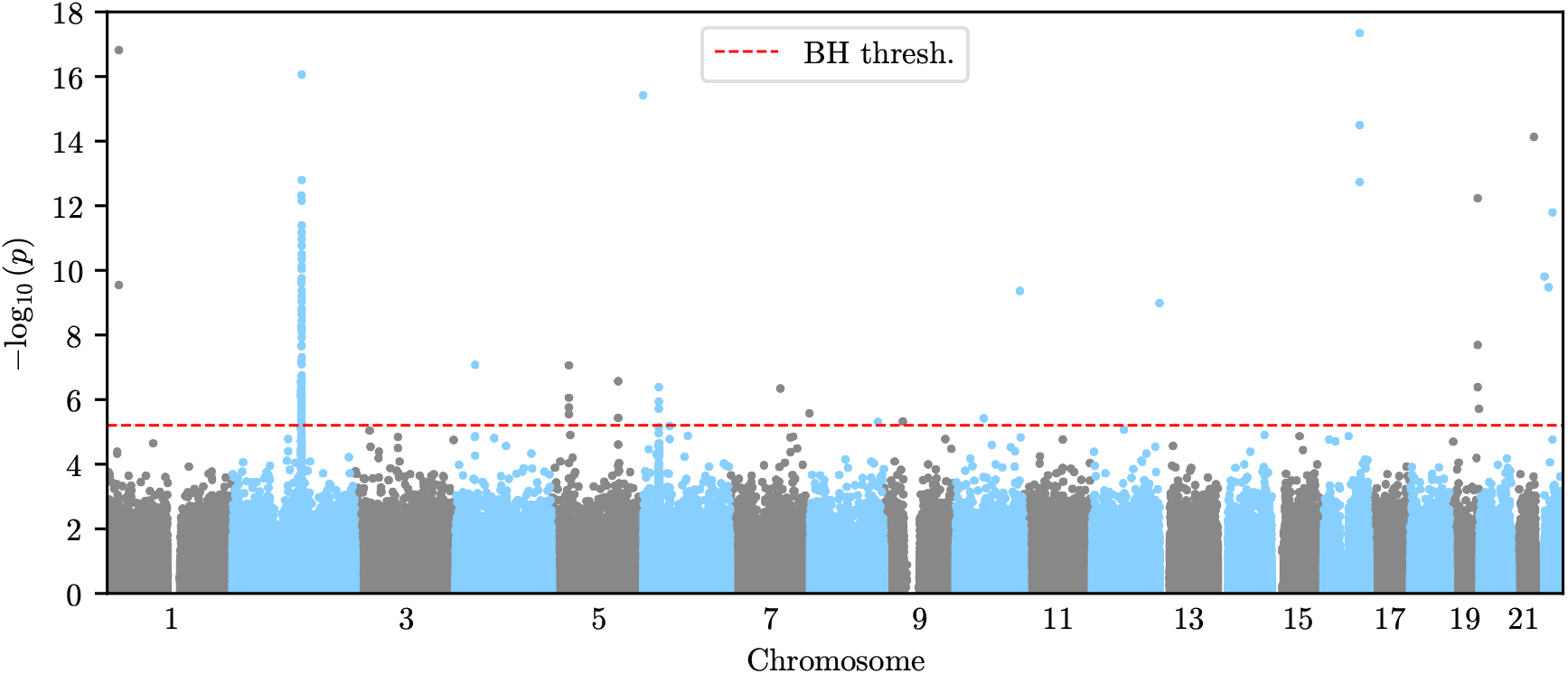
Manhattan plot of p-values obtained from the likelihood-ratio test under the *additive* mode of selection at all SNPs in the GB aDNA dataset. The significance threshold is obtained via the Ben-jamini–Hochberg procedure with an FDR of *α* = 0.05. We observe clusters of significant p-values on chromosomes 2, 5, and 6, as well as several isolated signals.

In addition to these clusters, we observe several isolated SNPs with p-values exceeding the BH threshold, but no surrounding SNPs show evidence of selection, a pattern that would not be expected under genetic hitchhiking. We thus do not believe that these isolated SNPs correspond to true signals of selection and they are likely artifacts in the dataset. However, several other regions have a SNP whose p-value exceeds the BH threshold, and multiple nearby SNPs exhibiting low p-values, thus giving some support that these potentially correspond to real signals of selection. Figures 13A and 13B show the p-values in a genomic region with low p-values surrounding a significant SNP on chromosome 5 and a region surrounding a SNP with a spuriously low p-value on chromosome 7, respectively.

**Figure 13:**
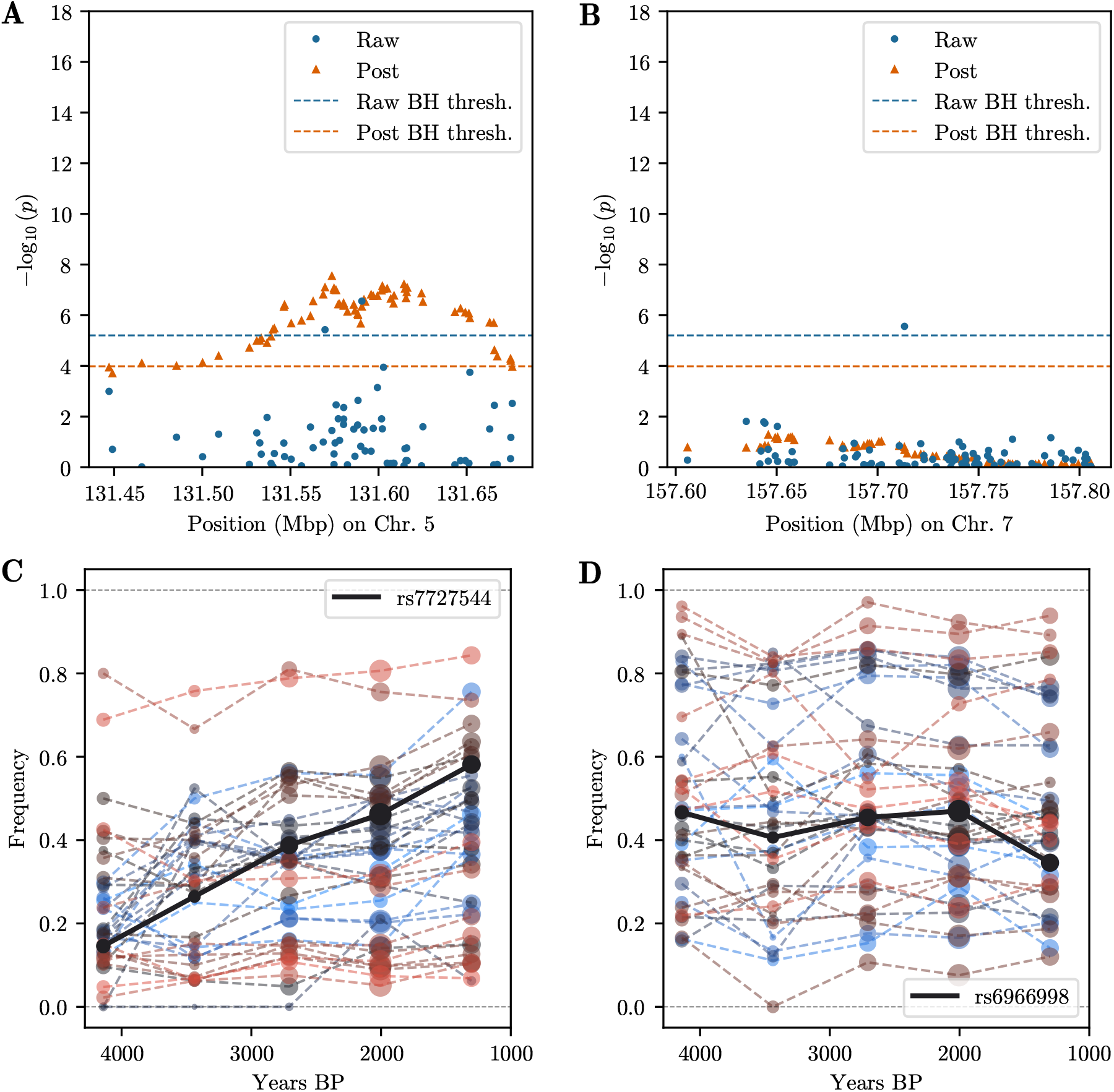
A) Manhattan plot of p-values in genomic region on chromosome 5, centered around SNPs with p-value exceeding BH threshold. Surrounding SNPs exhibit low p-values, as expected under genetic hitchhiking. Post-processed p-values exceed the respective BH threshold. B) Manhattan plot of p-values in genomic region on chromosome 7, centered around SNP with p-value exceeding BH threshold. Surrounding SNPs do not show evidence of selection. Post-processed p-values do not show any significant signal. Binned allele frequency trajectories for 20 SNPs centered around SNP with lowest p-value in Figure 13A. Nearby SNPs show correlated allele frequency change, indicative of genetic hitchhiking and true signal. Size of points indicates the number of samples; color hue indicates genomic position: red smaller, blue larger. Binned allele frequency trajectories for 20 SNPs centered around SNP with lowest p-value in Figure 13B. Nearby SNPs do not exhibit correlated allele frequency change, suggesting spurious signal at lead SNP.

To remove spurious SNPs while keeping significant SNPs in regions that show additional support at surrounding SNPs, we post-process the p-values using a modified version of Brown’s method (Brown 1975) for combining non-independent p-values. Specifically, we consider overlapping windows of 50 SNPs around each analyzed SNP. For each of these overlapping windows, we compute the negative sum of the logarithm of the p-values, including the focal SNP. We then fit the parameters of a scaled *χ*^2^ distribution to these sums, and use this fitted distribution to compute post-processed p-values for each SNP. We apply the BH method to this new set of p-values to obtain a second BH threshold. Isolated SNPs, such as the SNP in Figure 13B no longer exceed the corresponding BH threshold, while regions containing a significant SNP and additional support, such as the region in Figure 13A, have a broad peak exceeding the corresponding BH threshold.

After applying this post-processing procedure, we group significant p-values into distinct regions of significance. For a region to be considered significant, we require that at least one SNP within the region must have both a raw p-value and a post-processed p-value each exceeding its respective BH threshold. The post-processing procedure broadens p-value peaks; we therefore take each contiguous block of post-processed p-values exceeding the corresponding BH threshold as a separate candidate region. Under this criterion, for the one-dimensional *additive* EM-HMM algorithm, there are 8 distinct candidate regions. Table 1 lists these regions (in hg19 coordinates), any genes overlapping the region, the rsID of the SNP corresponding to the minimal p-value in the region, the reference and alternative alleles at this SNP, the number of significant SNPs (pre- and post-processing) in each region, the negative logarithm of the minimal p-value in the region, and the *additive ŝ* for the derived allele at the SNP with the lowest p-value with confidence intervals. Confidence intervals were obtained by simulating 1,000 replicates matching the sampling scheme, estimated initial frequency, and estimated selection coefficient of each lead SNP, using 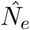, then bias-correcting all *ŝ* values and taking the 0.025 and 0.975 quantiles of the simulated replicates. Additionally, we present Manhattan plots of the p-values and the binned allele frequency trajectories in each significant genomic region in Section S.11 of the Supplementary Material.

**Table 1:** Genomic regions identified as significant under *additive* selection. We indicate genes that overlap the region, number of SNPs exceeding the BH threshold pre- and post-processing, − log_10_ *p*_*min*_, and *ŝ* for the derived allele at the SNP with − log_10_ *p*_*min*_. The asterisks at the HYDIN locus indicate that this signal is likely an artifact.

| Chr. | Genomic region (hg19) | Gene(s) | Lead SNP | Ref. | Alt. | Raw | Post | $-\log_{10} p_{min}$ | $\hat{s}(p_{min})$ |
| --- | --- | --- | --- | --- | --- | --- | --- | --- | --- |
| 2 | 135,161,631-137,087,324 | LCT | rs4988235 | G | A | 63 | 395 | 16.05 | 0.080 (0.057, 0.099) |
| 2 | 137,164,476-137,419,024 | LCT | rs580879 | C | T | 1 | 76 | 5.23 | -0.043 (-0.061, -0.022) |
| 4 | 38,680,186-38,806,462 | TLR10/1/6 | rs10008492 | C | T | 1 | 54 | 7.07 | -0.047 (-0.062, -0.026) |
| 5 | 33,854,740-34,004,707 | SLC45A2 | rs16891982 | C | G | 4 | 51 | 7.05 | 0.049 (0.029, 0.066) |
| 5 | 131,465,688-131,675,046 | SLC22A4 | rs7727544 | C | T | 2 | 67 | 6.56 | 0.042 (0.023, 0.056) |
| 6 | 31,008,368-31,091,197 | HLA | rs2535317 | C | T | 3 | 151 | 6.38 | -0.046 (-0.062, -0.025) |
| 7 | 100,212,254-100,361,675 | TFR2 | rs4434553 | A | G | 1 | 27 | 6.33 | 0.044 (0.025, 0.061) |
| **16 | 70,771,279-71,330,074 | HYDIN | rs79233902 | T | G | 3 | 51 | 17.34 | 0.242** |

#### 3.2.4 Functional genetic variation in significant genomic regions

With the exception of TFR2, all genomic regions that we detect as significant by applying our *additive* EM-HMM have previously been identified as targets of selection, although not all have been identified as such in populations from Great Britain. In the following paragraphs, we will discuss each of these regions in the context of the relevant literature.

The strongest signal in our analysis is the well-characterized LCT locus, which has been identified as a target of selection in populations from Great Britain (Mathieson and Terhorst 2022; Nait Saada et al. 2020; Field et al. 2016) and broader Western European populations (Bersaglieri et al. 2004; Itan et al. 2009; Mathieson et al. 2015). The lead SNP in our analysis, rs4988235, has previously been identified as the strongest signal of selection at the LCT locus (Itan et al. 2009; Peter et al. 2012; Mathieson and Terhorst 2022), and the derived allele at this SNP has been linked to the ability to digest lactase into adulthood (Enattah et al. 2002). Our estimated *additive* selection coefficient of 0.080 (CI: [0.057, 0.099]) is in line with other estimates provided in the literature (Peter et al. 2012; Mathieson and Terhorst 2022; Bersaglieri et al. 2004). Although the derived allele at this SNP has been linked to lactase persistence, recent studies argue that the introduction of milk consumption predates the increase in frequency of this allele, and that the recent strong selective pressure perhaps results from later famines where the allele proved beneficial (Burger et al. 2020; Evershed et al. 2022). Our approach also identified a secondary region of significance near the LCT locus which is likely a result of genetic hitchhiking.

Genomic variation in two of our candidate regions, TLR10/1/6 and HLA, is involved in immune regulation. Polymorphisms in the TLR10/1/6 gene cluster have been linked to incidence of several cancers, as well as tuberculosis and leprosy (Purdue et al. 2009; Sun et al. 2006; Ma et al. 2007; Wong et al. 2010). The TLR10/1/6 locus has been identified as a target of selection in a previous study using ancient DNA (Mathieson et al. 2015), although this work pooled samples from West Eurasia. Similarly, using a dataset of present-day individuals, Barreiro et al. (2009) found that the TLR genes, with the exception of TLR1, experience strong negative selection, and that the TLR10/1/6 cluster has undergone recent selection in non-African populations.

The HLA locus spans a large region on chromosome 6 and encodes a set of highly polymorphic genes critical for the function of the innate immune system. Previous studies have identified multiple signals of selection in this region (Mathieson and Terhorst 2022; Field et al. 2016), and the SNPs we identify overlap with one of these signals. Individual loci within the HLA region are believed to be under balancing or frequency-dependent selection to increase allelic diversity (Hedrick and Thomson 1983; Bronson et al. 2013). For that reason, it is somewhat surprising that genomic scans for positive selection, here and in the literature (Mathieson and Terhorst 2022; Field et al. 2016), find signals at these loci. Additionally, our scan for *heterozygote difference*, which includes *overdominance*, a model of balancing selection, reveals no significant signals in the HLA region. The likely explanation is that several alleles changing in frequency are detected as positive selection, but the short time horizon considered here is not sufficient to detect alleles under long-term balancing selection.

The SLC45A2 locus has also been well-characterized as a target of selection in European populations (Lao et al. 2007; Beleza et al. 2013). Polymorphism at this locus is associated with differences in hair and skin pigmentation (Soejima and Koda 2007; Hysi et al. 2018). Studies using ancient DNA data from both Western Europe broadly and Great Britain specifically have identified the SLC45A2 locus as a target of selection (Mathieson et al. 2015; Mathieson and Terhorst 2022). In addition, our estimated selection coefficient at the lead SNP matches values obtained from both analyses of present-day and ancient DNA: Our MLE is *s* = 0.049 (CI: [0.029, 0.066]), Beleza et al. (2013) estimate *s* in the range of 0.04 to 0.05, and Mathieson and Terhorst (2022) estimate *s* = 0.043.

The gene SLC22A4 is contained within the larger IBD5 locus, which consists of a group of genes with polymorphisms linked to gastrointestinal disorders such as Crohn’s disease (Fisher et al. 2006). Huff et al. (2012) found that genetic variants in SLC22A4 increases absorption of the antioxidant ergothioneine and show signals of positive selection, likely due to the low amounts of ergothioneine in early Neolithic farmer diets. Furthermore, they argue that variants linked to Crohn’s disease likely increased in frequency via genetic hitchhiking. The SLC22A4/IBD5 locus was also identified as a target of selection using ancient DNA from Western Europeans (Mathieson et al. 2015).

Genetic variation at the TFR2 locus has not been identified as a target of selection using either contemporary or ancient genomic samples. Figure S.36 in the Supplementary Material shows that the allele frequency at the lead SNP and surrounding SNPs shift in concert indicating that this is potentially a real target of selection rather than a false signal. Mutations at the TFR2 locus cause type 3 hereditary hemochromatosis, which is characterized by abnormally high systemic iron levels (Girelli et al. 2002). In addition, a haplotype that includes the variant at the lead SNP we identify as under selection has been correlated with Parkinson’s disease (Rhodes et al. 2014).

The HYDIN locus contains the SNP with the most significant p-value in our dataset. However, upon inspection of the p-values in this genomic region, provided in Figure S.37A in the Supplementary Material, we observe that the locus indeed contains three SNPs with very low p-values, but also two 100 kbp regions without any SNPs. These empty regions are a result of our filtering procedure, which removed a large number of SNPs with minor allele frequency below 0.05. In addition, the significant SNPs at the HYDIN locus have an extremely low binned minor allele frequency at all but the last timepoint, see Figure S.37B in the Supplementary Material. Since the gene HYDIN on chromosome 16 has a pseudogene on chromosome 1 (Dutcher and Brody 2019), these unusual patterns are potentially a result of mismapped sequence reads, and we thus believe that the signal of selection at the HYDIN locus is spurious.

Lastly, we more explicitly compare the results of our analysis to those of Mathieson and Terhorst (2022), a recent study that identified targets of selection in a temporal dataset similar to ours. The authors analyze present-day and ancient DNA samples from the AADR localized to England, dated to under 4,450 years BP, and find signals of selection in five genomic regions. Three regions (LCT, SLC45A2, HLA) are identified in our study as well, three regions (TLR10/1/6, SLC22A4, TFR2) are only identified in our analysis, and two regions (DHCR7, HERC2) are only identified by Mathieson and Terhorst. Of the loci identified in both studies, the estimates of the selection coefficients at the most significant SNPs largely agree: the coefficients in the LCT, SLC45A2, and HLA regions are 0.080 (CI: [0.057, 0.099), 0.049 (CI: [0.029, 0.066]), and 0.046 (CI: [0.025, 0.061]) in our study and 0.064, 0.043, and 0.046 in Mathieson and Terhorst’s study, respectively. Figures S.38 and S.39 in the Supplementary Material show the p-values computed using our *additive* EM-HMM and binned allele frequency trajectories in both regions identified as significant by Mathieson and Terhorst (2022) that are not significant in our analysis. At the DHCR7 locus on chromosome 11, the post-processed p-value are close to exceeding the BH threshold, but no raw p-value reaches significance. The HERC2 locus on chromosome 15 also exhibits low p-values, although to a lesser degree than DHCR7. In contrast to our conservative approach of only analyzing samples genotyped using the 1240K assay, Mathieson and Terhorst also analyze samples genotyped using whole genome data (including present-day samples), use a method that has the potential to detect selection where the *additive* coefficient changes over time, and use a different post-processing when combining signals at neighboring SNPs. Thus, a perfect alignment of signals is not expected.

#### 3.2.5 Inferring the mode of selection in the GB aDNA dataset

In addition to the *additive* EM-HMM, we analyzed the GB aDNA dataset under each of the other three one-parameter modes of selection – *dominant, recessive*, and *heterozygote difference*, see Section S.9 in the Supplementary Material, where we provide full Manhattan plots for each one-parameter mode of selection, as well as tables analogous to Table 1. Additionally, we provide a table of genome-wide Spearman rank correlation coefficients between the log-likelihood ratio statistics of all one-parameter modes for the top 1% of SNPs by *additive* p-value in Figure S.28 of the Supplementary Material. Both the *dominant* and the *recessive* EM-HMM have high correlation with the *additive* EM-HMM, and identify the same regions as the *additive* analysis, although the LCT locus is not split into two distinct regions in the *dominant* analysis.

In contrast, the *heterozygote difference* EM-HMM has a lower correlation with the other modes and only identifies LCT, the two SLC loci, and HYDIN as significant. The inability of this mode to detect selection at the HLA locus is at first somewhat surprising, given that balancing selection or frequency-dependent selection should result in a dynamic similar to *overdominant* selection. However, the signature that would provide strong support for these types of selection would be alleles that remain at intermediate frequencies longer than expected under neutrality, and thus the short time-horizon considered here is likely not sufficient.

We also used the procedure detailed in Section 2.3.2 to infer the most likely mode of selection at each SNP. We computed the *δ* statistic for all SNPs in the GB aDNA dataset and used the *χ*^2^(1) distribution to obtain p-values for each locus. We applied our procedure based on Brown’s method to identify significant regions, using a BH threshold at an FDR of *α* = 0.05. SNPs in the significant regions whose raw p-value exceeded the BH threshold were classified as the one-parameter mode of selection with the highest log-likelihood.

The mode inference procedure identifies the same set of loci as the *additive* single-alternative procedure as significant. Figure 14 shows the resulting raw p-values at the LCT locus, with significant SNPs colored by their inferred selection mode. Out of the 68 SNPs exceeding the BH threshold in this region, 30 are classified as *additive*, but the other SNPs show different selection modes. The fact that a majority of loci in this region are classified as *additive* could indicate support for the hypothesis that LCT is evolving under *additive* selection; however, the inferred mode at the lead SNP rs4988235 is *dominant*. Lactase persistence functions as a dominant trait (Swallow 2003), lending further support to the inference of *dominant* selection at the lead SNP. Furthermore, the method presented by Mathieson and Terhorst (2022) can model non-constant selection coefficients, and the authors provide evidence that the selection at the LCT locus has weakened over time; a dynamics resembling constant *dominant* selection. We do caution against over-interpretation of these results, as the simulation study in Section 3.2.2 shows that identifying even a constant mode of selection is challenging in this dataset, and thus a much greater number and density of samples is likely necessary for accurate classification. Lastly, we note that the one-parameter mode with the highest log-likelihood ratio at the lead SNP for each identified region is as follows: LCT – *dominance*, TLR10/1/6 – *additive*, SLC45A2 – *heterozygote difference*, SLC22A4 – *dominance*, HLA – *recessive*, and TFR2 – *additive*.

**Figure 14:**
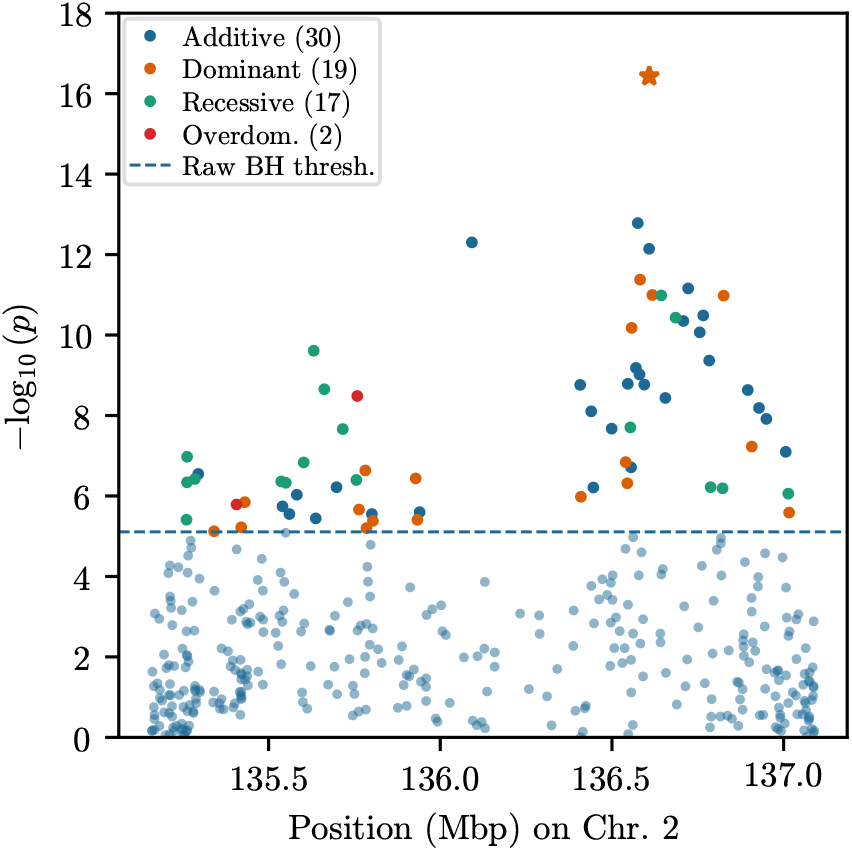
Manhattan plot of raw p-values at the LCT locus. P-values are computed using the procedure to identify the mode of selection described in Section 2.3.2, and significant SNPs are colored by inferred mode. The lead SNP is indicated by a larger star-shaped marker. The majority of SNPs in this region are classified as *additive*, although the lead SNP is classified as *dominant*.

### 3.3 Coat coloration locus ASIP in domesticated horses

#### 3.3.1 Description of dataset

In this section, we apply out method to a dataset presented by Ludwig et al. (2009), where the authors extracted ancient DNA at six loci that affect coat coloration in domesticated horses from a set of samples in Eurasia and found evidence for selection at the ASIP and MC1R loci. Specifically, we apply our method to the ASIP locus. Figure 15A shows the sample allele frequency of the derived allele at this locus over time. The samples exhibit a sharp increase in the frequency of the derived allele, followed by leveling off at a frequency of approximately 0.5. The underlying sampled allele counts can be found in Table S.5 of the Supplementary Material.

**Figure 15:**
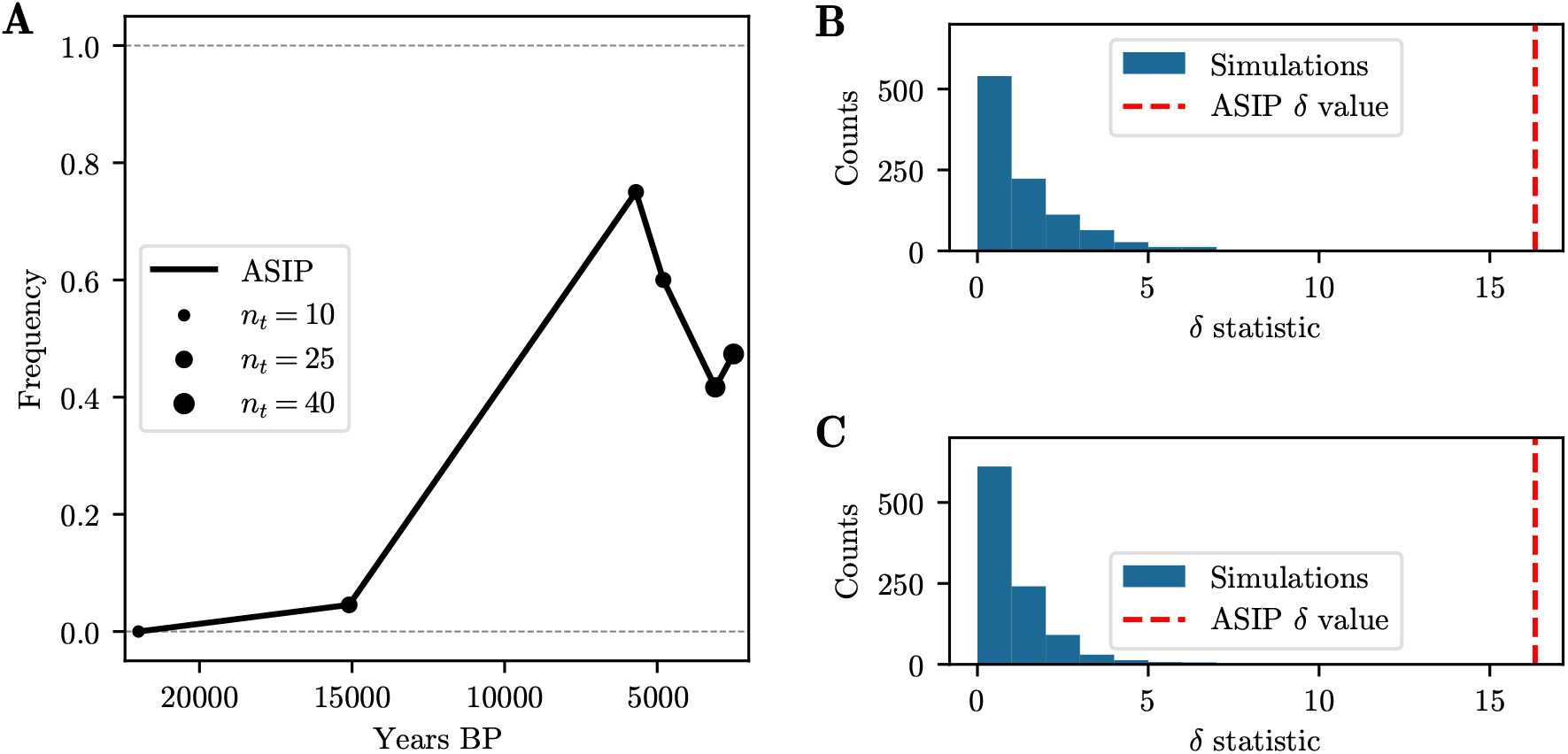
A) Derived allele frequency over time at the ASIP locus. The size of the points indicates the number of samples. B) & C) Histograms of *δ* statistic for 1,000 simulated neutral replicates matching the ASIP locus, using B) initial frequency estimated under neutrality or C) initial frequency estimated using the heterozygote difference mode. In both cases, the original dataset has a higher *δ* statistic (indicated by red dashed line) than any simulated replicate, providing strong evidence against neutrality.

This dataset has been re-analyzed in several studies under different modes of selection, with differing results; Malaspinas et al. (2012) analyzed the ASIP locus under *recessive* selection and do not find evidence for selection, Steinrücken et al. (2014) find evidence for *overdominant* selection, while more recently He et al. (2023) analyze the locus under *recessive* selection and conclude that it is not under selection. The ASIP locus is known to act via a *recessive* mechanism – horses with two copies of the derived allele are black, otherwise they are bay colored (Rieder et al. 2001). It is therefore somewhat unexpected that the ASIP locus shows evidence for *overdominant* selection and not for *recessive* selection.

#### 3.3.2 Single-alternative and multiple-alternative inference at the ASIP locus

We applied our EM-HMM to estimate the selection coefficients and parameters of the initial distribution at the ASIP locus under each one-parameter selection mode. Following He et al. (2023) we assume *N*_*e*_ = 16,000 and a generation time of 8 years. For the one-parameter modes, we use the *χ*^2^(1)-based log-likelihood ratio test to compute p-values. We report the resulting estimates of the selection coefficients, the log-likelihood differences, and the p-values in Table 2. We observe that the evidence for *overdominance* selection is strongest, with a single-alternative p-value of 5.33·10^−5^. In contrast to Malaspinas et al. (2012) and He et al. (2023), we do find that the ASIP locus shows evidence for *recessive* selection, although the p-value (2.0 · 10^−2^) is not very strong.

**Table 2:** Estimates of selection coefficients, log-likelihood differences, and p-values for all one-parameter modes. Estimates are given for both the full ASIP dataset, and the dataset truncated after the first three samples.

| | | Add. ($s_2$) | Dom. ($s_2$) | Rec. ($s_2$) | Het. diff. ($s_1$) |
| --- | --- | --- | --- | --- | --- |
| Full | $\hat{s}$ | 0.0025 | 0.0022 | 0.0023 | 0.0048 |
| | $\ell\ell - \ell\ell_0$ | 5.28 | 6.67 | 2.72 | 8.16 |
| | p-value | $1.15 \cdot 10^{-3}$ | $2.59 \cdot 10^{-4}$ | $1.96 \cdot 10^{-2}$ | $5.33 \cdot 10^{-5}$ |
| Trunc. | $\hat{s}$ | 0.0057 | 0.0043 | 0.0090 | 0.0047 |
| | $\ell\ell - \ell\ell_0$ | 10.52 | 10.38 | 8.63 | 7.57 |
| | p-value | $4.51 \cdot 10^{-6}$ | $5.21 \cdot 10^{-6}$ | $3.26 \cdot 10^{-5}$ | $9.98 \cdot 10^{-5}$ |

Although the estimated selection coefficients are low for all modes of selection (e.g. *ŝ* = 0.0048 for *heterozygote difference*), the dataset comprises roughly 2,500 generations, which is over ten times as many generations as the GB aDNA dataset. This increases power to detect weaker selection; for example, the AUC values for *s* = 0.005 for the 100-generation simulation in Figure S.2C of the Supplementary Material are only slightly above 0.5, indicating minimal power to detect selection, whereas those in the 1000-generation simulation plotted in Figure S.7C are all above 0.9.

In addition to the single alternative tests, we also computed the test statistic *δ* for multiple alternatives (see Section 2.3.2), but used parametrized bootstrap simulations to assess its significance. To this end, we simulated two sets of 1,000 neutral replicates matching the sampling scheme and number of generations to the original data at the ASIP locus. One set was simulated using the initial frequency estimated under the neutral EM-HMM and the other set with the initial frequency from the *heterozygote difference* EM-HMM, the mode with the highest likelihood among the one-parameter modes. We simulate these two sets to cover different plausible scenarios. Figures 15B and 15C show histograms of the *δ* statistics for the 1,000 simulated replicates using the neutral initial frequency and the *heterozygote difference* initial frequency, respectively, with the *δ* statistic of the original data indicated by a vertical line. In both cases, the *δ* statistic of the original data is much larger than any of the simulated replicates, indicating significant evidence for non-neutral dynamics. *Overdominance* selection (specifically, the *heterozygote difference* one-parameter mode with *s*_1_ *>* 0) has the highest log-likelihood. Thus, the data at the ASIP locus supports *overdominance* selection as the most likely mode. Additionally, *recessive* selection has the lowest log-likelihood out of all one-parameter modes of selection. Since the data shows a sharp increase in frequency, followed by a plateau around 0.5, it is expected that *overdominance* is most strongly supported.

The genetic mechanism of the derived allele at the ASIP locus is *recessive*, yet the data strongly suggests that *overdominance* is the most and *recessive* selection is the least likely mode of selection. We propose the following two possible explanations for this discrepancy. First, the selection coefficient of the derived allele may have decreased during horse domestication, and some form of balancing selection might be acting after the initial increase. The point estimates of He et al. (2023) indeed suggest stronger selection prior to domestication, followed by weaker selection thereafter; this is also consistent with the findings of Wutke et al. (2016). To explore this hypothesis further, we truncated the data at the ASIP locus after the first three timepoints, to analyze just the period of frequency increase, and report the results in Table 2. We find that, indeed, the evidence for *recessive* selection is stronger than the evidence for *overdominance*, however, *additive* and *dominant* selection are also significant and have stronger p-values. A second possible explanation for our findings could be epistasis, since the derived allele at the ASIP locus has epistatic interactions with another coat-coloration locus that has been shown to be under selection, the MC1R locus (Rieder et al. 2001), which could affect the effective mode of selection. Future work may resolve these questions, but additional samples are likely necessary.

## 4 Discussion

In this work, we presented a novel method to compute maximum likelihood estimates (MLEs) for general diploid selection coefficients from time series genetic data. To this end, we extended the framework of Mathieson and McVean (2013) for the *additive* case and derived an EM-HMM algorithm to estimate the parameters of diploid selection. We show that the diploid EM-HMM framework can also be constrained to bespoke one-parameter models of selection via the method of Lagrange multipliers. We furthermore introduced a novel likelihood-based procedure for inferring the best fitting diploid mode of selection from temporal data between *additive, recessive, dominant*, and *over-* or *underdominant* selection. To our knowledge, our study is the first to address the statistical problem of explicitly determining the mode of selection from given time series genetic data. Additionally, we implement a method to estimate a constant population size *N*_*e*_ for a given dataset, allowing for better modeling of the dynamics of genetic drift in the HMM. To further improve power to detect selection and remove spurious signals, we also introduced a procedure based on Brown’s method to combine p-values across linked loci.

Using simulation studies, we show that the estimated selection coefficients are accurate across a range of selection parameters, population parameters, and sampling schemes. However, we find that determining the mode of selection from time series data is challenging, and only yields reliable results when selection is strong. We also demonstrate that assuming *additive* selection when analyzing data simulated under different modes of selection yields comparable power to reject neutrality as when the data is simulated under *additive* selection, implying that analyzing given data assuming *additive* selection may be sufficient for scans of directional selection. However, the estimated selection coefficients are inaccurate when the mode is misspecified. In addition, we demonstrate that our procedure to account for variable population size leads to well-calibrated estimates and p-values. However, this is likely related to the short time horizon and the fact that the population size steadily increases from *N*_*e*_ ≈ 10,000 to *N*_*e*_ ≈ 400,000 (see Figure S.15 in the Supplementary Material). A more extreme history like exponential growth or severe bottlenecks will likely be more challenging, and a practitioner would have to re-assess the method in such a scenario.

We apply our method to time series genetic data obtained from 504 ancient individuals in the AADR (Mallick et al. 2024) from Great Britain dated to under 4,450 BP, and identify six genomic regions with signals of selection. These regions, except TFR2, have been identified as targets of selection in previous studies, and we discuss them in the context of the relevant literature. The regions are identified as significant under multiple directional selection modes (*additive, recessive, dominant*). When classifying the mode of selection from the data, however, the results are inconclusive: For example, we find that a majority of the SNPs at the LCT locus provide evidence for *additive* selection, but the lead SNPs is classified as *dominant*. In addition, we reanalyze a time-series dataset consisting of 146 samples over 2,400 generations from the ASIP locus involved in coat coloration in horses, and show evidence for selection under different non-additive modes.

Note that our HMM-implementation uses the Chebychev nodes to compute the single-generation transition matrix accurately across the hidden state space, and the integrals account for the probability mass that is absorbed in the boundary. Capturing these features accurately is important when implementing the model (Tataru et al. 2017; Paris et al. 2019), and consequently, we believe that our algorithm identifies the MLEs from a given temporal datasets and computes the likelihoods with high accuracy. Thus, the statistical properties of the MLEs and the likelihood-ratio tests exhibited in our simulation studies in Section 3.1 and Section 3.2.2 are likely not exclusive to our method but potentially characterize the MLEs and the power of the respective tests in general under the given population genetic model, regardless whether our method or a different likelihood-based method is used for the analysis.

Moreover, we characterize the statistical properties in a range of scenarios, but if these scenarios do not cover the exact scheme encountered in a specific empirical dataset, our simulation framework can be readily modified to characterize the statistical properties in the respective scenario. Naturally, the power to identify and characterize selection does depend on the exact sampling scheme: Strong selection can be readily identified, even when samples are limited to a short time period. However, weaker selection requires sampling more data over a longer time period. For example, in our analysis of the GB aDNA dataset, Figure 9C and Figure 10 demonstrate limited power to detect selection as strong as *s* = 0.01 in the respective scenario.

Our method computes the MLEs of general diploid selection parameters, and we believe that this is useful to researchers in at least two regards: (1) Our approach can be used to infer the mode of selection from a given temporal dataset. While we demonstrate that selection needs to be fairly strong for reliable classification, our framework can be used to characterize statistical power in a given scenario, and determine whether additional samples at potentially additional timepoints are necessary. (2) If the selection mode operating on the genetic variants in a given temporal dataset is known a priori, for example, *dominance* at the LCT locus or *underdominance* dynamics resulting from stabilizing selection on complex traits, our method enables researchers to estimate the selection coefficients accurately under the correct model. We demonstrate that assuming the wrong mode of selection can yield inaccurate estimates.

In practice, we recommend the following approach to analyze given time series genetic data, potentially at a large number of loci: If computational resources are limited, researchers should apply the *additive* EM-HMM to obtain MLEs of the *additive* selection coefficient at each locus, and use standard likelihood-ratio testing to identify outliers. As detailed in Section 3.1.5, the likelihood-ratio test under *additive* selection can identify non-neutral replicates, even if the mode of selection is misspecified, but estimated coefficients are inaccurate. For reference, the *additive* analysis of the 743,417 SNPs for 504 samples over 125 generations in our GB aDNA dataset took roughly 5,000 cpu-hours. If additional computational resources are available, we recommend analyzing the data under each bespoke one-parameter selection mode, as well as the unconstrained mode, to characterize signals in the data that are not correctly described by *additive* selection. The results can then also be used to identify the mode of selection from the data, but as we demonstrate, accuracy is limited. In addition, we strongly recommend performing a data-matched simulation study, as presented in Section 3.2.2 or Section S.6 in the Supplementary Material. Such data-matched simulations enable exact characterization of the statistical power and accuracy of the approach in the specific scenario.

As for similar approaches, the HMM underlying our method assumes that the population is panmictic, and violations of this assumption can dilute signals or introduce spurious signals. Future work to address this shortcoming could proceed along at least two possible avenues: (1) Controlling for population structure by using Principal Components as covariates in the estimation procedure directly (Luu et al. 2017; Ju and Mathieson 2021), or (2) explicitly including the population structure and exchange of migrants in the underlying population genetic model (Joseph and Pe’er 2019). Moreover, our approach estimates selection coefficients from temporal data at the focal locus only, and does not incorporate the allele frequency dynamics at linked loci. In our analysis of the GB aDNA dataset, we do leverage signals across loci using a novel post-processing approach to combine p-values in a genomic window. This post-processing can reduce signal-to-noise ratios in genomic scans for selection in general. However, incorporating the genetic variation at multiple SNPs using a proper likelihood model for the multi-locus dynamics under the Wright-Fisher process has the potential to account for chromosomal linkage more accurately and result in a more robust inference (Terhorst et al. 2015; He et al. 2020).

While we focus on analyzing time series genetic data at a single locus in this study, the capacity of our method to characterize selection modes more general than *additive* only also has potential benefits when studying polygenic selection on complex traits: In models of stabilizing polygenic selection around an optimal trait, the genetic variants affecting the trait experience *underdominant* selection dynamics, which can be readily addressed using our framework.

## Acknowledgements

We want to thank Xinyi Li, Xiaoheng Cheng, Constanza de la Fuente, and Maanasa Raghavan for helpful comments on the method and the data analysis. Moreover, we thank Jeremy Berg, Maryn Carlson and Rowan Hart for comments on the manuscript. In addition, we thank the members of the Raghavan, Berg, and Novembre labs for valuable feedback throughout the project.

## List of Supplementary Files

**Supplementary Material S1** – Document containing additional details of the method, as well as figures and tables to supplement the analyses.

## References

Barghi, Neda, Raymond Tobler, Viola Nolte, Ana Marija Jakšić, François Mallard, Kathrin Anna Otte, Marlies Dolezal, Thomas Taus, Robert Kofler, and Christian Schlötterer (2019). “Genetic Redundancy Fuels Polygenic Adaptation in Drosophila”. PLOS Biology 17(2), e3000128.

Barreiro, Luis B., Meriem Ben-Ali, Hélène Quach, Guillaume Laval, Etienne Patin, Joseph K. Pickrell, Christiane Bouchier, Magali Tichit, Olivier Neyrolles, Brigitte Gicquel, Judith R. Kidd, Kenneth K. Kidd, Alexandre Alcaïs, Josiane Ragimbeau, Sandra Pellegrini, Laurent Abel, Jean-Laurent Casanova, and Lluís Quintana-Murci (2009). “Evolutionary Dynamics of Human Toll-Like Receptors and Their Different Contributions to Host Defense”. PLOS Genetics 5(7), e1000562.

Barton, Nick (1986). “The Maintenance of Polygenic Variation through a Balance between Mutation and Stabilizing Selection”. Genetical Research 47(3), pp. 209–216.

Beleza, Sandra, António M. Santos, Brian McEvoy, Isabel Alves, Cláudia Martinho, Emily Cameron, Mark D. Shriver, Esteban J. Parra, and Jorge Rocha (2013). “The Timing of Pigmentation Lightening in Europeans”. Molecular Biology and Evolution 30(1), pp. 24–35.

Benjamini, Yoav and Yosef Hochberg (1995). “Controlling the False Discovery Rate: A Practical and Powerful Approach to Multiple Testing”. Journal of the Royal Statistical Society: Series B (Methodological) 57(1), pp. 289–300.

Bersaglieri, Todd, Pardis C. Sabeti, Nick Patterson, Trisha Vanderploeg, Steve F. Schaffner, Jared A. Drake, Matthew Rhodes, David E. Reich, and Joel N. Hirschhorn (2004). “Genetic Signatures of Strong Recent Positive Selection at the Lactase Gene”. American Journal of Human Genetics 74(6), pp. 1111–1120.

Bignell, Graham R. et al. (2010). “Signatures of Mutation and Selection in the Cancer Genome”. Nature 463, pp. 893–898.

Bishop, Christopher M. (2006). Pattern Recognition and Machine Learning. Information Science and Statistics. New York: Springer.

Bollback, Jonathan P., Thomas L. York, and Rasmus Nielsen (2008). “Estimation of 2Nes from Temporal Allele Frequency Data”. Genetics 179(1), pp. 497–502.

Bronson, Paola G., Steven J. Mack, Henry A. Erlich, and Montgomery Slatkin (2013). “A Sequence-Based Approach Demonstrates That Balancing Selection in Classical Human Leukocyte Antigen (HLA) Loci Is Asymmetric”. Human Molecular Genetics 22(2), pp. 252–261.

Brown, Morton B. (1975). “A Method for Combining Non-Independent, One-Sided Tests of Significance”. Biometrics 31(4), pp. 987–992.

Browning, Sharon R. and Brian L. Browning (2015). “Accurate Non-parametric Estimation of Recent Effective Population Size from Segments of Identity by Descent”. American Journal of Human Genetics 97(3), pp. 404–418.

Burger, Joachim et al. (2020). “Low Prevalence of Lactase Persistence in Bronze Age Europe Indicates Ongoing Strong Selection over the Last 3,000 Years”. Current Biology 30(21), 4307–4315.e13.

Bustamante, Carlos D., Adi Fledel-Alon, Scott Williamson, Rasmus Nielsen, Melissa Todd Hubisz, Stephen Glanowski, David M. Tanenbaum, Thomas J. White, John J. Sninsky, Ryan D. Hernandez, Daniel Civello, Mark D. Adams, Michele Cargill, and Andrew G. Clark (2005). “Natural Selection on Protein-Coding Genes in the Human Genome”. Nature 437, pp. 1153–1157.

Cheng, Xiaoheng and Matthias Steinrücken (2023). “diplo-locus: A lightweight toolkit for inference and simulation of time-series genetic data under general diploid selection”. bioRxiv. 10.1101/2023.10.12.562101.

Chintalapati, Manjusha, Nick Patterson, and Priya Moorjani (2022). “The Spatiotemporal Patterns of Major Human Admixture Events during the European Holocene”. eLife 11, e77625.

Dehasque, Marianne, María C. Ávila-Arcos, David Díez-del-Molino, Matteo Fumagalli, Katerina Guschanski, Eline D. Lorenzen, Anna-Sapfo Malaspinas, Tomas Marques-Bonet, Michael D. Martin, Gemma G. R. Murray, Alexander S. T. Papadopulos, Nina Overgaard Therkildsen, Daniel Wegmann, Love Dalén, and Andrew D. Foote (2020). “Inference of Natural Selection from Ancient DNA”. Evolution Letters 4(2), pp. 94–108.

De Vladar, Harold P. and Nick Barton (2014). “Stability and Response of Polygenic Traits to Stabilizing Selection and Mutation”. Genetics 197(2), pp. 749–767.

Dutcher, Susan K. and Steven L. Brody (2019). “HY-DIN’ in the Cilia: Discovery of Central Pair– related Mutations in Primary Ciliary Dyskinesia”. American Journal of Respiratory Cell and Molecular Biology 62(3), pp. 281–282.

Enattah, Nabil Sabri, Timo Sahi, Erkki Savilahti, Joseph D. Terwilliger, Leena Peltonen, and Irma Järvelä (2002). “Identification of a Variant Associated with Adult-Type Hypolactasia”. Nature Genetics 30(2), pp. 233–237.

Evershed, Richard P. et al. (2022). “Dairying, Diseases and the Evolution of Lactase Persistence in Europe”. Nature 608, pp. 336–345.

Ewens, Warren J. (2004). Mathematical Population Genetics. 2nd ed. Vol. I. Theoretical Introduction. Springer.

Ferrer-Admetlla, Anna, Christoph Leuenberger, Jeffrey D. Jensen, and Daniel Wegmann (2016). “An Approximate Markov Model for the Wright–Fisher Diffusion and Its Application to Time Series Data”. Genetics 203(2), p. 831.

Field, Yair, Evan A Boyle, Natalie Telis, Ziyue Gao, Kyle J. Gaulton, David Golan, Loic Yengo, Ghislain Rocheleau, Philippe Froguel, Mark I. McCarthy, and Jonathan K. Pritchard (2016). “Detection of Human Adaptation during the Past 2000 Years”. Science 354, pp. 760–764.

Fisher, Sheila A., Jochen Hampe, Clive M. Onnie, Mark J. Daly, Christine Curley, Shaun Purcell, Jeremy Sanderson, John Mansfield, Vito Annese, Alastair Forbes, Cathryn M. Lewis, Stefan Schreiber, John D. Rioux, and Christopher G. Mathew (2006). “Direct or Indirect Association in a Complex Disease: The Role of SLC22A4 and SLC22A5 Functional Variants in Crohn Disease”. Human Mutation 27(8), pp. 778–785.

Foll, Matthieu, Yu-Ping Poh, Nicholas Renzette, Anna Ferrer-Admetlla, Claudia Bank, Hyunjin Shim, Anna-Sapfo Malaspinas, Gregory Ewing, Ping Liu, Daniel Wegmann, Daniel R. Caffrey, Konstantin B. Zeldovich, Daniel N. Bolon, Jennifer P. Wang, Timothy F. Kowalik, Celia A. Schiffer, Robert W. Finberg, and Jeffrey D. Jensen (2014). “Influenza Virus Drug Resistance: A Time-Sampled Population Genetics Perspective”. PLOS Genetics 10(2), e1004185.–.

Foll, Matthieu, Hyunjin Shim, and Jeffrey D. Jensen (2015). “WFABC: a Wright–Fisher ABC-based approach for inferring effective population sizes and selection coefficients from time-sampled data”. Molecular Ecology Resources 15(1), pp. 87–98.

Gemmell, Neil J. and Jon Slate (2006). “Heterozygote Advantage for Fecundity”. PLOS ONE 1(1), e125.

Girelli, Domenico, Claudia Bozzini, Antonella Roetto, Federica Alberti, Filomena Daraio, Romano Colombari, Oliviero Olivieri, Roberto Corrocher, and Clara Camaschella (2002). “Clinical and Pathologic Findings in Hemochromatosis Type 3 Due to a Novel Mutation in Transferrin Receptor 2 Gene”. Gastroenterology 122(5), pp. 1295–1302.

Gretzinger, Joscha et al. (2022). “The Anglo-Saxon Migration and the Formation of the Early English Gene Pool”. Nature 610, pp. 112–119.

He, Zhangyi, Xiaoyang Dai, Mark Beaumont, and Feng Yu (2020). “Detecting and Quantifying Natural Selection at Two Linked Loci from Time Series Data of Allele Frequencies with Forward-in-Time Simulations”. Genetics 216, pp. 521–541.

He, Zhangyi, Xiaoyang Dai, Wenyang Lyu, Mark Beaumont, and Feng Yu (2023). “Estimating Temporally Variable Selection Intensity from Ancient DNA Data”. Molecular Biology and Evolution 40(3), msad008.

Hedrick, Philip W (2012). “What Is the Evidence for Heterozygote Advantage Selection?” Trends in Ecology & Evolution 27(12), pp. 698–704.

Hedrick, Philip W and Glenys Thomson (1983). “Evidence for balancing selection at HLA”. Genetics 104(3), pp. 449–456.

Hoffmann, Laurence D., Gerald L. Bradley, and Kenneth H. Rosen (2010). Applied Calculus for Business, Economics, and the Social and Life Sciences. Expanded 10th ed. / Laurence D. Hoffmann, Gerald L. Bradley. New York, NY: McGraw-Hill.

Hofreiter, Michael, Johanna L. A. Paijmans, Helen Goodchild, Camilla F. Speller, Axel Barlow, Gloria G. Fortes, Jessica A. Thomas, Arne Ludwig, and Matthew J. Collins (2015). “The Future of Ancient DNA: Technical Advances and Conceptual Shifts”. BioEssays 37(3), pp. 284–293.

Huff, Chad D., David J. Witherspoon, Yuhua Zhang, Chandler Gatenbee, Lee A. Denson, Subra Kugathasan, Hakon Hakonarson, April Whiting, Chadwick T. Davis, Wilfred Wu, Jinchuan Xing, W. Scott Watkins, Michael J. Bamshad, Jonathan P. Bradfield, Kazima Bulayeva, Tatum S. Simonson, Lynn B. Jorde, and Stephen L. Guthery (2012). “Crohn’s Disease and Genetic Hitchhiking at IBD5”. Molecular Biology and Evolution 29(1), pp. 101–111.

Hysi, Pirro G. et al. (2018). “Genome-Wide Association Meta-Analysis of Individuals of European Ancestry Identifies New Loci Explaining a Substantial Fraction of Hair Color Variation and Heritability”. Nature Genetics 50(5), pp. 652–656.

Iranmehr, Arya, Ali Akbari, Christian Schlötterer, and Vineet Bafna (2017). “Clear: Composition of Likelihoods for Evolve and Resequence Experiments”. Genetics 206(2), pp. 1011–1023.

Itan, Yuval, Adam Powell, Mark A. Beaumont, Joachim Burger, and Mark G. Thomas (2009). “The Origins of Lactase Persistence in Europe”. PLOS Computational Biology 5(8), e1000491.

Joseph, Tyler A. and Itsik Pe’er (2019). “Inference of Population Structure from Time-Series Genotype Data”. The American Journal of Human Genetics 105(2), pp. 317–333.

Ju, Dan and Iain Mathieson (2021). “The evolution of skin pigmentation-associated variation in West Eurasia”. Proceedings of the National Academy of Sciences 118(1), e2009227118.

Koch, E., N. J. Connally, N. Baya, M. P. Reeve, M. Daly, B. Neale, E. S. Lander, A. Bloemendal, and S. Sunyaev (2024). “Genetic association data are broadly consistent with stabilizing selection shaping human common diseases and traits”. bioRxiv. 10.1101/2024.06.19.599789.

Lachance, Joseph and Sarah A. Tishkoff (2013). “Population Genomics of Human Adaptation”. Annual Review of Ecology, Evolution, and Systematics 44(1), pp. 123–143.

Lao, O., J. M. de Gruijter, K. van Duijn, A. Navarro, and M. Kayser (2007). “Signatures of Positive Selection in Genes Associated with Human Skin Pigmentation as Revealed from Analyses of Single Nucleotide Polymorphisms”. Annals of Human Genetics 71, pp. 354–369.

Ludwig, Arne, Melanie Pruvost, Monika Reissmann, Norbert Benecke, Gudrun A. Brockmann, Pedro Castaños, Michael Cieslak, Sebastian Lippold, Laura Llorente, Anna-Sapfo Malaspinas, Montgomery Slatkin, and Michael Hofreiter (2009). “Coat Color Variation at the Beginning of Horse Domestication”. Science 324(5926), p. 485.

Luu, Keurcien, Eric Bazin, and Michael G. B. Blum (2017). “pcadapt: an R package to perform genome scans for selection based on principal component analysis”. Molecular Ecology Resources 17(1), pp. 67–77.

Ma, Xin, Yuhua Liu, Brian B. Gowen, Edward A. Graviss, Andrew G. Clark, and James M. Musser (2007). “Full-Exon Resequencing Reveals Toll-Like Receptor Variants Contribute to Human Susceptibility to Tuberculosis Disease”. PLOS ONE 2(12), e1318.

Malaspinas, Anna-Sapfo (2016). “Methods to Characterize Selective Sweeps Using Time Serial Samples: An Ancient DNA Perspective”. Molecular Ecology 25(1), pp. 24–41.

Malaspinas, Anna-Sapfo, Orestis Malaspinas, Steven N. Evans, and Montgomery Slatkin (2012). “Estimating Allele Age and Selection Coefficient from Time-Serial Data”. Genetics 192(2), pp. 599–607.

Mallick, Swapan, Adam Micco, Matthew Mah, Harald Ringbauer, Iosif Lazaridis, Iñigo Olalde, Nick Patterson, and David Reich (2024). “The Allen Ancient DNA Resource (AADR) a curated compendium of ancient human genomes”. Scientific Data 11(1), p. 182.

Mathews, John and Kurtis Fink (2003). Numerical Methods Using Matlab. 4th edition. Upper Saddle River, N.J: Pearson.

Mathieson, Iain and Gil McVean (2013). “Estimating Selection Coefficients in Spatially Structured Populations from Time Series Data of Allele Frequencies”. Genetics 193(3), pp. 973–984.

Mathieson, Iain and Jonathan Terhorst (2022). “Direct Detection of Natural Selection in Bronze Age Britain”. Genome Research, gr.276862.122.

Mathieson, Iain et al. (2015). “Genome-Wide Patterns of Selection in 230 Ancient Eurasians”. Nature 528, pp. 499–503.

Moorjani, Priya, Sriram Sankararaman, Qiaomei Fu, Molly Przeworski, Nick Patterson, and David Reich (2016). “A Genetic Method for Dating Ancient Genomes Provides a Direct Estimate of Human Generation Interval in the Last 45,000 Years”. Proceedings of the National Academy of Sciences 113(20), pp. 5652–5657.

Nait Saada, Juba Georgios Kalantzis, Derek Shyr, Fergus Cooper, Martin Robinson, Alexander Gusev, and Pier Francesco Palamara (2020). “Identity-by-Descent Detection across 487,409 British Samples Reveals Fine Scale Population Structure and Ultra-Rare Variant Associations”. Nature Communications 11(1), p. 6130.

Nielsen, Rasmus (2005). “Molecular Signatures of Natural Selection”. Annual Review of Genetics 39(1), pp. 197–218.

Olalde, Iñigo et al. (2018). “The Beaker Phenomenon and the Genomic Transformation of Northwest Europe”. Nature 555, pp. 190–196.

Orlando, Ludovic, Robin Allaby, Pontus Skoglund, Clio Der Sarkissian, Philipp W. Stockhammer, María C. Ávila-Arcos, Qiaomei Fu, Johannes Krause, Eske Willerslev, Anne C. Stone, and Christina Warinner (2021). “Ancient DNA Analysis”. Nature Reviews Methods Primers 1(1), pp. 1–26.

Palmer, Duncan S., Wei Zhou, Liam Abbott, Emilie M. Wigdor, Nikolas Baya, Claire Churchhouse, Cotton Seed, Tim Poterba, Daniel King, Masahiro Kanai, Alex Bloemendal, and Benjamin M. Neale (2023). “Analysis of Genetic Dominance in the UK Biobank”. Science 379, pp. 1341–1348.

Paris, Cyriel, Bertrand Servin, and Simon Boitard (2019). “Inference of Selection from Genetic Time Series Using Various Parametric Approximations to the Wright-Fisher Model.” G3 9(12), pp. 4073–4086.

Patterson, Nick et al. (2022). “Large-Scale Migration into Britain during the Middle to Late Bronze Age”. Nature 601, pp. 588–594.

Peter, Benjamin M., Emilia Huerta-Sanchez, and Rasmus Nielsen (2012). “Distinguishing between Selective Sweeps from Standing Variation and from a De Novo Mutation”. PLOS Genetics 8(10), e1003011.

Purdue, Mark P. et al. (2009). “A Pooled Investigation of Toll-like Receptor Gene Variants and Risk of Non-Hodgkin Lymphoma”. Carcinogenesis 30(2), pp. 275–281.

Rabiner, L.R. (1989). “A Tutorial on Hidden Markov Models and Selected Applications in Speech Recognition”. Proceedings of the IEEE 77(2), pp. 257–286.

Rhodes, Shannon L., Daniel D. Buchanan, Ismaïl Ahmed, Kent D. Taylor, Marie-Anne Loriot, Janet S. Sinsheimer, Jeff M. Bronstein, Alexis Elbaz, George D. Mellick, Jerome I. Rotter, and Beate Ritz (2014). “Pooled Analysis of Iron-Related Genes in Parkinson’s Disease: Association with Transferrin”. Neurobiology of Disease 62, pp. 172–178.

Rieder, Stefan, Sead Taourit, Denis Mariat, Bertrand Langlois, and Gérard Guérin (2001). “Mutations in the Agouti (ASIP), the Extension (MC1R), and the Brown (TYRP1) Loci and Their Association to Coat Color Phenotypes in Horses (Equus Caballus)”. Mammalian Genome 12(6), pp. 450–455.

Sanjak, Jaleal S., Julia Sidorenko, Matthew R. Robinson, Kevin R. Thornton, and Peter M. Visscher (2018). “Evidence of directional and stabilizing selection in contemporary humans”. Proceedings of the National Academy of Sciences 115(1), pp. 151–156.

Sawyer, S. A. and D. L. Hartl (1992). “Population Genetics of Polymorphism and Divergence”. Genetics 132(4), pp. 1161–1176.

Schlötterer, C., R. Kofler, E. Versace, R. Tobler, and S. U. Franssen (2015). “Combining Experimental Evolution with Next-Generation Sequencing: A Powerful Tool to Study Adaptation from Standing Genetic Variation”. Heredity 114(5), pp. 431–440.

Schraiber, Joshua G., Steven N. Evans, and Montgomery Slatkin (2016). “Bayesian Inference of Natural Selection from Allele Frequency Time Series”. Genetics 203(1), pp. 493–511.

Simons, Yuval B., Kevin Bullaughey, Richard R. Hudson, and Guy Sella (2018). “A population genetic interpretation of GWAS findings for human quantitative traits”. PLOS Biology 16(3), e2002985.

Smith, John Maynard and John Haigh (1974). “The Hitch-Hiking Effect of a Favourable Gene”. Genetics Research 23(1), pp. 23–35.

Soejima, Mikiko and Yoshiro Koda (2007). “Population Differences of Two Coding SNPs in Pigmentation-Related Genes SLC24A5 and SLC45A2”. International Journal of Legal Medicine 121(1), pp. 36–39.

Steinrücken, Matthias, Anand Bhaskar, and Yun S. Song (2014). “A Novel Spectral Method for Inferring General Diploid Selection from Time Series Genetic Data”. The Annals of Applied Statistics 8(4), pp. 2203–2222.

Sun, Jielin, Fredrik Wiklund, Fang-Chi Hsu, Katarina Bälter, S. Lilly Zheng, Jan-Erik Johansson, Baoli Chang, Wennuan Liu, Tao Li, Aubrey R. Turner, Liwu Li, Ge Li, Hans-Olov Adami, William B. Isaacs, Jianfeng Xu, and Henrik Grönberg (2006). “Interactions of Sequence Variants in Interleukin-1 Receptor–Associated Kinase4 and the Toll-Like Receptor 6-1-10 Gene Cluster Increase Prostate Cancer Risk”. Cancer Epidemiology, Biomarkers & Prevention 15(3), pp. 480–485.

Swallow, Dallas M. (2003). “Genetics of Lactase Persistence and Lactose Intolerance”. Annual Review of Genetics 37, pp. 197–219.

Tataru, Paula, Maria Simonsen, Thomas Bataillon, and Asger Hobolth (2017). “Statistical Inference in the Wright–Fisher Model Using Allele Frequency Data”. Systematic Biology 66(1), e30–e46.

Taus, Thomas, Andreas Futschik, and Christian Schlötterer (2017). “Quantifying Selection with Pool-Seq Time Series Data”. Molecular Biology and Evolution 34(11), pp. 3023–3034.

Terhorst, Jonathan, Christian Schlötterer, and Yun S. Song (2015). “Multi-Locus Analysis of Genomic Time Series Data from Experimental Evolution”. PLOS Genetics 11(4), e1005069.

Varadhan, Ravi and Christophe Roland (2008). “Simple and Globally Convergent Methods for Accelerating the Convergence of Any EM Algorithm”. Scandinavian Journal of Statistics 35(2), pp. 335–353.

Vitti, Joseph J., Sharon R. Grossman, and Pardis C. Sabeti (2013). “Detecting Natural Selection in Genomic Data”. Annual Review of Genetics 47(1), pp. 97–120.

Vlachos, Christos, Claire Burny, Marta Pelizzola, Rui Borges, Andreas Futschik, Robert Kofler, and Christian Schlötterer (2019). “Benchmarking Software Tools for Detecting and Quantifying Selection in Evolve and Resequencing Studies”. Genome Biology 20(1), p. 169.

Wang, J. (2001). “A Pseudo-Likelihood Method for Estimating Effective Population Size from Temporally Spaced Samples”. Genetical Research 78(3), pp. 243–257.

Watterson, G. A. (1982). “Testing Selection at a Single Locus”. Biometrics 38(2), p. 323.

Wilks, S. S. (1938). “The Large-Sample Distribution of the Likelihood Ratio for Testing Composite Hypotheses”. The Annals of Mathematical Statistics 9(1), pp. 60–62.

Wong, Sunny H. et al. (2010). “Leprosy and the Adaptation of Human Toll-Like Receptor 1”. PLOS Pathogens 6(7), e1000979.

Wutke, Saskia, Norbert Benecke, Edson Sandoval-Castellanos, Hans-Jürgen Döhle, Susanne Friederich, Javier Gonzalez, Jón Hallsteinn Hallsson, Michael Hofreiter, Lembi Lõugas, Ola Magnell, Arturo Morales-Muniz, Ludovic Orlando, Albína Hulda Pálsdóttir, Monika Reissmann, Matej Ruttkay, Alexandra Trinks, and Arne Ludwig (2016). “Spotted Phenotypes in Horses Lost Attractiveness in the Middle Ages”. Scientific Reports 6(1), p. 38548.

