## Supplementary Material for "A novel expectation-maximization approach to infer general diploid selection from time-series genetic data"




#### S.1 Derivation of maximum-likelihood estimators

##### S.1.1 Maximum-likelihood estimator for additive selection

Here, we derive the maximum likelihood estimator for the selection coefficient $\hat{s}$ for a single bi-allelic locus in a constant-size haploid population following Watterson (1982). We present some details of this derivation to give appropriate context for the extension to the diploid model. In the additive model, we have $1+2s$, $1+s$, and 1 as the fitness values of the $AA$, $Aa$, and $aa$ genotype, respectively, for given selection coefficient $s$. The mean change in the $A$ allele frequency after selection is then given by

$$p' \approx p + sp(1-p).$$

As presented in Section 2.1 in the main text, the allele frequency dynamics in the Wright-Fisher model can be approximated by

$$p_{t+1} \sim \mathcal{N}\big(p'_t, \tfrac{1}{2N_e} p_t(1-p_t)\big),$$

where $N_e$ is the effective population size. Now, let $p_1, \ldots, p_T$ denote the exact frequency trajectory of the $A$ allele in the population. We can then write the likelihood defined in equation (1) in the main text as

$$L_s^{(C)}(p_1, \ldots, p_T) = \Big(\prod_{t=1}^{T-1} \frac{1}{\sqrt{2\pi\sigma_t^2}}\Big) \exp\Big(-\frac{1}{2}\sum_{t=1}^{T-1}\Big(\frac{p_{t+1} - p_t - sp_t(1-p_t)}{\sigma_t}\Big)^2\Big),$$

$$\ln L = \sum_{t=1}^{T-1}\Big(-\frac{1}{2}\ln(2\pi\sigma_t^2) - \frac{\big(p_{t+1} - p_t - sp_t(1-p_t)\big)^2}{2\sigma_t^2}\Big), \tag{S.1}$$

with $\sigma_t^2 = \frac{1}{2N_e} p_t(1-p_t)$. The maximum likelihood estimator $\hat{s}$ can be obtained by setting the derivative of equation (S.1) with respect to $s$ equal to zero. Omitting the terms $-\frac{1}{2}\ln(2\pi\sigma_t^2)$, since they do not depend on $s$, we have:

$$
\begin{aligned}
\frac{\partial \ln L}{\partial s} = 0 \quad & \Rightarrow \frac{\partial}{\partial s}\left(\sum_{t=1}^{T-1} -\frac{\big(p_{t+1} - p_t - s p_t(1-p_t)\big)^2}{2\sigma_t^2}\right) = 0 \\
& \Rightarrow \sum_{t=1}^{T-1} -\frac{1}{2\sigma_t^2}\frac{\partial}{\partial s}\big(p_{t+1} - p_t - s p_t(1-p_t)\big)^2 = 0 \\
& \Rightarrow \cancel{N_e} \sum_{t=1}^{T-1} \frac{1}{\cancel{-2p_t(1-p_t)}}\big(p_{t+1} - p_t - s p_t(1-p_t)\big) \cdot \cancel{\big(-2p_t(1-p_t)\big)} = 0 \\
& \Rightarrow \hat{s} = \frac{\sum_{t=1}^{T-1}(p_{t+1} - p_t)}{\sum_{t=1}^{T-1}\big(p_t(1-p_t)\big)} \\
& \Rightarrow \hat{s} = \frac{p_T - p_1}{\sum_{t=1}^{T-1} p_t(1-p_t)}.
\end{aligned}
$$

Thus, $\hat{s}$ is the total change in allele frequency divided by the sum of the allelic variance in each generation. The later is proportional to the expected allele frequency change in each generation, and the factor of proportionality is $s$.

##### S.1.2 Diploid maximum-likelihood estimates

In the full diploid model, selection is characterized by two parameters $s_1$ and $s_2$, as introduced in Section 2.1 in the main text. Then, the distribution of the allele frequency in the next generation $Y_{t+1}$ given the allele frequency in the current generation $Y_t = p_t$ can be approximated by a normal distribution with

$$Y_{t+1} \sim \mathcal{N}\big(p_t', \tfrac{1}{2N_e} p_t(1-p_t)\big),$$

where $p_t' := p_t + p_t(1-p_t)(s_1(1-2p_t) + s_2 p_t)$. The likelihood of a particular allele frequency trajectory $p_1, \ldots, p_T$ is then given by

$$L_{s_1,s_2}^{(C)}(p_1, \ldots, p_T) = \Big(\prod_{t=1}^{T-1} \frac{1}{\sqrt{2\pi\sigma_t^2}}\Big) \exp\Big(-\frac{1}{2}\sum_{t=1}^{T-1}\big(\frac{p_{t+1} - p_t'}{\sigma_t}\big)^2\Big),$$

and the log-likelihood by

$$\ln L_{s_1,s_2}^{(C)}(p_1, \ldots, p_T) = \sum_{t=1}^{T-1}\Big(-\frac{1}{2}\ln(2\pi\sigma_t^2) - \frac{(p_{t+1} - p_t')^2}{2\sigma_t^2}\Big), \qquad \text{(S.2)}$$

where again $\sigma_t^2 = \frac{1}{2N_e} p_t(1-p_t)$. The maximum likelihood estimators $\widehat{s_1}$ and $\widehat{s_2}$ can again be obtained by setting the derivative of equation (S.2) with respect to $s_1$ and $s_2$ equal to zero. Omitting the terms $-\frac{1}{2}\ln(2\pi\sigma_t^2)$, we have:

$$
\begin{aligned}
\frac{\partial \ln L}{\partial s_i} &= \frac{\partial}{\partial s_i}\left(\sum_{t=1}^{T-1} -\frac{(p_{t+1}-p_t')^2}{2\sigma_t^2}\right) \\
&= \sum_{t=1}^{T-1} -\frac{1}{2\sigma_t^2}\frac{\partial}{\partial s_i}\big(p_{t+1}-p_t-p_t(1-p_t)\big(s_1(1-2p_t)+s_2p_t\big)\big)^2 \stackrel{!}{=} 0
\end{aligned}
$$

Since $\frac{d}{dx}(a-bx)^2 = -2b(a-bx)$, we get the following two equations:

$$
\begin{aligned}
s_1\colon \quad & \sum_{t=1}^{T-1} \frac{1}{2\sigma_t^2} 2p_t(1-p_t)(1-2p_t)\big(p_{t+1}-p_t-p_t(1-p_t)\big(s_1(1-2p_t)+s_2p_t\big)\big) = 0, \\
s_2\colon \quad & \sum_{t=1}^{T-1} \frac{1}{2\sigma_t^2} 2p_t^2(1-p_t)\big(p_{t+1}-p_t-p_t(1-p_t)\big(s_1(1-2p_t)+s_2p_t\big)\big) = 0.
\end{aligned}
$$

Substituting $\sigma_t^2 = \frac{1}{2N_e}p_t(1-p_t)$, these expressions can be simplified to obtain:

$$
\begin{aligned}
& \sum_{t=1}^{T-1} N_e(1-2p_t)\big(p_{t+1}-p_t-p_t(1-p_t)\big(s_1(1-2p_t)+s_2p_t\big)\big) = 0, \\
& \sum_{t=1}^{T-1} N_e p_t\big(p_{t+1}-p_t-p_t(1-p_t)\big(s_1(1-2p_t)+s_2p_t\big)\big) = 0.
\end{aligned}
$$

Now, we collect terms proportional to $s_1$ and $s_2$ in both equations, to obtain

$$
\cancel{N_e}\Big(s_1 \underbrace{\sum_{t=1}^{T-1} p_tq_t(1-2p_t)^2}_{a} + s_2 \underbrace{\sum_{t=1}^{T-1} p_t^2q_t(1-2p_t)}_{b}\Big) = \cancel{N_e}\underbrace{\sum_{t=1}^{T-1}(1-2p_t)(p_{t+1}-p_t)}_{c}, \tag{S.3}
$$

$$
\cancel{N_e}\Big(s_1 \underbrace{\sum_{t=1}^{T-1} p_t^2q_t(1-2p_t)}_{b} + s_2 \underbrace{\sum_{t=1}^{T-1} p_t^3q_t}_{d}\Big) = \cancel{N_e}\underbrace{\sum_{t=1}^{T-1} p_t(p_{t+1}-p_t)}_{e}. \tag{S.4}
$$

with $q_t = 1-p_t$. The MLEs $\widehat{s_1}$ and $\widehat{s_2}$, can now be obtained as the solutions of the linear system

$$
\begin{cases} as_1 + bs_2 = c \\ bs_1 + ds_2 = e. \end{cases}
$$

This system can be solved using Gaussian elimination as follows:

$$
\begin{cases} as_1 + bs_2 = c \\ bs_1 + ds_2 = e \end{cases} \Rightarrow \begin{cases} ads_1 + bds_2 = cd \\ b^2s_1 + bds_2 = eb \end{cases}
$$

$$
\Rightarrow (ad-b^2)s_1 = cd - eb \Rightarrow \boxed{s_1 = \frac{cd-be}{ad-b^2}}.
$$

Similarly, we get:

$$s_2 = \frac{c}{b} - \frac{a}{b} s_1 => s_2 = \frac{c}{b} - \frac{acd - abe}{b(ad - b^2)}$$

$$=> s_2 = \frac{acd - b^2 c - (acd - abe)}{abd - b^3} => \boxed{s_2 = \frac{ae - bc}{ad - b^2}}.$$

Substituting the expressions from equation (S.3) and equation (S.4) into the above solution yields

$$\widehat{s_1} = \frac{\sum_t (1-2p_t)(p_{t+1}-p_t)\sum_t p_t^3 q_t - \sum_t p_t^2 q_t (1-2p_t)\sum_t p_t(p_{t+1}-p_t)}{\sum_t p_t q_t (1-2p_t)^2 \sum_t p_t^3 q_t - \{\sum_t p_t^2 q_t (1-2p_t)\}^2}, \tag{S.5}$$

and

$$\widehat{s_2} = \frac{\sum_t p_t q_t (1-2p_t)^2 \sum_t p_t(p_{t+1}-p_t) - \sum_t p_t^2 q_t (1-2p_t)\sum_t (1-2p_t)(p_{t+1}-p_t)}{\sum_t p_t q_t (1-2p_t)^2 \sum_t p_t^3 q_t - \{\sum_t p_t^2 q_t (1-2p_t)\}^2}. \tag{S.6}$$

To simplify these expression, we simplify the numerators and denominator separately, and obtain:

$$\begin{aligned}
&\text{Numerator of (S.5)}\\
&= \sum_t (1-2p_t)(p_{t+1}-p_t)\sum_t p_t^3 q_t - \sum_t p_t^2 q_t (1-2p_t)\sum_t p_t(p_{t+1}-p_t)\\
&= \sum_t (p_{t+1}-p_t-2p_t(p_{t+1}-p_t))\sum_t p_t^3 q_t - \sum_t (p_t^2 q_t - 2p_t^3 q_t)\sum_t p_t(p_{t+1}-p_t)\\
&= \sum_t (p_{t+1}-p_t)\sum_t p_t^3 q_t - 2\sum_t p_t(p_{t+1}-p_t)\sum_t p_t^3 q_t\\
&\qquad - \sum_t p_t(p_{t+1}-p_t)\sum_t p_t^2 q_t + 2\sum_t p_t^3 q_t \sum_t p_t(p_{t+1}-p_t),
\end{aligned}$$

Recognizing that the telescopic sum $\sum_t (p_{t+1}-p_t)$ equals $p_T - p_1$, we get

$$\text{Numerator of (S.5)} = (p_T - p_1)\sum_t p_t^3 q_t - \sum_t p_t(p_{t+1}-p_t)\sum_t p_t^2 q_t.$$

Similarly, we can simplify the numerator of equation (S.6), and arrive at

$$\begin{aligned}
&\text{Numerator of (S.6)}\\
&= \sum_t p_t q_t (1-2p_t)^2 \sum_t p_t(p_{t+1}-p_t) - \sum_t p_t^2 q_t (1-2p_t)\sum_t (1-2p_t)(p_{t+1}-p_t)\\
&= \sum_t (p_t q_t - 4p_t^2 q_t + 4p_t^3 q_t)\sum_t p_t(p_{t+1}-p_t)\\
&\qquad - \sum_t (p_t^2 q_t - 2p_t^3 q_t)\sum_t (p_{t+1}-p_t) + \sum_t (2p_t^2 q_t - 4p_t^3 q_t)\sum_t p_t(p_{t+1}-p_t)\\
&= \sum_t (p_t q_t - 2p_t^2 q_t)\sum_t p_t(p_{t+1}-p_t) - (p_T - p_1)\sum_t (p_t^2 q_t - 2p_t^3 q_t)\\
&= \sum_t p_t q_t (1-2p_t)\sum_t p_t(p_{t+1}-p_t) - (p_T - p_1)\sum_t p_t^2 q_t (1-2p_t).
\end{aligned}$$

Lastly, simplifying the denominator of both expressions yields

$$
\begin{aligned}
&\text{Denominator of (S.5) and (S.6)}\\
&= \sum_t p_t q_t (1-2p_t)^2 \sum_t p_t^3 q_t - \Big(\sum_t p_t^2 q_t (1-2p_t)\Big)^2\\
&= \sum_t (p_t q_t - 4p_t^2 q_t + 4p_t^3 q_t) \sum_t p_t^3 q_t - \Big(\sum_t (p_t^2 q_t - 2p_t^3 q_t)\Big)^2\\
&= \sum_t (p_t q_t - \cancel{4p_t^2 q_t} + \cancel{4p_t^3 q_t}) \sum_t p_t^3 q_t - \sum_t p_t^2 q_t \sum_t p_t^2 q_t + \cancel{4\sum_t p_t^2 q_t \sum_t p_t^3 q_t} - \cancel{4\sum_t p_t^3 q_t \sum_t p_t^3 q_t}\\
&= \sum_t p_t q_t \sum_t p_t^3 q_t - \Big(\sum_t p_t^2 q_t\Big)^2
\end{aligned}
$$

Combining these expressions, the maximum likelihood estimators of $s_1$ and $s_2$ are:

$$
\begin{aligned}
\widehat{s_1} &= \frac{(p_T - p_1)\sum_t p_t^3 q_t - \sum_t p_t(p_{t+1}-p_t)\sum_t p_t^2 q_t}{\sum_t p_t q_t \sum_t p_t^3 q_t - \big(\sum_t p_t^2 q_t\big)^2}\\
\widehat{s_2} &= \frac{\sum_t p_t q_t (1-2p_t)\sum_t p_t(p_{t+1}-p_t) - (p_T - p_1)\sum_t p_t^2 q_t (1-2p_t)}{\sum_t p_t q_t \sum_t p_t^3 q_t - \big(\sum_t p_t^2 q_t\big)^2}.
\end{aligned}
$$

Note that we assumed a constant population size for simplicity. However, this is not restrictive, and these estimators could readily be extended to scenarios with varying population size.

#### S.2 Maximization-step

##### S.2.1 Additive selection

To derive the update in the M-step for the HMM if selection is *additive*, we proceed similar to Section S.1.1. We determine the maximum by setting the derivative of the conditional log-likelihood to zero and solving for $s$. Again, we use the shorthand notation

$$
\mathbb{E}[\,\cdot\,] := \mathbb{E}_{s^{(k)}}\big[\,\cdot\,\big|O_t = o_t\,\forall\, 1 \le t \le T\big]
$$

for the conditional expectation, and obtain

$$
\mathbb{E}\big[\ln L_s^{(C)}(F_1,\ldots,F_T)\big] = \mathbb{E}\Big[\sum_{t=1}^{T-1}\Big(-\frac{1}{2}\ln(2\pi\sigma_t^2) - \frac{\big(F_{t+1} - F_t - s^{(k)}F_t(1-F_t)\big)^2}{2\sigma_t^2}\Big)\Big],
$$

with $\sigma_t^2 := \frac{1}{2N_e}F_t(1-F_t)$. Note that the conditional expectation is a function of the current parameter estimate $s^{(k)}$, but not of the parameter $s$ that we maximize over. Thus, the order of the expected value and the derivative can be exchanged, and we arrive at

$$
-\sum_{t=1}^{T-1}\mathbb{E}\Big[\frac{1}{2\sigma_t^2}\frac{\partial}{\partial s}\big(F_{t+1} - F_t - sF_t(1-F_t)\big)^2\Big] = 0,
$$

which, similar to Section S.1.1, yields

$$
s^{(k+1)} = \frac{\mathbb{E}[F_T] - \mathbb{E}[F_1]}{\sum_{t=1}^{T-1}\mathbb{E}[F_t(1-F_t)]}.
$$

This expression is the same as Equation (18) by Mathieson and McVean 2013.

##### S.2.2 General diploid selection

To derive the parameter update for the M-step in the diploid case given in equation (2) in the main text, we again set the derivative of the conditional log-likelihood with respect to the parameters equal to zero. The conditional log-likelihood is given by

$$\begin{aligned}\mathbb{E}\left[\ln L^{(C)}_{s_1,s_2}(F_1,\dots,F_T)\right] \\ = -\frac{1}{2}\sum_t \mathbb{E}\left[\frac{(F_{t+1}-F_t-s_1F_t(1-F_t)(1-2F_t)-s_2F_t^2(1-F_t)^2}{\sigma_t^2}\right],\end{aligned} \tag{S.7}$$

where again

$$\mathbb{E}[\,\cdot\,] := \mathbb{E}_{s_1^{(k)},s_2^{(k)}}\left[\,\cdot\,\middle|O_t = o_t\,\forall\, 1\le t\le T\right].$$

Taking the derivative of expression (S.7) with respect to $s_1$ and $s_2$, exchanging the order of the expectation and the derivative, and equating it to zero yields the linear system

$$\begin{aligned}s_1\sum_{t=1}^{T-1}\mathbb{E}\left[F_t(1-F_t)(1-2F_t)^2\right] + s_2\sum_{t=1}^{T-1}\mathbb{E}\left[F_t^2(1-F_t)(1-2F_t)\right] \\ = \sum_{t=1}^{T-1}\mathbb{E}\left[(1-2F_t)(F_{t+1}-F_t)\right],\end{aligned} \tag{S.8}$$

$$s_1\sum_{t=1}^{T-1}\mathbb{E}\,F_t^2(1-F_t)(1-2F_t)\big] + s_2\sum_{t=1}^{T-1}\left[F_t^3(1-F_t)\right] = \sum_{t=1}^{T-1}\mathbb{E}\left[F_t(F_{t+1}-F_t)\right], \tag{S.9}$$

as in equations (S.3) and (S.4). Similar to Section S.1.2, the solutions of this system, and thus the updates for the M-step, are given by

$$s_1^{(k+1)} = \frac{\big(\mathbb{E}[F_T]-\mathbb{E}[F_1]\big)\sum_t\mathbb{E}\left[F_t^3(1-F_t)\right]-\sum_t\mathbb{E}\left[F_t(F_{t+1}-F_t)\right]\sum_t\mathbb{E}\left[F_t^2(1-F_t)\right]}{\sum_t\mathbb{E}\left[F_t(1-F_t)\right]\sum_t\mathbb{E}\left[F_t^3(1-F_t)\right]-\big(\sum_t\mathbb{E}\left[F_t^2(1-F_t)\right]\big)^2},$$

$$s_2^{(k+1)} = \frac{\sum_t\mathbb{E}\left[F_t(1-F_t)(1-2F_t)\right]\sum_t\mathbb{E}\left[F_t(F_{t+1}-F_t)\right]-\big(\mathbb{E}[F_T]-\mathbb{E}[F_1]\big)\sum_t\mathbb{E}\left[F_t^2(1-F_t)(1-2F_t)\right]}{\sum_t\mathbb{E}\left[F_t(1-F_t)\right]\sum_t\mathbb{E}\left[F_t^3(1-F_t)\right]-\big(\sum_t\mathbb{E}\left[F_t^2(1-F_t)\right]\big)^2}.$$

##### S.2.3 Linear constraints in Lagrange multiplier formalism

In Section 2.2.2 in the main text, we introduced the formalism of Lagrange multipliers to obtain an MLE estimator in the general diploid HMM framework under linear constraints. To this end, we have to solve the system of equations

$$\begin{aligned}\frac{\partial}{\partial s_1}\mathbb{E}\left[\ln L^{(C)}_{s_1,s_2}(F_1,\dots,F_T)\right] &= \lambda\frac{\partial}{\partial s_1}g(s_1,s_2)\\ \frac{\partial}{\partial s_2}\mathbb{E}\left[\ln L^{(C)}_{s_1,s_2}(F_1,\dots,F_T)\right] &= \lambda\frac{\partial}{\partial s_2}g(s_1,s_2)\\ as_1-bs_2 &= 0,\end{aligned}$$

where $g(s_1, s_2) = as_1 - bs_2$. The expectation is again over the posterior distribution of $F_1, \ldots, F_T$, conditional on the observed data and current values $s_1^{(k)}$, $s_2^{(k)}$. Similar to equations (S.8) and (S.9), we obtain

$$2N_e \sum_{t=1}^{T-1} \mathbb{E}\left[(F_{t+1} - F_t)(1 - 2F_t)\right] - s_1 \mathbb{E}\left[F_t(1 - F_t)(1 - 2F_t)^2\right] - s_2 \mathbb{E}\left[F_t^2(1 - F_t)(1 - 2F_t)\right] = \lambda a,$$

$$2N_e \sum_{t=1}^{T-1} \mathbb{E}\left[(F_{t+1} - F_t)F_t\right] - s_1 \mathbb{E}\left[F_t^2(1 - F_t)(1 - 2F_t)\right] - s_2 \mathbb{E}\left[F_t^3(1 - F_t)\right] = -\lambda b,$$

for the first two equations. We can then multiply the first equation by $b$, the second by $a$, and then sum to remove $\lambda$, and obtain

$$\begin{aligned} 2N_e \sum_{t=1}^{T-1} b\,\mathbb{E}\left[(F_{t+1} - F_t)(1 - 2F_t)\right] - bs_1 \mathbb{E}\left[F_t(1 - F_t)(1 - 2F_t)^2\right] - bs_2 \mathbb{E}\left[F_t^2(1 - F_t)(1 - 2F_t)\right] \\ + a\,\mathbb{E}\left[(F_{t+1} - F_t)F_t\right] - as_1 \mathbb{E}\left[F_t^2(1 - F_t)(1 - 2F_t)\right] - as_2 \mathbb{E}\left[F_t^3(1 - F_t)\right] = 0. \end{aligned}$$

Since $as_1 = bs_2$, we can multiply by $a$ and substitute $s_1 = \frac{b}{a}s_2$. Then, rearranging, we have

$$\begin{aligned} &\sum_{t=1}^{T-1} ab\,\mathbb{E}\left[(F_{t+1} - F_t)(1 - 2F_t)\right] + a^2\,\mathbb{E}\left[(F_{t+1} - F_t)F_t\right] \\ &= s_2 \sum_{t=1}^{T-1} b^2\,\mathbb{E}\left[F_t(1-F_t)(1-2F_t)^2\right] + ab\,\mathbb{E}\left[F_t^2(1-F_t)(1-2F_t)\right] + ab\,\mathbb{E}\left[F_t^2(1-F_t)(1-2F_t)\right] + a^2\,\mathbb{E}\left[F_t^3(1-F_t)\right] \end{aligned}$$

Combining terms and solving for $s_2$ yields

$$s_2 = \frac{a \sum_{t=1}^{T-1} b\,\mathbb{E}\left[(F_{t+1} - F_t)(1 - 2F_t)\right] + a\,\mathbb{E}\left[(F_{t+1} - F_t)F_t\right]}{\sum_{t=1}^{T-1} \mathbb{E}\left[F_t(1 - F_t)\big(aF_t + b(1 - 2F_t)\big)^2\right]},$$

and, since $s_1 = \frac{b}{a}s_2$,

$$s_1 = \frac{b \sum_{t=1}^{T-1} b\,\mathbb{E}\left[(F_{t+1} - F_t)(1 - 2F_t)\right] + a\,\mathbb{E}\left[(F_{t+1} - F_t)F_t\right]}{\sum_{t=1}^{T-1} \mathbb{E}\left[F_t(1 - F_t)\big(aF_t + b(1 - 2F_t)\big)^2\right]}.$$

For *additive*, *dominant*, *recessive*, and *heterozygote difference* selection, we have $(a, b) = (2, 1)$, $(1, 1)$, $(1, 0)$, and $(0, 1)$, respectively. Note that we do not need to obtain an explicit solution for $\lambda$, since the value is not used in the EM update.

#### S.3 Boxplots, Q-Q plots, and AUC tables for additional simulation parameters

Figure S.1 to Figure S.8 in this Section show the results obtained from simulations similar to Figure 3 in the main text under different parameter combinations for number of generations $T \in \{101, 251, 1001\}$ and the initial condition fixed frequency at $p \in \{0.005, 0.25\}$ or standing variation. Note that *recessive* selection is omitted for the initial conditions fixed frequency at $p = 0.005$ and standing variation, since the selected allele is often lost, resulting in inaccurate estimates. Similarly, *underdominance* is omitted for the initial condition fixed frequency at $p = 0.005$.

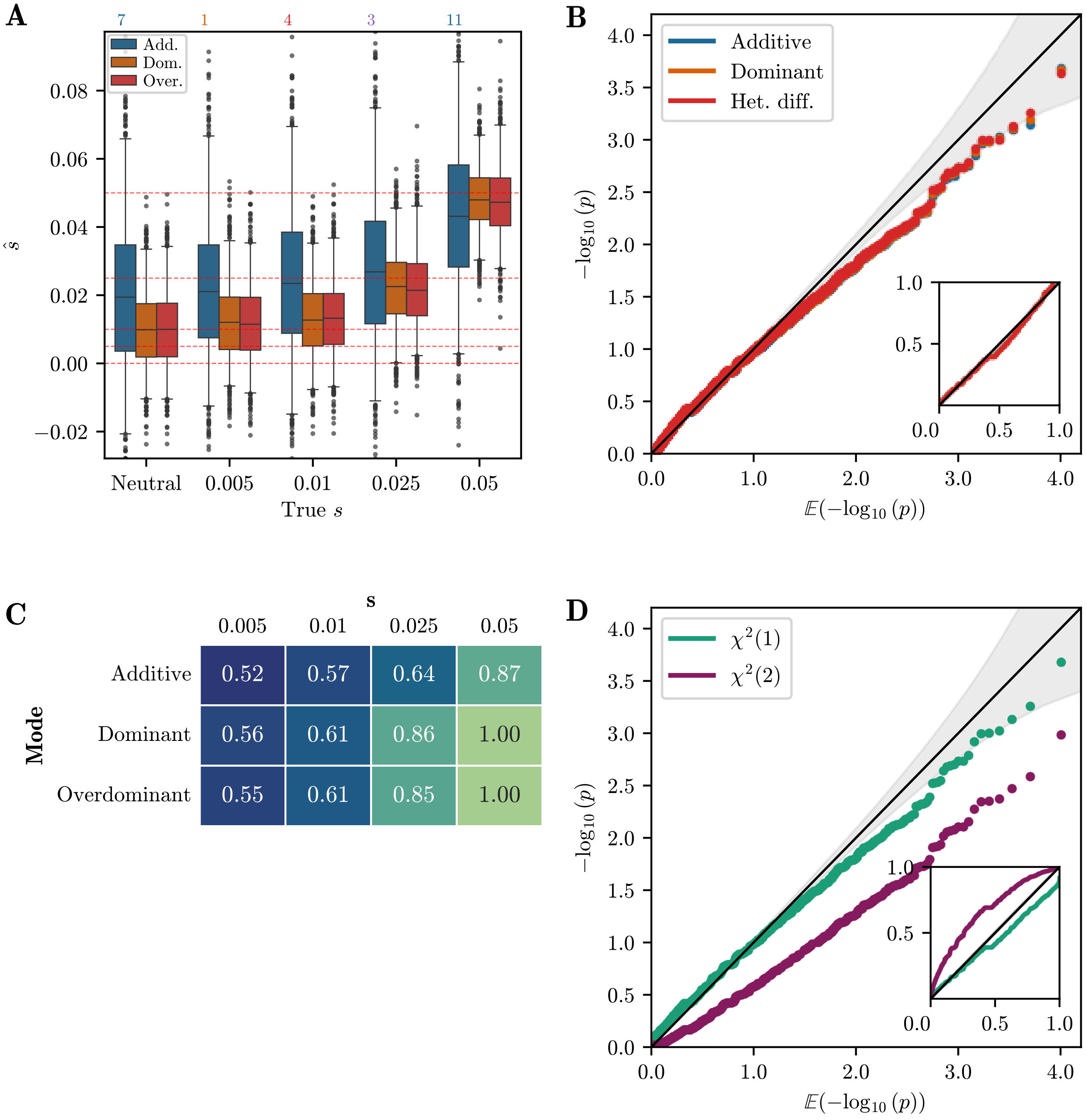


**Figure S.1:** A) Boxplot of $\hat{s}$ for 1,000 replicates simulated under each selection mode. Whiskers extend to 2.5% and 97.5%-tiles. Number of estimates outside plotting range indicated above the plot. B) Q-Q plot of $-\log_{10}$(p) against $\mathbb{E}[-\log_{10}(\text{p})]$ of single-alternative tests for neutral replicates. Inset shows same plot for raw values. C) Table of AUC values based on likelihood-ratios for each selection mode and selection strength simulated. D) Q-Q plot of $-\log_{10}$(p) against $\mathbb{E}[-\log_{10}(\text{p})]$ for the $\delta$ statistic, assuming different distributions. For all simulations, the number of generations is $T = 101$ and the initial condition is fixed at frequency $p = 0.005$.

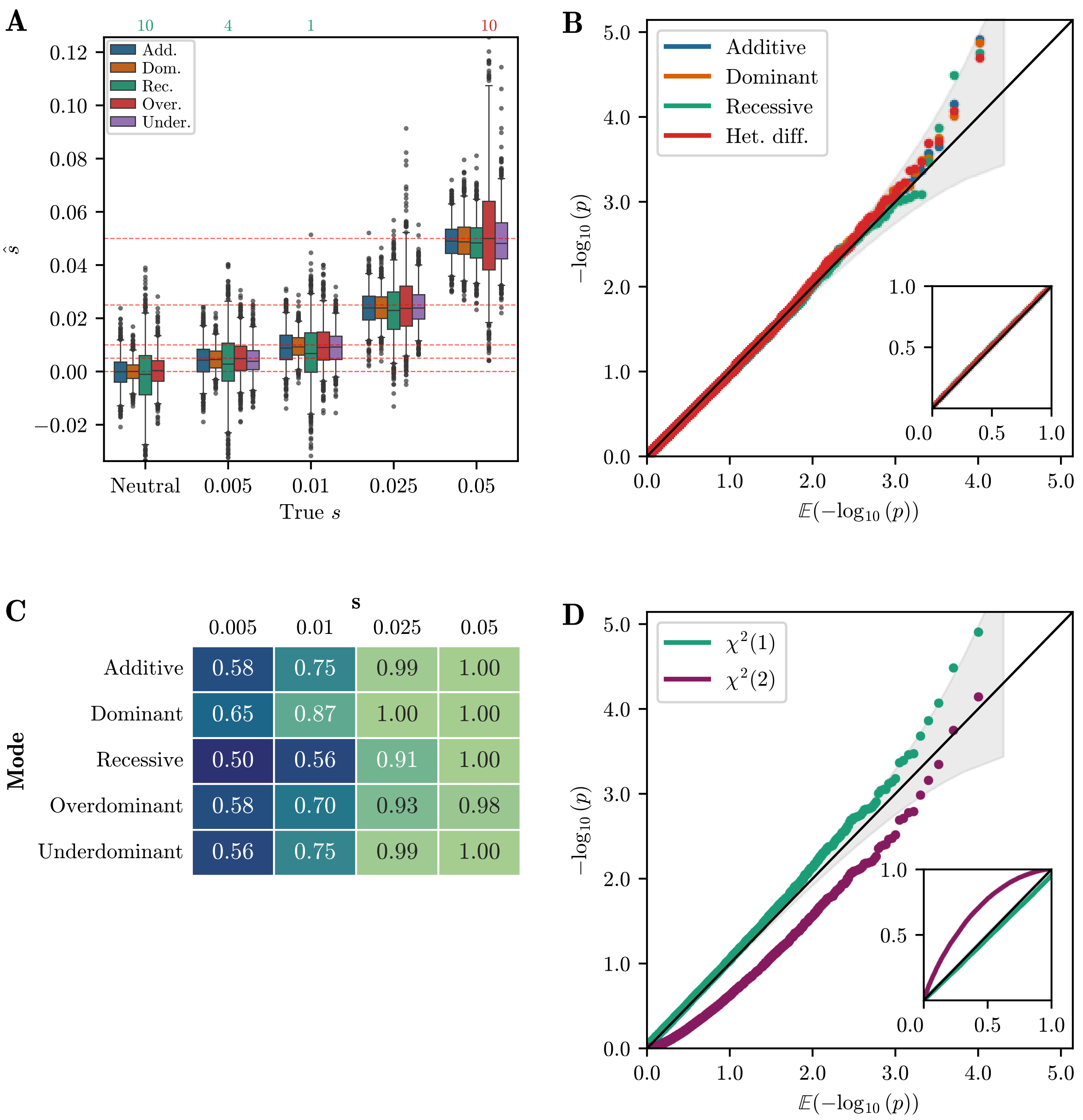


**Figure S.2:** A) Boxplot of $\hat{s}$ for 1,000 replicates simulated under each selection mode. Whiskers extend to 2.5% and 97.5%-tiles. Number of estimates outside plotting range indicated above the plot. B) Q-Q plot of $-\log_{10}(\mathrm{p})$ against $\mathbb{E}[-\log_{10}(\mathrm{p})]$ of single-alternative tests for neutral replicates. Inset shows same plot for raw values. C) Table of AUC values based on likelihood-ratios for each selection mode and selection strength simulated. D) Q-Q plot of $-\log_{10}(\mathrm{p})$ against $\mathbb{E}[-\log_{10}(\mathrm{p})]$ for the $\delta$ statistic, assuming different distributions. For all simulations, the number of generations is $T = 101$ and the initial condition is fixed at frequency $p = 0.25$.

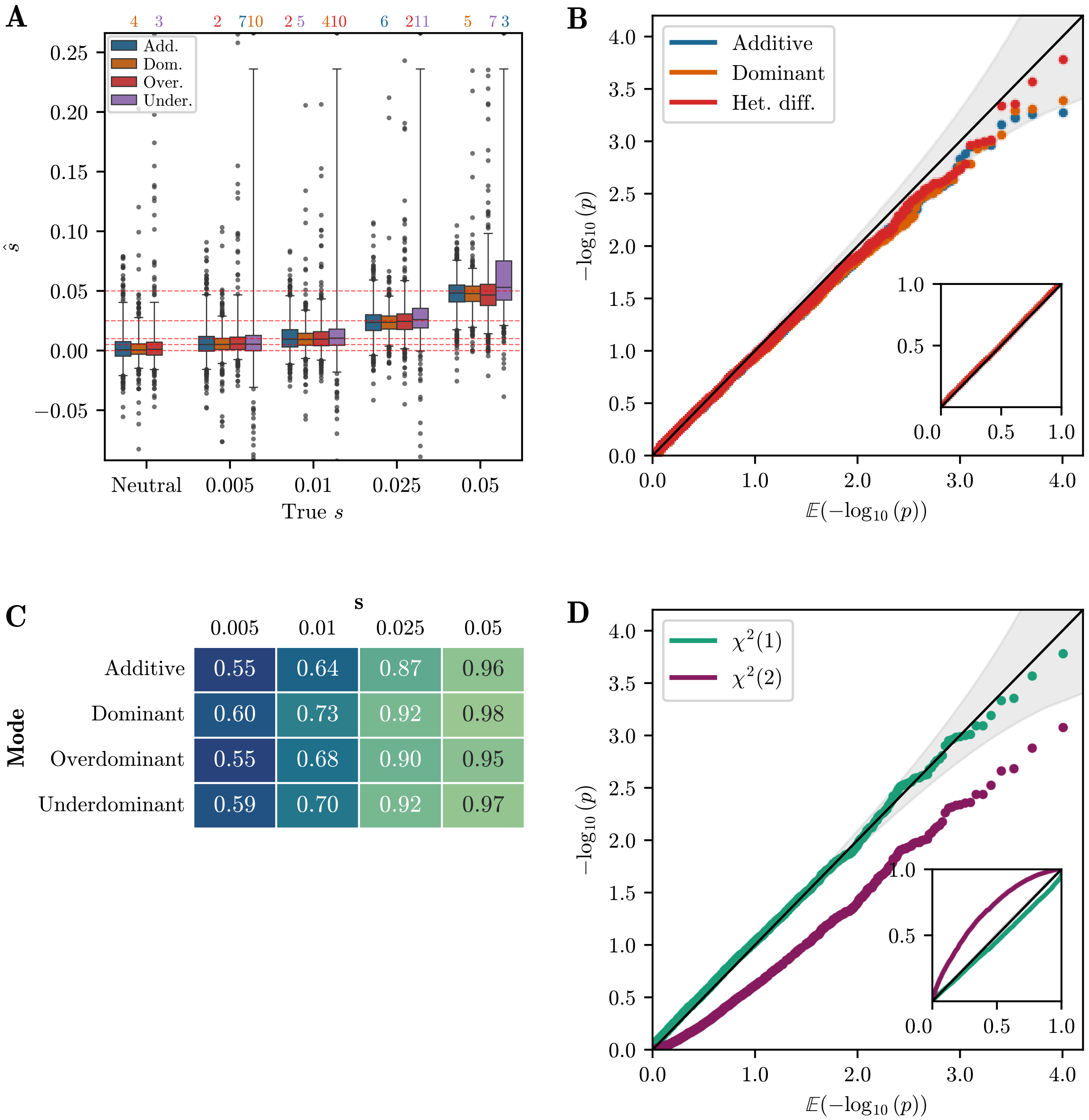


**Figure S.3:** A) Boxplot of $\hat{s}$ for 1,000 replicates simulated under each selection mode. Whiskers extend to 2.5% and 97.5%-tiles. Number of estimates outside plotting range indicated above the plot. B) Q-Q plot of $-\log_{10}(\text{p})$ against $\mathbb{E}[-\log_{10}(\text{p})]$ of single-alternative tests for neutral replicates. Inset shows same plot for raw values. C) Table of AUC values based on likelihood-ratios for each selection mode and selection strength simulated. D) Q-Q plot of $-\log_{10}(\text{p})$ against $\mathbb{E}[-\log_{10}(\text{p})]$ for the $\delta$ statistic, assuming different distributions. For all simulations, the number of generations is $T = 101$ and the initial condition is standing variation.

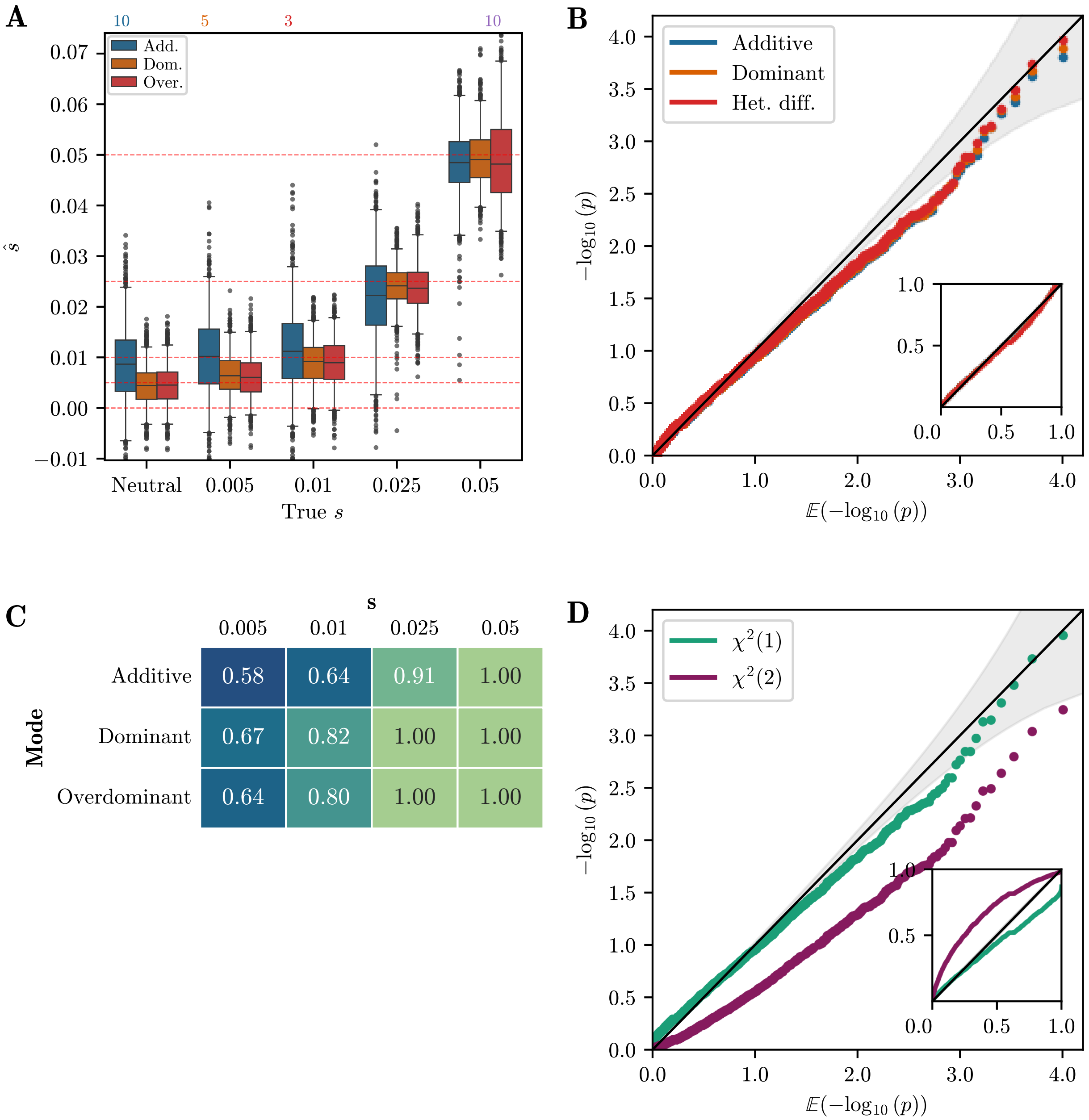


**Figure S.4:** A) Boxplot of $\hat{s}$ for 1,000 replicates simulated under each selection mode. Whiskers extend to 2.5% and 97.5%-tiles. Number of estimates outside plotting range indicated above the plot. B) Q-Q plot of $-\log_{10}(\text{p})$ against $\mathbb{E}[-\log_{10}(\text{p})]$ of single-alternative tests for neutral replicates. Inset shows same plot for raw values. C) Table of AUC values based on likelihood-ratios for each selection mode and selection strength simulated. D) Q-Q plot of $-\log_{10}(\text{p})$ against $\mathbb{E}[-\log_{10}(\text{p})]$ for the $\delta$ statistic, assuming different distributions. For all simulations, the number of generations is $T = 251$ and the initial condition is fixed at frequency $p = 0.005$.

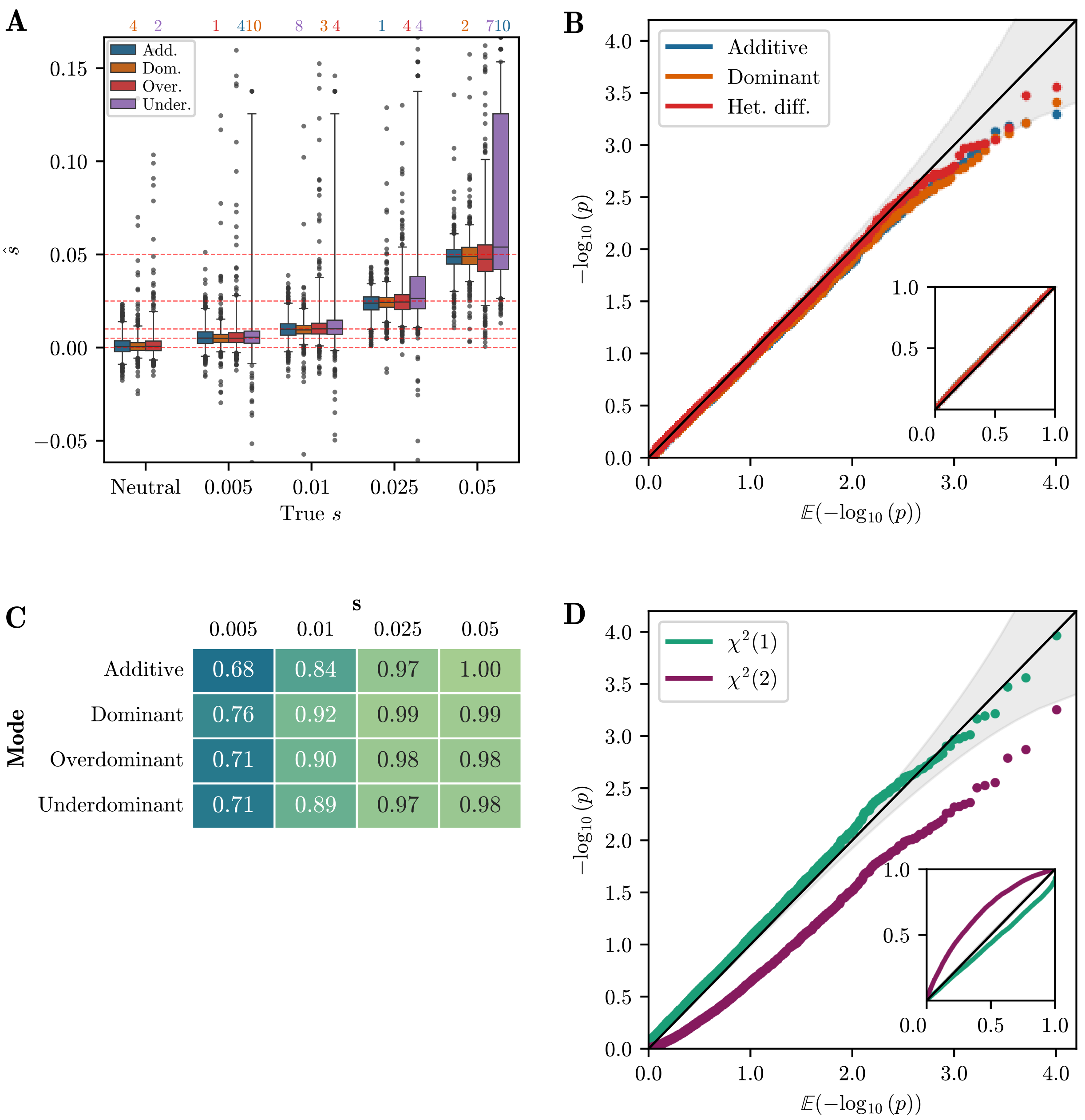


**Figure S.5:** A) Boxplot of $\hat{s}$ for 1,000 replicates simulated under each selection mode. Whiskers extend to 2.5% and 97.5%-tiles. Number of estimates outside plotting range indicated above the plot. B) Q-Q plot of $-\log_{10}(\text{p})$ against $\mathbb{E}[-\log_{10}(\text{p})]$ of single-alternative tests for neutral replicates. Inset shows same plot for raw values. C) Table of AUC values based on likelihood-ratios for each selection mode and selection strength simulated. D) Q-Q plot of $-\log_{10}(\text{p})$ against $\mathbb{E}[-\log_{10}(\text{p})]$ for the $\delta$ statistic, assuming different distributions. For all simulations, the number of generations is $T = 251$ and the initial condition is standing variation.

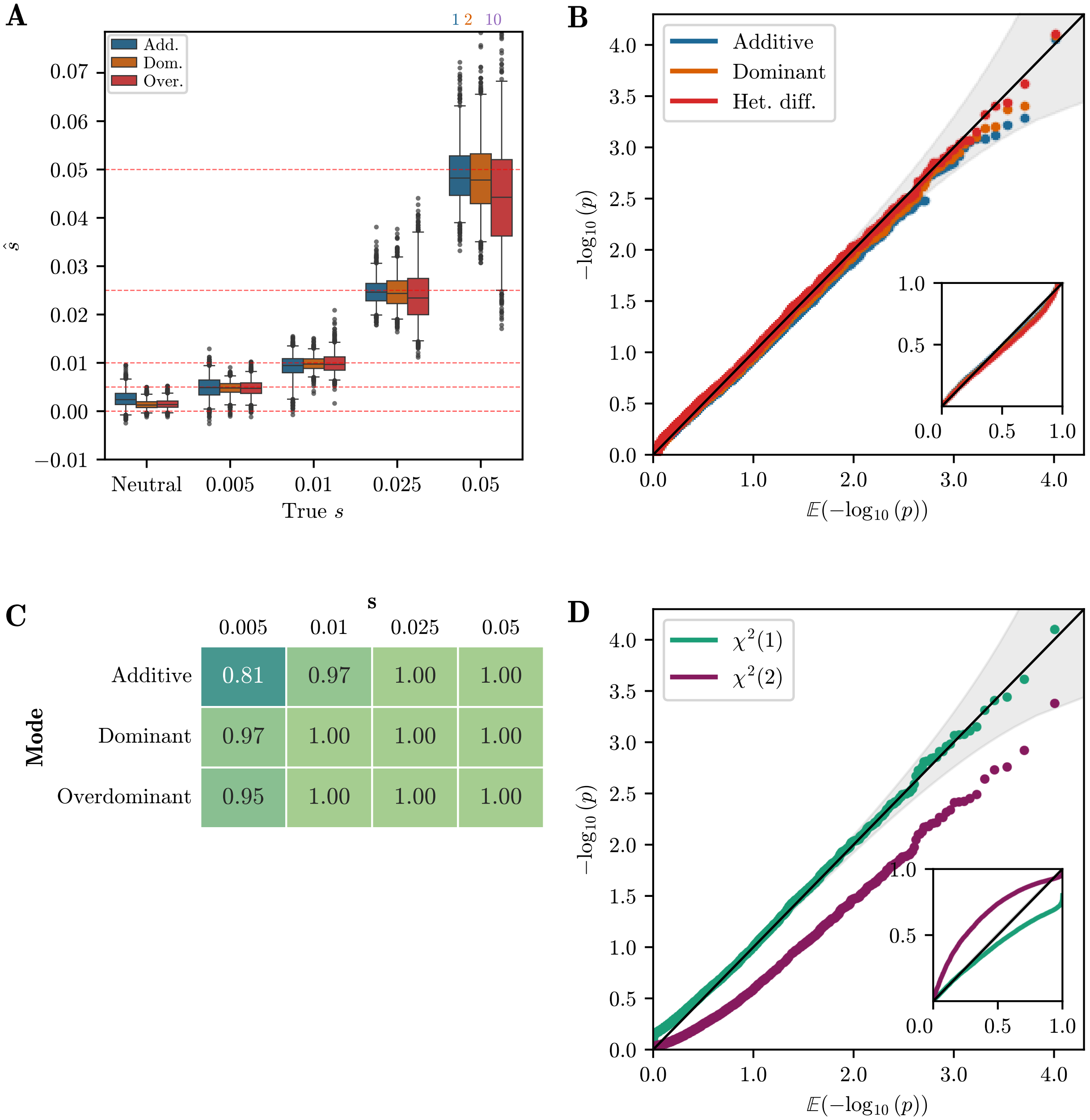


**Figure S.6:** A) Boxplot of $\hat{s}$ for 1,000 replicates simulated under each selection mode. Whiskers extend to 2.5% and 97.5%-tiles. Number of estimates outside plotting range indicated above the plot. B) Q-Q plot of $-\log_{10}(\text{p})$ against $\mathbb{E}[-\log_{10}(\text{p})]$ of single-alternative tests for neutral replicates. Inset shows same plot for raw values. C) Table of AUC values based on likelihood-ratios for each selection mode and selection strength simulated. D) Q-Q plot of $-\log_{10}(\text{p})$ against $\mathbb{E}[-\log_{10}(\text{p})]$ for the $\delta$ statistic, assuming different distributions. For all simulations, the number of generations is $T = 1001$ and the initial condition is fixed at frequency $p = 0.005$.

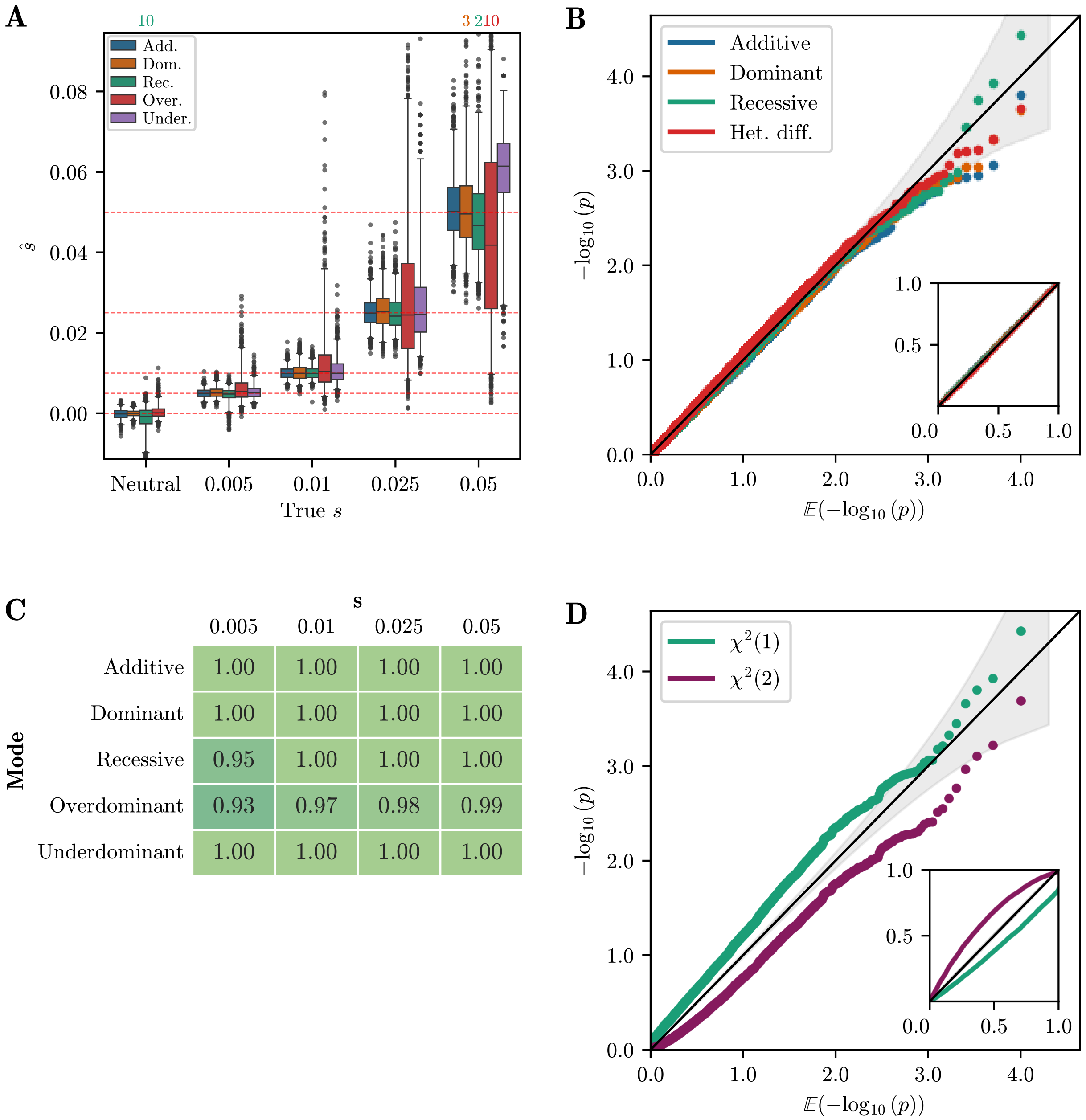


| Mode | 0.005 | 0.01 | 0.025 | 0.05 |
|---|---|---|---|---|
| Additive | 1.00 | 1.00 | 1.00 | 1.00 |
| Dominant | 1.00 | 1.00 | 1.00 | 1.00 |
| Recessive | 0.95 | 1.00 | 1.00 | 1.00 |
| Overdominant | 0.93 | 0.97 | 0.98 | 0.99 |
| Underdominant | 1.00 | 1.00 | 1.00 | 1.00 |

**Figure S.7:** A) Boxplot of $\hat{s}$ for 1,000 replicates simulated under each selection mode. Whiskers extend to 2.5% and 97.5%-tiles. Number of estimates outside plotting range indicated above the plot. B) Q-Q plot of $-\log_{10}(\mathrm{p})$ against $\mathbb{E}[-\log_{10}(\mathrm{p})]$ of single-alternative tests for neutral replicates. Inset shows same plot for raw values. C) Table of AUC values based on likelihood-ratios for each selection mode and selection strength simulated. D) Q-Q plot of $-\log_{10}(\mathrm{p})$ against $\mathbb{E}[-\log_{10}(\mathrm{p})]$ for the $\delta$ statistic, assuming different distributions. For all simulations, the number of generations is $T = 1001$ and the initial condition is fixed at frequency $p = 0.25$.

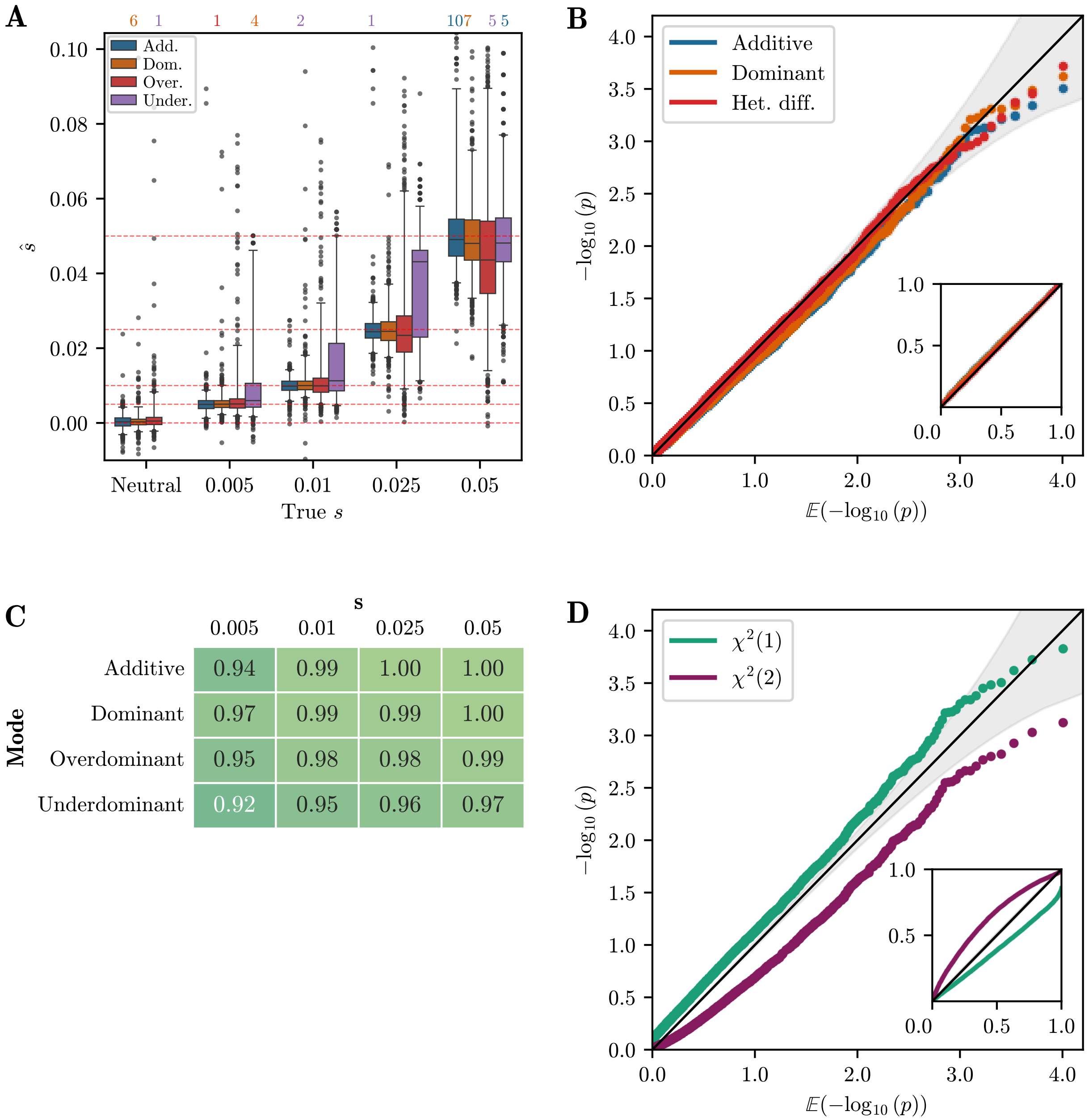


**Figure S.8:** A) Boxplot of $\hat{s}$ for 1,000 replicates simulated under each selection mode. Whiskers extend to 2.5% and 97.5%-tiles. Number of estimates outside plotting range indicated above the plot. B) Q-Q plot of $-\log_{10}(\text{p})$ against $\mathbb{E}[-\log_{10}(\text{p})]$ of single-alternative tests for neutral replicates. Inset shows same plot for raw values. C) Table of AUC values based on likelihood-ratios for each selection mode and selection strength simulated. D) Q-Q plot of $-\log_{10}(\text{p})$ against $\mathbb{E}[-\log_{10}(\text{p})]$ for the $\delta$ statistic, assuming different distributions. For all simulations, the number of generations is $T = 1001$ and the initial condition is standing variation.

#### S.4 Unconstrained EM estimation accuracy

In this section, we investigate the accuracy of the unconstrained HMM-EM in estimating selection coefficients for each mode. Figure S.9 shows a scatterplot of the unconstrained MLE $(\hat{s}_1, \hat{s}_2)$ estimates for the data described in Section 3.1.1 in the main text, simulated with $N_e = 10{,}000$, the number of generations $T = 251$, and a fixed initial frequency of $p = 0.25$. We plot the estimates for all modes of selection with $s = 0.05$. We find that estimation is accurate for *recessive*, *additive*, *dominant*, and *overdominant* selection. For *underdominant* selection, $s_1$ is estimated accurately but $s_2$ is poorly estimated.

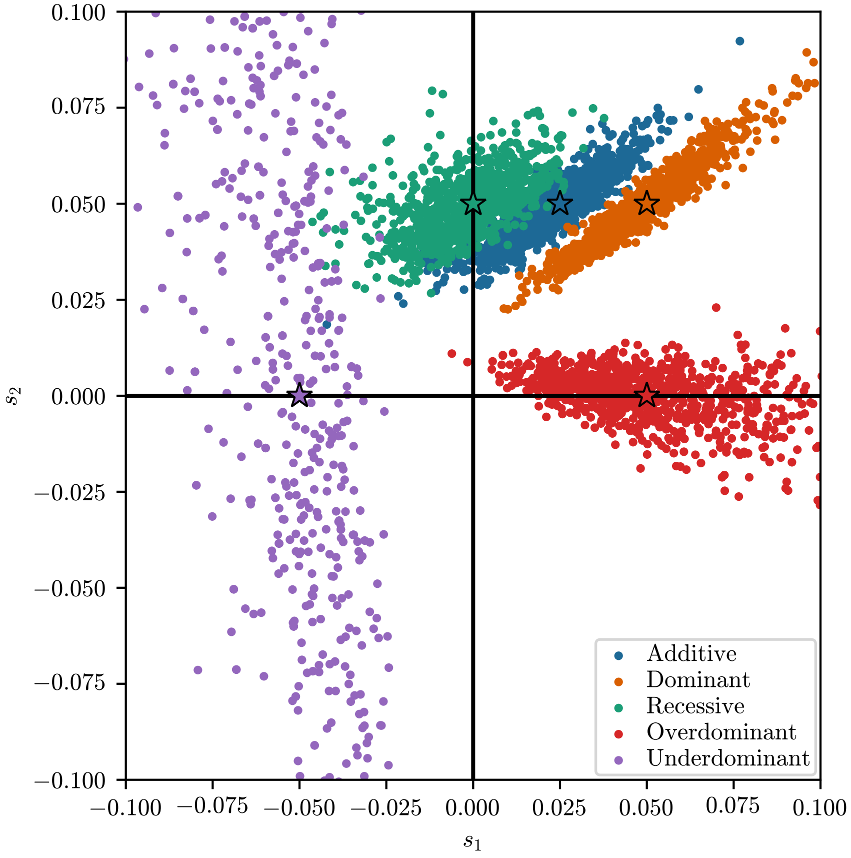


**Figure S.9:** Scatterplot of $(\hat{s}_1, \hat{s}_2)$ estimates for the unconstrained HMM-EM for all constrained modes of selection. *Recessive*, *additive*, *dominant*, and *overdominant* selection are well-estimated. For *underdominant* selection, $s_1$ is well-estimated. For all simulations, the number of generations is $T = 251$ and the initial condition is fixed at frequency $p = 0.25$. The strength of selection is $s = 0.05$ in all cases.

#### S.5 Comparing discretizations of the hidden state space

In this section, we compares the accuracy of the EM-HMM algorithm for different discretizations of the hidden state space in the HMM. In Figure S.10, we vary the number of states, whether or not the hidden states are spaced linearly or using the Chebyshev nodes, and whether the initial frequency of the replicate is estimated or not. 500 hidden states spaced as Chebyshev nodes with the initial condition estimated is the most accurate, though whether or not the initial condition is estimated does not drastically alter the results.

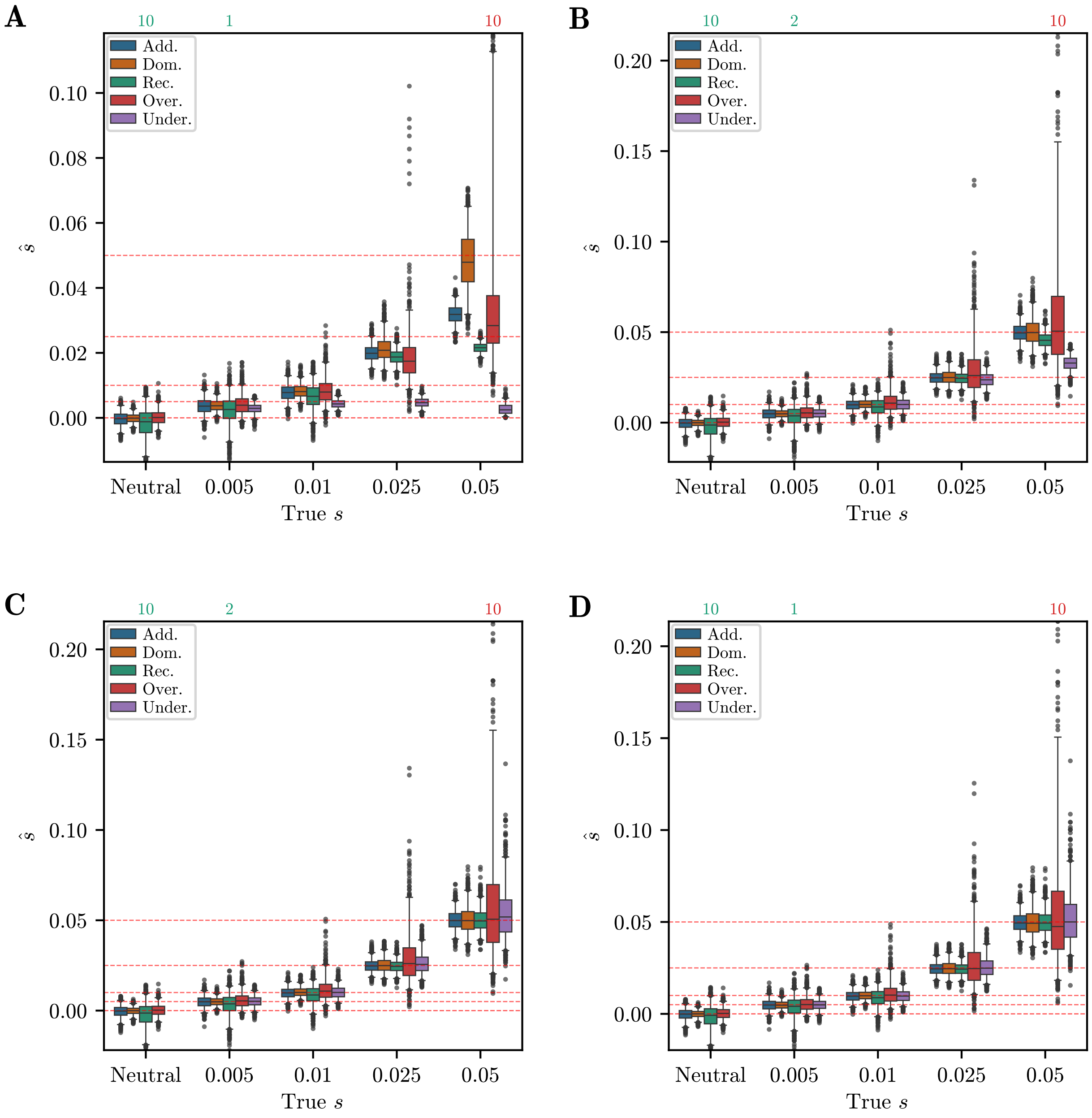


**Figure S.10:** Boxplots of $\hat{s}$ for 1,000 samples under different discretizations of the hidden state space: A) 100 hidden states, equidistantly spaced, initial condition is not estimated, but fixed to true frequency. B) 500 hidden states, equidistantly spaced, initial condition is not estimated. C) 500 hidden states, Chebychev nodes, initial condition is not estimated. D) 500 hidden states, Chebychev nodes, initial condition is estimated. For all boxplots, the datasets are simulated with the number of generations $T = 251$ and an initial condition a fixed frequency at $p = 0.25$.

#### S.6 Results for data-matched simulations with constant $N_e$

In this section, we present an equivalent set of plots to Section 3.2.2 in the main text for the dataset simulated under a constant $N_e = 9{,}715$, matching the $N_e$ estimated from the GB aDNA dataset. As with the IBDNe dataset analyzed in the main text, the sampling scheme, initial frequency, and missingness of each replicate is drawn from the empirical distribution of the GB aDNA dataset. Figure S.11 shows boxplots, AUCs, and Q-Q plots; Figure S.12 shows the corresponding confusion table; and Figure S.13 shows stripplots for the conditional estimates.

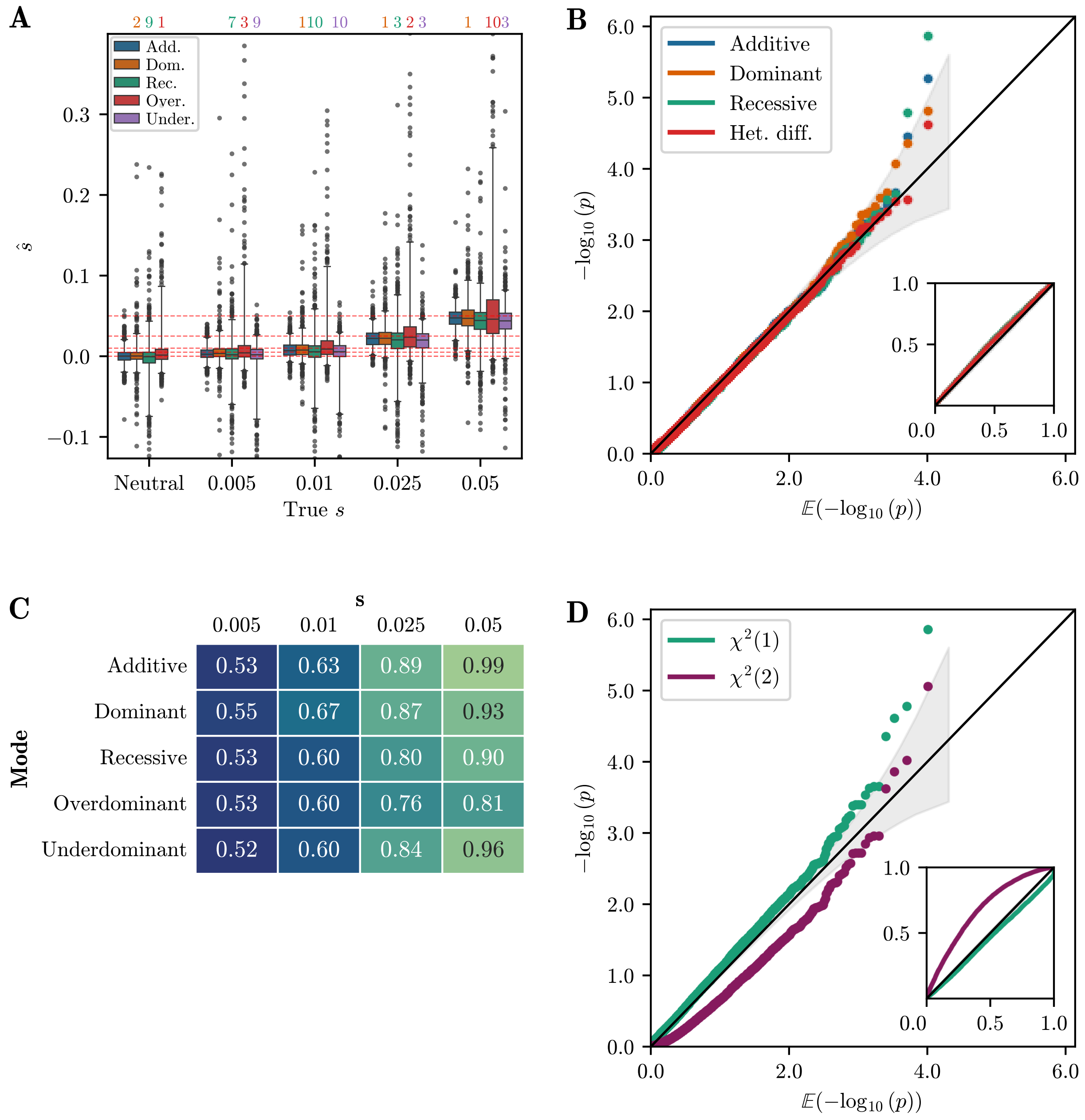


**Figure S.11:** A) Boxplots of MLEs $\hat{s}$ for all one-parameter selection modes. Each boxplot shows 1,000 replicates out of the 10,000 simulated. Whiskers extend to 2.5% and 97.5%-tiles. Estimates are largely unbiased for small $s$, and slightly biased downward for large $s$. B) Q-Q plot of $-\log_{10}$(p-value) against $\mathbb{E}[-\log_{10}$(p-value)] for single-alternative tests obtained using the $\chi^2(1)$ distribution. Inset shows raw p-value against expected p-value. As for the simulated datasets, p-values are well calibrated. C) Table of AUC values for data-matched simulations using likelihood-ratios for each one-parameter selection mode. D) Q-Q plot of $-\log_{10}$(p-value) against $\mathbb{E}[-\log_{10}$(p-value)] for multiple-alternative test statistic $\delta$ using $\chi^2(1)$ and $\chi^2(2)$ distributions. The $\chi^2(1)$ distribution provides the best fit. For all simulated replicates, the number of generations is $T = 125$, $N_e = 9{,}715$ for all generations, and the initial frequency is randomly drawn from the initial frequencies estimated in the GB aDNA dataset.

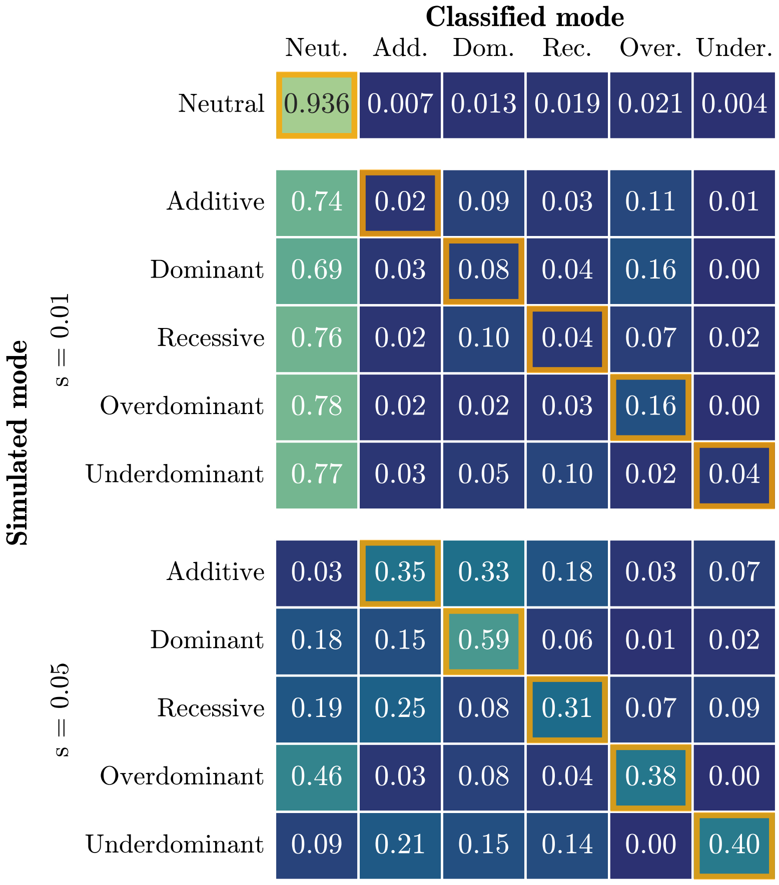


**Figure S.12:** Confusion matrix for procedure to infer selection mode applied to data-matched simulations, using a p-value threshold of 0.05. Each cell represents the fraction of replicates that were classified as a particular mode. Performance is worse than for the simulations in Section 3.1.1 of the main text – the correct model is only inferred for the plurality of replicates for $s = 0.05$. For all simulations, 10,000 replicates are simulated under a given selection mode and strength, the number of generations is $T = 125$, $N_e = 9{,}715$ for all generations, and the initial frequency is randomly drawn from the set of initial frequencies in the GB aDNA dataset.

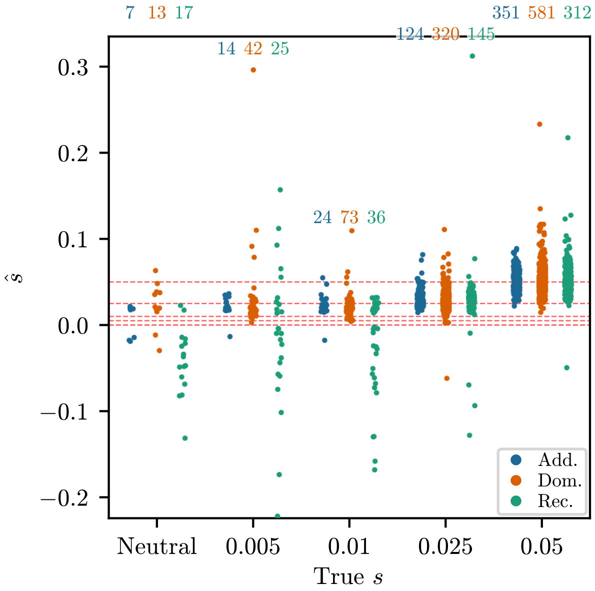


**Figure S.13:** Strip plots of $\hat{s}$ against true $s$, conditioned on inferring non-neutrality for the neutral replicates and on inferring the correct selection mode among multiple alternatives for non-neutral replicates. Each strip plot is for 1,000 replicates randomly chosen from the 10,000 simulated. Number above each strip indicates number of replicates correctly classified. For $s = 0.005$ and $s = 0.01$, the inferred values are biased upwards, due to the "winner's curse". For all simulations, the number of generations is $T = 125$, $N_e = 9{,}715$ for all generations, and the initial frequency is randomly drawn from the distribution estimated in the GB aDNA dataset.

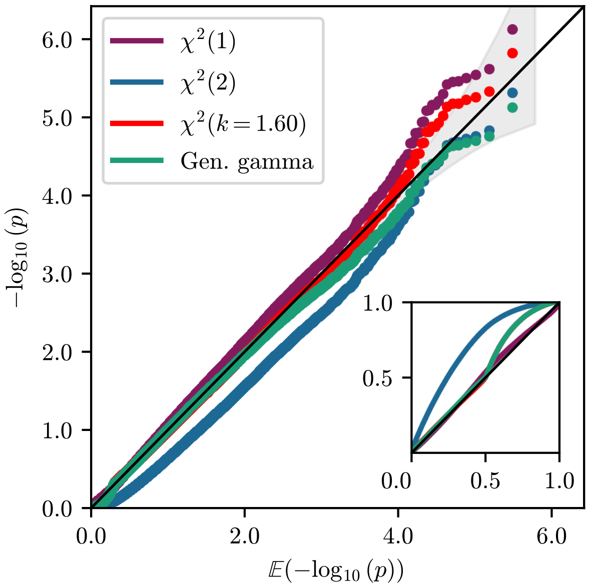


**Figure S.14:** Q-Q plot of $-\log_{10}$(p-value) against $\mathbb{E}[-\log_{10}(\text{p-value})]$ of the $\delta$ statistic for a dataset of 300,000 replicates simulated with the same parameters as the IBDNe dataset. The $\delta$ statistic is plotted for a $\chi^2(1)$ distribution, a $\chi^2(2)$ distribution, a $\chi^2$ distribution where the degrees of freedom has been fit, and a fitted generalized gamma distribution. We fitted to an additional dataset of 10,000 with the same parameters as the IBDNe dataset. As with Figure 9 in the main text, the $\chi^2(1)$ distribution is slightly anti-conservative, but stable in the tail of the distribution.

#### S.7 Additional demographic inference results

This section contains three supplementary plots investigating the demographic inference: Figure S.15 depicts a plot of the $N_e$ at each generation extracted from Browning and Browning (2015); Figure S.16 shows boxplots of estimated $N_e$ values for 25 batches of 10,000 replicates for the IBDNe and const-Ne datasets, analogous to Figure 8 of the main text; and Figure S.17 shows plots of the interpolated log-likelihood surfaces for each batch for the IBDNe and const-Ne datasets.

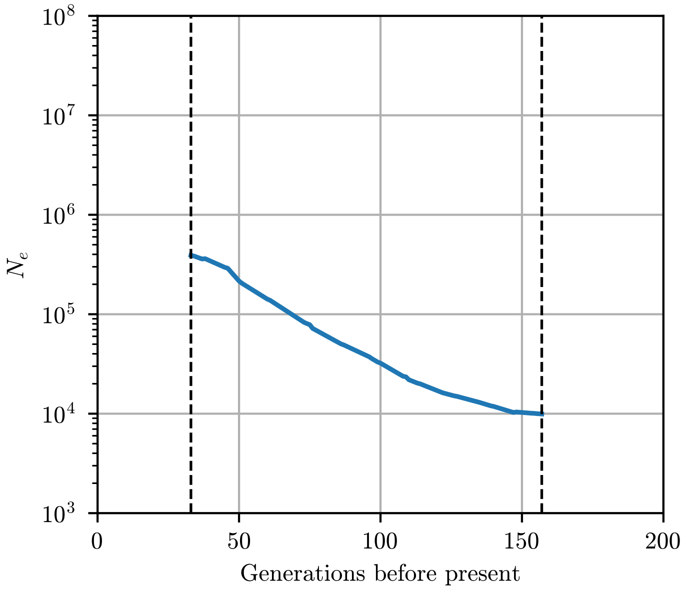


**Figure S.15:** Trajectory of $N_e$ at each generation used to simulate the IBDNe dataset, as extracted from Figure 4A in Browning and Browning (2015). Dashed vertical lines represent the extend of the sampling times in the GB aDNA dataset.

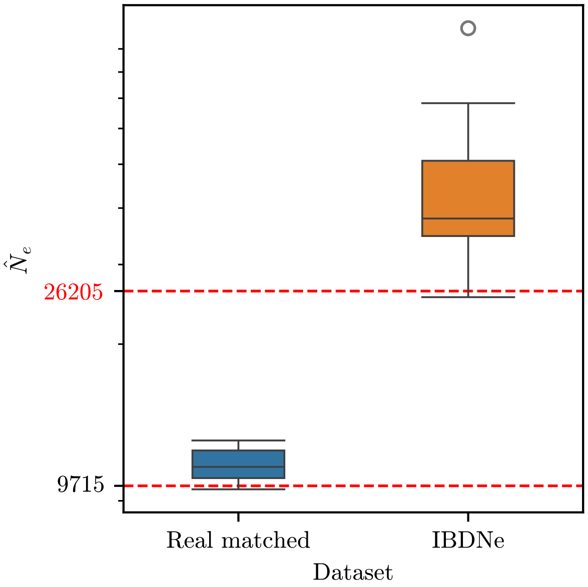


**Figure S.16:** Boxplots of estimated $N_e$ values for 25 batches of 10,000 replicates simulated under the IBDNe and const-Ne conditions. As for the boxplots presented in Figure 8, $N_e$ estimates for the const-Ne data are biased upwards. For the IBDNe dataset, we indicate the harmonic mean of the $N_e$ values at each generation.

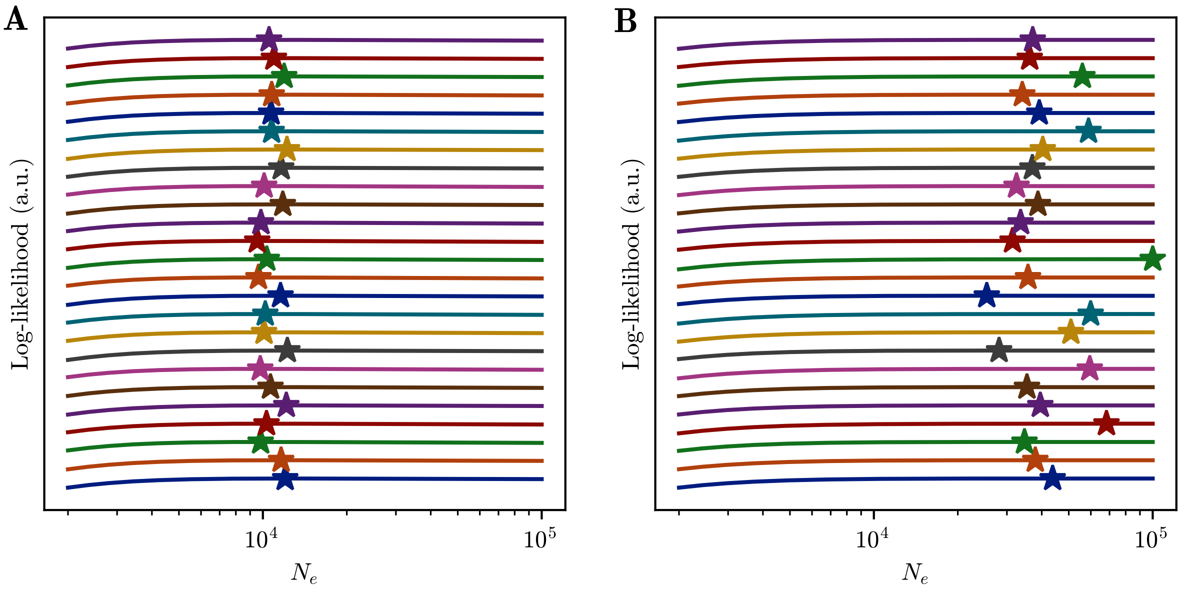


**Figure S.17:** Plots of composite, interpolated log-likelihood surfaces for each of the 25 batches of A) const-Ne and B) IBDNe simulated replicates. Beyond $N_e$ = 10,000, surfaces are very flat, making precise estimation of $N_e$ difficult, and necessitating the conditioning procedure described in Section 3.1.6. Each line represents the sum over 10,000 replicates of log-likelihood calculated at 21 grid points, and interpolated using cubic splines. The star denotes the maximum log-likelihood of this interpolated surface. Lines are evenly spaced for illustrative purposes; log-likelihoods are similar between batches.

#### S.8 Statistics for GB aDNA dataset

In this section, we present plots characterizing the composition of the GB aDNA dataset and some relevant statistics. Figure S.18 shows a map of the location and age of all samples in the dataset used for analysis, Figure S.19 depicts two PCA plots of the ancient samples, one including 9 global and the other 5 European populations from the 1000 Genomes project (Auton et al. 2015), Figure S.20 shows the missingness across SNPs, and Figure S.21 the estimated initial frequencies.

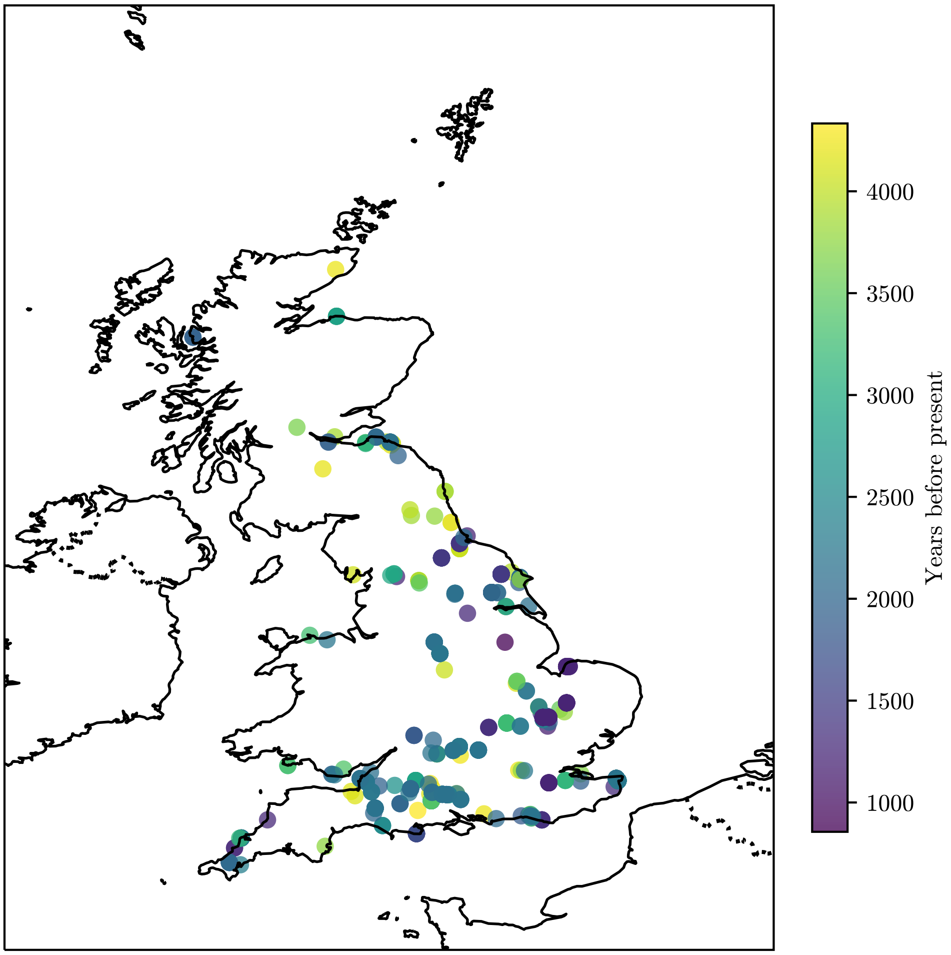


**Figure S.18:** Map of the sample locations within Great Britain, where the color indicates the sampling date before present for all samples used in our analysis. Made with Natural Earth (`https://www.naturalearthdata.com/`, public domain license).

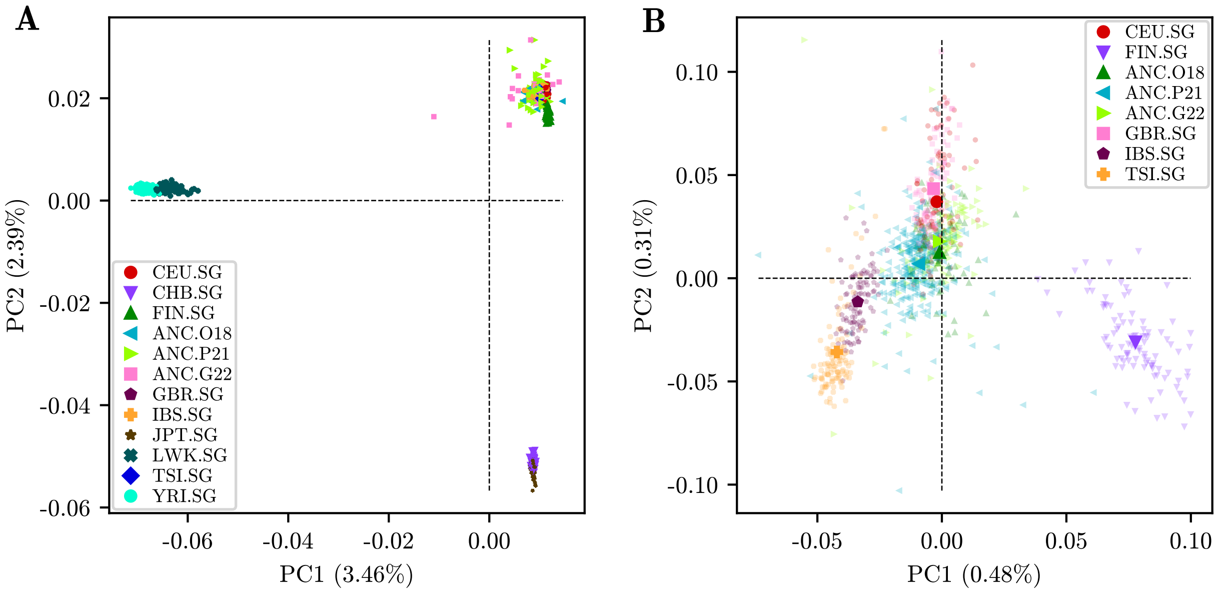


**Figure S.19:** First two axes of a PCA analysis run on the GB aDNA dataset together with A) nine global populations or B) five European populations from the 1000 Genomes project. Populations from the 1000 Genomes project are labeled by the respective population code with the suffix `.SG`. Ancient samples are labeled by the respective publication: `ANC.O18` (Olalde et al. 2018), `ANC.P21` (Patterson et al. 2022), and `ANC.G22` (Gretzinger et al. 2022). Large symbols represent centroids of each population. Ancient samples cluster with the European samples in the global PCA. They cluster close to the CEU and GBR samples in the European plot, and do not exhibit strong structure.

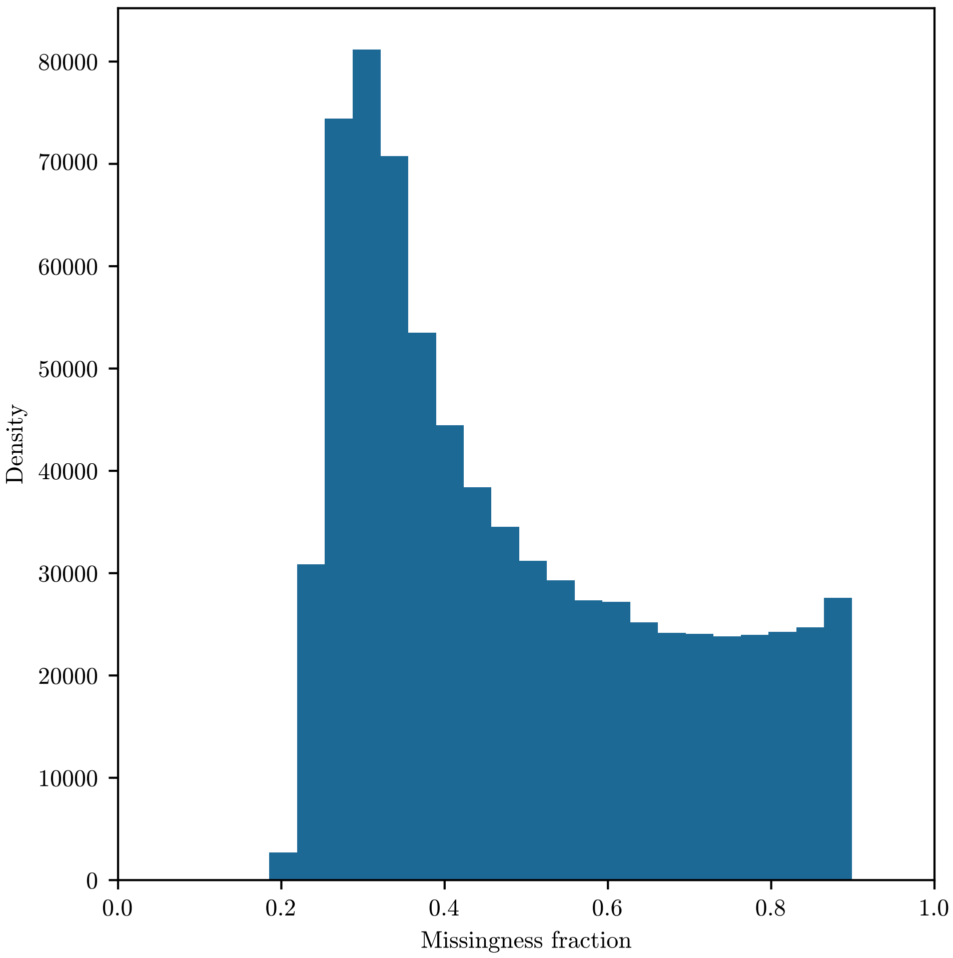


**Figure S.20:** Histograms of the fraction of pseudo-haploid samples with missing genotypes across SNPs in the GB aDNA dataset that pass our filters. Since we filter out SNPs with $\geq$ 90% missingness, the histogram does not extend past 0.90.

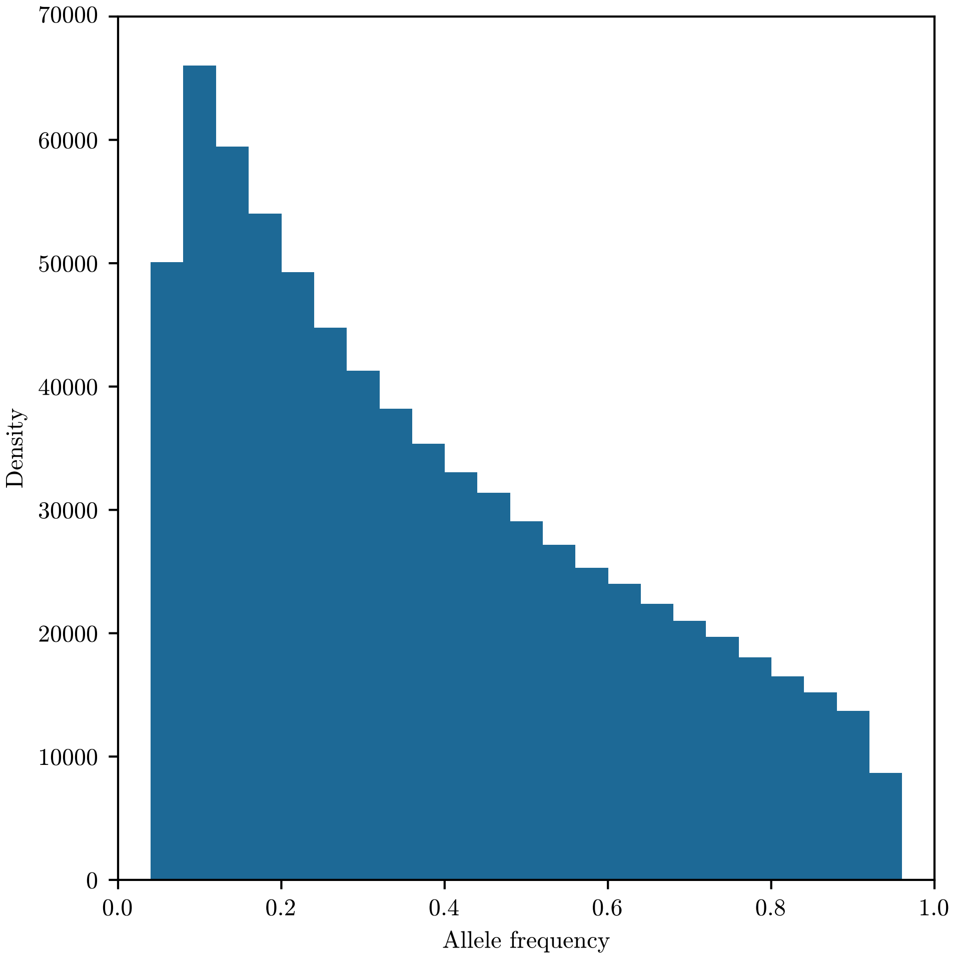


**Figure S.21:** Histogram of the mean of the beta distribution ($\frac{\hat{\alpha}}{\hat{\alpha}+\hat{\beta}}$) estimated as the initial distribution for each SNP in the GB aDNA dataset that passes our filters. This histogram is substantially different from an allele frequency distribution expected at equilibrium under population genetic models, reflecting the ascertainment of the SNPs in the GB aDNA dataset and our criterion to include only SNPs with minor allele frequency $> 5\%$.

#### S.9 Results for GB aDNA dataset

Figure S.22 shows a Manhattan plot of the p-values under the *additive* mode, when we analyze a all samples in the geographic region, combining samples genotyped using the 1240K capture panel and samples genotyped using whole genome sequencing. The plot shows many spurious signals of selection. Figure S.23 depicts a Q-Q plot of the p-values obtained under all one-parameter modes and the multi-alternative test for all SNPs in the GB aDNA dataset (1240K capture only) that pass filtering. A Manhattan plot of the p-values under *dominance* is given in Figure S.24, under *recessive* in Figure S.25, under *heterozygote difference* in Figure S.26, and using the multiple alternative procedure in Figure S.27. Similar to Table 1 in the main text, Table S.1, Table S.2, Table S.3, and Table S.4, show the genomic region identified under *dominance*, *heterozygote difference*, *recessive*, and multiple alternative, respectively. Figure S.28A shows the genome-wide Spearman rank correlation of signed log-likelihood ratio statistics (the log-likelihood ratio is multiplied by -1 if $\hat{s} < 0$) for the top 1% of SNPs (by *additive* p-value) between *additive*, *dominance*, and *recessive* selection. Figure S.28B shows the Spearman rank correlation of the unsigned log-likelihood ratio statistics between *heterozygote difference* and the three aforementioned modes for the same 1% of SNPs.

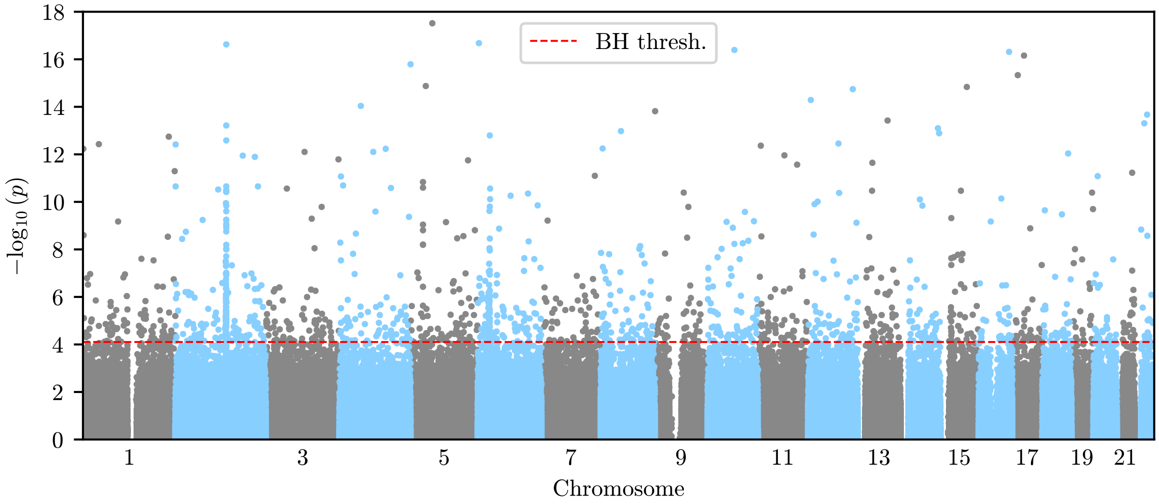


**Figure S.22:** Manhattan plot of the p-values when analyzing a dataset that contains all samples in the geographic region Great Britain, including samples genotyped using the 1240K capture panel, as well as samples genotyped using whole-genome sequencing. Although p-values are generally more significant then using the samples from 1240K capture only, we observe many isolated significant p-values that we deem spurious signals. We thus proceed without the samples genotyped using whole genome sequencing.

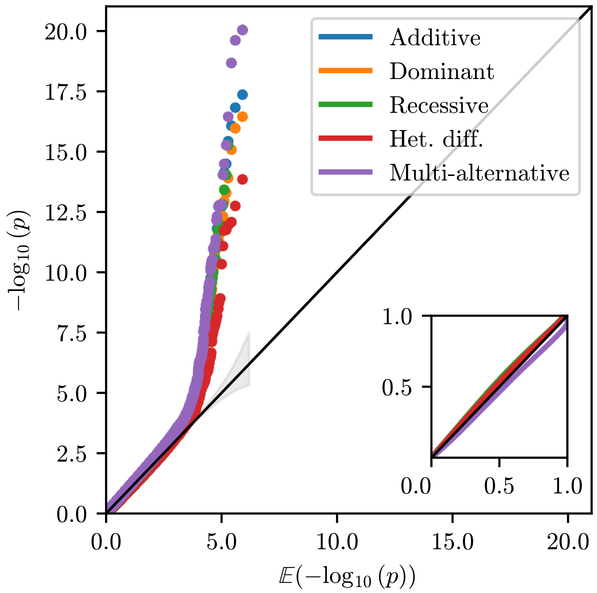


**Figure S.23:** Q-Q plot for the p-values at all SNPs in the GB aDNA dataset, obtained using the single-alternative tests, as well as the test for multiple alternatives, resulting from the procedure to identify the mode of selection. We see an enrichment of low p-values, corresponding to the signals in the data.

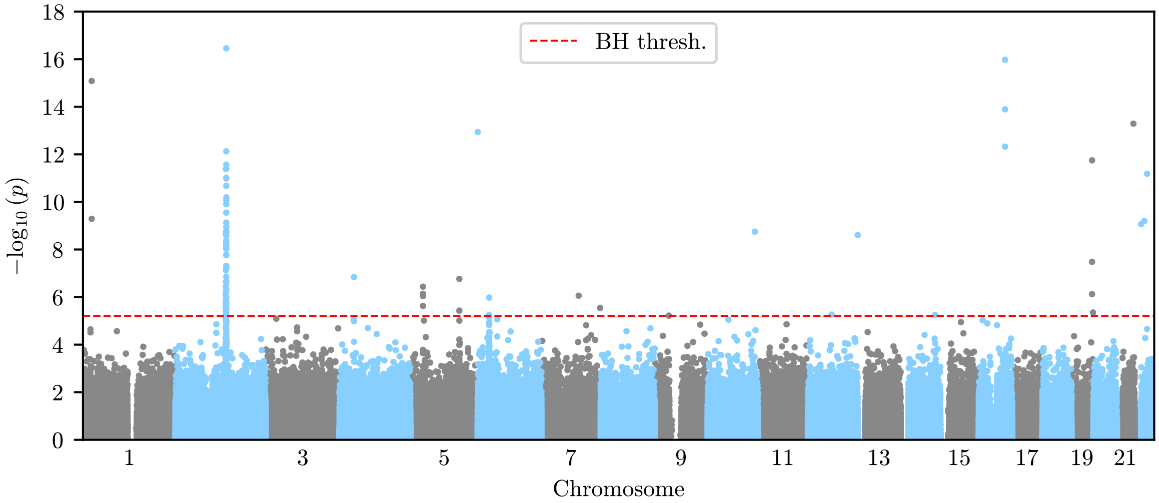


**Figure S.24:** Manhattan plot of the p-values under the *dominant* mode. Regions of significance largely overlap with those under the *additive* mode.

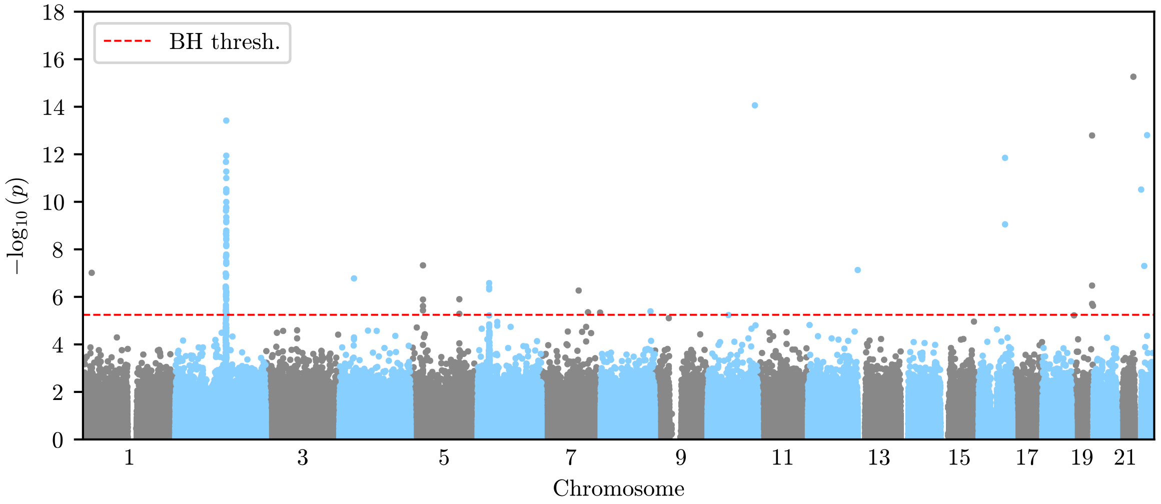


**Figure S.25:** Manhattan plot of the p-values under the *recessive* mode. Regions of significance largely overlap with those under the *additive* mode.

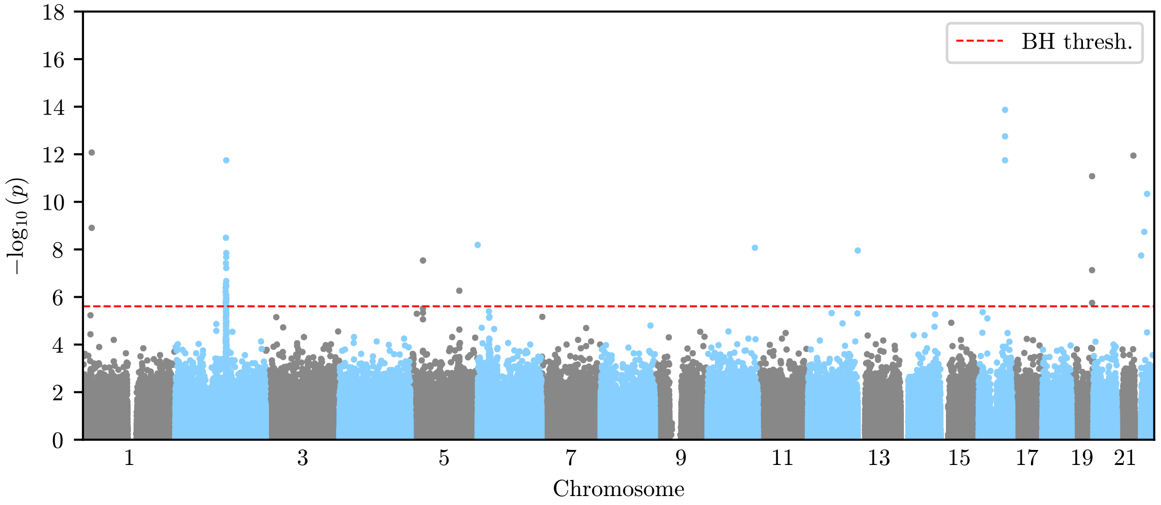


**Figure S.26:** Manhattan plot of the p-values under the *heterozygote difference* mode. In general, the p-values are less significant than under the other single-alternative tests.

| Chr. | Genomic region (hg19) | Gene(s) | Lead SNP | Ref. | Alt. | Raw | Post | $-\log_{10} p_{min}$ | $\hat{s}(p_{min})$ |
|---|---|---|---|---|---|---|---|---|---|
| 2 | 135,161,631-137,087,603 | LCT | rs4988235 | G | A | 62 | 397 | 16.43 | 0.069 (0.050, 0.086) |
| 4 | 38,662,059-38,806,462 | TLR10/1/6 | rs10008492 | C | T | 1 | 58 | 6.83 | -0.045 (-0.066, -0.020) |
| 5 | 33,854,740-34,004,707 | SLC45A2 | rs185146 | C | T | 4 | 51 | 6.43 | 0.127 (0.063, 0.201) |
| 5 | 131,379,656-131,675,864 | SLC22A4 | rs7727544 | C | T | 2 | 94 | 6.76 | 0.034 (0.020, 0.048) |
| 6 | 31,008,598-31,091,197 | HLA | rs2535317 | C | T | 2 | 150 | 5.98 | -0.038 (-0.056, -0.015) |
| 7 | 100,198,386-100,365,613 | TFR2 | rs4434553 | A | G | 1 | 29 | 6.04 | 0.040 (0.023, 0.055) |
| **16 | 70,771,279-71,330,074 | HYDIN | rs79233902 | T | G | 3 | 51 | 15.95 | 0.135** |

**Table S.1:** Genomic regions identified as significant under dominant selection. The asterisks at the HYDIN locus indicate that we believe this to not be a true signal of selection.

| Chr. | Genomic region (hg19) | Gene(s) | Lead SNP | Ref. | Alt. | Raw | Post | $-\log_{10} p_{min}$ | $\hat{s}(p_{min})$ |
|---|---|---|---|---|---|---|---|---|---|
| 2 | 135,167,600-137,085,934 | LCT | rs4988235 | G | A | 22 | 390 | 11.74 | 0.106 (0.056, 0.213) |
| 5 | 33,853,341-34,006,510 | SLC45A2 | rs16891982 | C | G | 1 | 53 | 7.53 | -0.076 (-0.092, 0.131) |
| 5 | 131,329,591-131,665,378 | SLC22A4 | rs7727544 | C | T | 1 | 106 | 6.26 | 0.078 (0.036, 0.167) |
| **16 | 70,771,279-71,330,074 | HYDIN | rs79233902 | T | G | 3 | 51 | 13.85 | 0.138** |

**Table S.2:** Genomic regions identified as significant under heterozygote difference selection. The asterisks at the HYDIN locus indicate that we believe this to not be a true signal of selection.

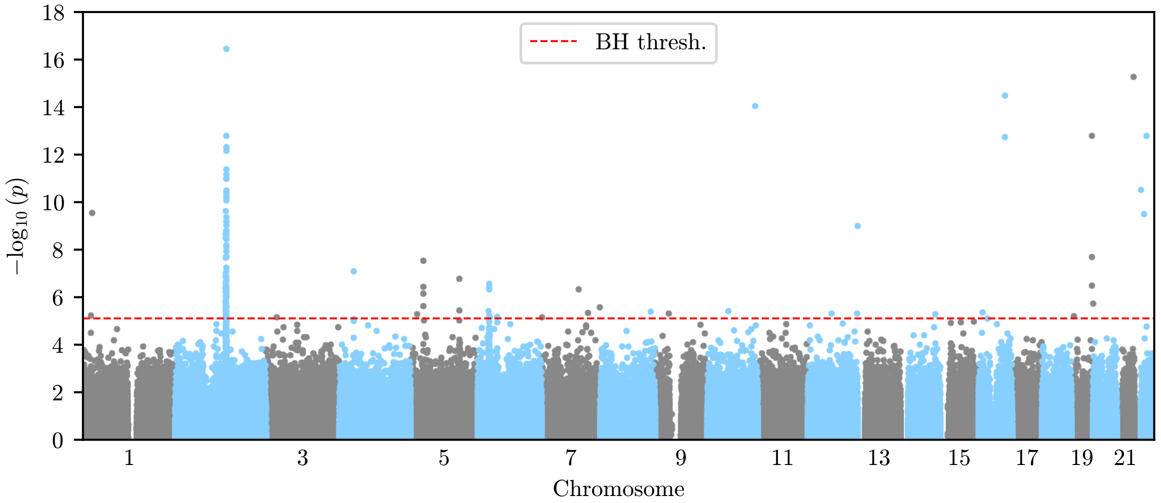


**Figure S.27:** Manhattan plot of the p-values under the multiple alternative procedure. P-values are similar to the *additive* mode.

| Chr. | Genomic region (hg19) | Gene(s) | Lead SNP | Ref. | Alt. | Raw | Post | $-\log_{10} p_{min}$ | $\hat{s}(p_{min})$ |
|---|---|---|---|---|---|---|---|---|---|
| 2 | 135,161,631-137,087,324 | LCT | rs4988235 | G | A | 56 | 395 | 13.41 | 0.124 (-0.085, 0.267) |
| 2 | 137,155,850-137,419,024 | LCT | rs580879 | C | T | 1 | 81 | 5.26 | -0.059 (-0.089, 0.030) |
| 4 | 38,689,031-38,806,462 | TLR10/1/6 | rs10008492 | C | T | 1 | 52 | 6.77 | -0.046 (-0.065, -0.026) |
| 5 | 33,853,341-34,004,707 | SLC45A2 | rs16891982 | C | G | 4 | 52 | 7.32 | 0.035 (0.020, 0.048) |
| 5 | 131,526,736-131,675,046 | SLC22A4 | rs7727544 | C | T | 2 | 63 | 5.89 | 0.054 (0.007, 0.084) |
| 6 | 31,005,726-31,091,197 | HLA | rs2535315 | C | T | 3 | 154 | 6.57 | -0.063 (-0.090, -0.033) |
| 7 | 100,159,567-100,361,675 | TFR2 | rs4434553 | A | G | 1 | 31 | 6.25 | 0.046 (0.015, 0.066) |
| **16 | 70,771,279-71,330,074 | HYDIN | rs79233902 | T | G | 3 | 51 | 20.02 | 0.994** |

**Table S.3:** Genomic regions identified as significant under recessive selection. The asterisks at the HYDIN locus indicate that we believe this to not be a true signal of selection.

| Chr. | Genomic region (hg19) | Gene(s) | Lead SNP | Ref. | Alt. | Raw | Post | $-\log_{10} p_{min}$ | $\hat{s}(p_{min})$ (Sel. type) |
|---|---|---|---|---|---|---|---|---|---|
| 2 | 135,161,631-137,087,603 | LCT | rs4988235 | G | A | 68 | 397 | 16.43 | 0.066 (Dominant) |
| 2 | 137,169,495-137,421,838 | LCT | rs580879 | C | T | 1 | 73 | 5.26 | -0.056 (Recessive) |
| 4 | 38,688,362-38,806,462 | TLR10/1/6 | rs10008492 | C | T | 1 | 53 | 7.07 | -0.044 (Additive) |
| 5 | 33,854,740-34,006,510 | SLC45A2 | rs16891982 | C | G | 4 | 52 | 7.53 | -0.052 (Overdom.) |
| 5 | 131,359,614-131,675,864 | SLC22A4 | rs7727544 | C | T | 2 | 105 | 6.76 | 0.033 (Dominant) |
| 6 | 31,008,368-31,093,699 | HLA | rs2535315 | C | T | 5 | 161 | 6.57 | -0.060 (Recessive) |
| 7 | 100,212,254-100,350,763 | TFR2 | rs4434553 | A | G | 1 | 26 | 6.33 | 0.041 (Additive) |
| **16 | 70,771,279-71,330,074 | HYDIN | rs79233902 | T | G | 3 | 51 | 20.02 | 0.994 (Recessive)** |

**Table S.4:** Genomic regions identified as significant under the multiple alternative procedure. The last column denotes the strength of selection estimated under the one-parameter mode with the highest log-likelihood, as per our mode inference procedure. The asterisks at the HYDIN locus indicate that we believe this to not be a true signal of selection.

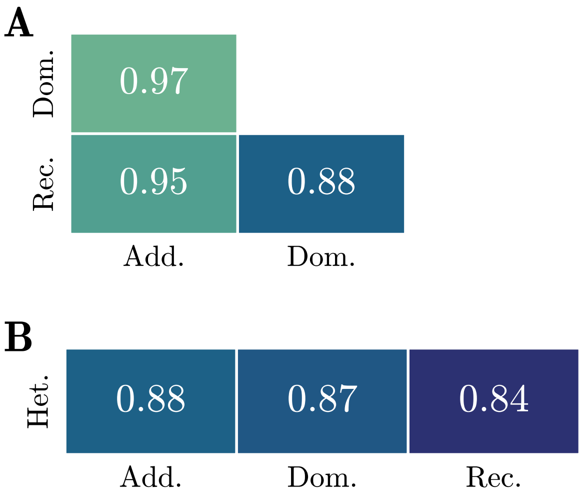


**Figure S.28:** A) Genome-wide Spearman rank correlation coefficients for signed log-likelihood ratios between *additive*, *recessive*, and *dominant* selection modes for the top 1% of SNPs by *additive* p-value. B) Genome-wide Spearman rank correlation coefficients for unsigned log-likelihood ratios between *heterozygote difference* selection and all other one-dimensional modes for the same 1% of SNPs. We observe strong correlation between the different modes.

#### S.10 Permutation testing

In this section, we present Q-Q plots resulting from permuting sampling times, both in the GB aDNA dataset, as well as the const-Ne simulations under neutrality, described in Section S.6. Figure S.29A shows the *additive* p-values of simulated replicates, as well as *additive* p-values computed after independently permuting the sampling times of each sample for each replicate. Figure S.29B shows the p-values obtained from all SNPs in the GB aDNA dataset under the *additive* mode and the p-values obtained after permuting the sampling times associated with each sampled individual. We do not see an enrichment of low p-values, demonstrating that the signal in the data is reliable.

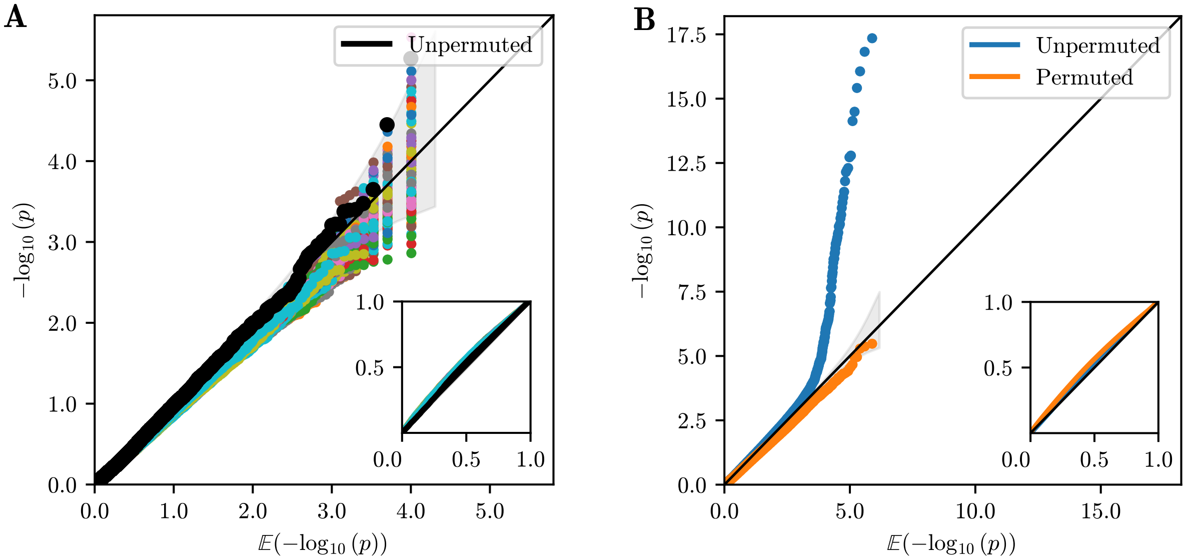


**Figure S.29:** A) Q-Q plot of $-\log_{10}(\mathrm{p})$ against $\mathbb{E}[-\log_{10}(\mathrm{p})]$ obtained under the *additive* mode from the neutral data-matched simulations. The color-coded p-values are obtained from permuting the sampling times for each replicate 100 times. We observe that permuting results in slightly distorted p-values. B) Q-Q plot of $-\log_{10}(\mathrm{p})$ against $\mathbb{E}[-\log_{10}(\mathrm{p})]$ obtained under the *additive* mode for the GB aDNA dataset. The p-values before permutations are enriched for low values, whereas permuting the sampling times removes this enrichment. The permuted p-values fall below the the $y = x$ line, since permuting does not correctly capture the temporal structure of the neutral allele frequency dynamics.

#### S.11 Manhattan and binned frequency plots for significant genomic regions

Figure S.30 to Figure S.37 in this section show, for all regions identified as under additive selection (see Table 1 in the main text), Manhattan plots of *additive* p-values under additive selection for the entire region and binned allele frequency plots for the 20 SNPs surrounding the lead SNP within each region. The color of points and lines in the allele frequency plot reflects the number of SNPs between a given SNP and the focal SNP, with a gradient from red (previous SNPs along the genome) to black (focal) to blue (subsequent SNPs). With the exception of HYDIN, all regions show elevated p-values at surrounding SNPs and correlated allele frequency changes.

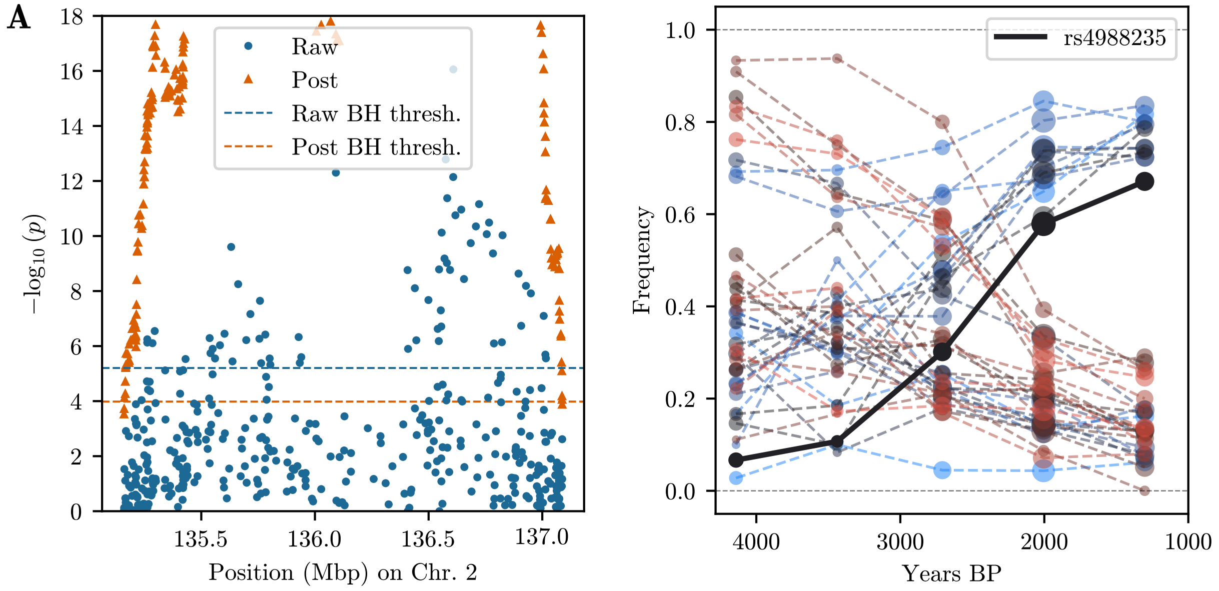


**Figure S.30:** *Additive* Manhattan plot and binned allele frequency plot for the first LCT region on chromosome 2.

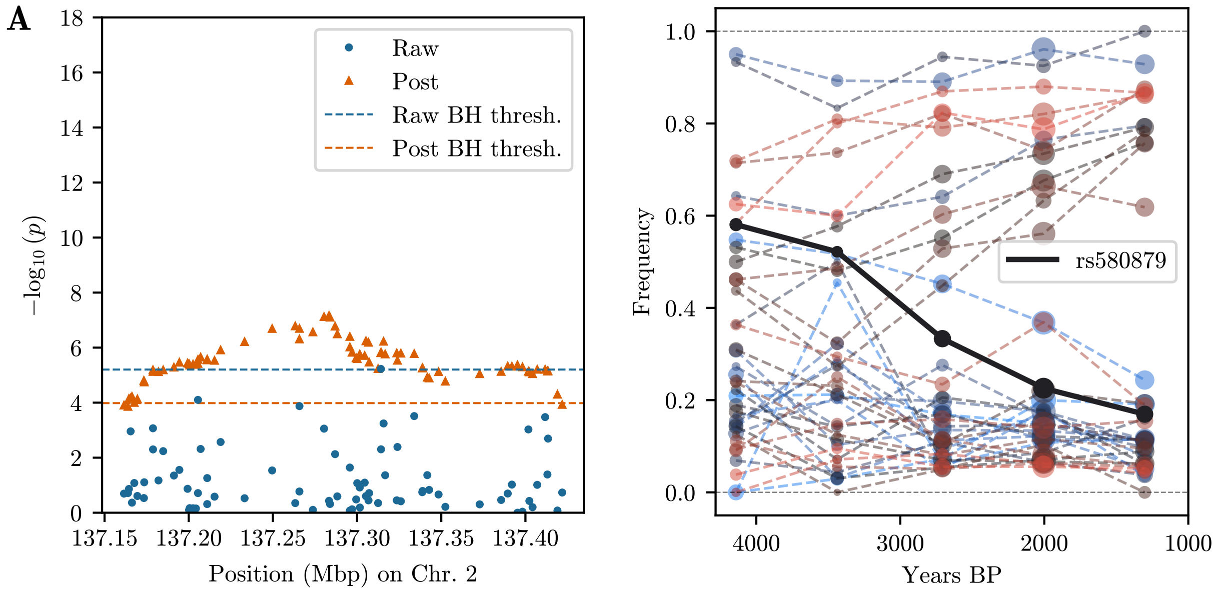


**Figure S.31:** *Additive* Manhattan plot and binned allele frequency plot for the second LCT region on chromosome 2.

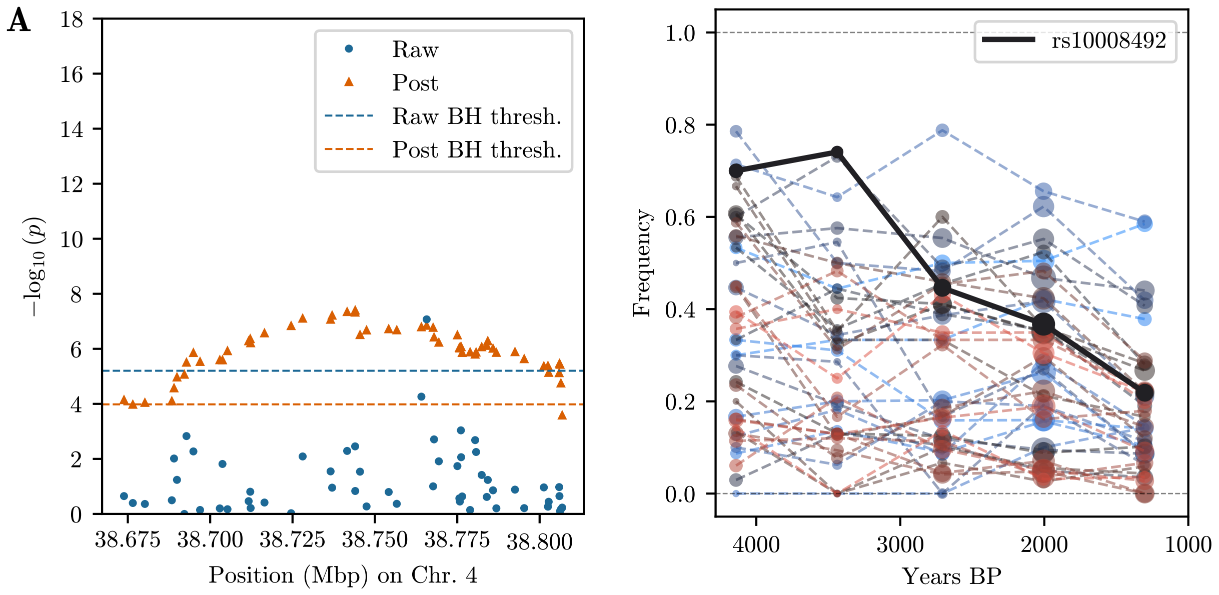


**Figure S.32:** *Additive* Manhattan plot and binned allele frequency plot for the TLR10/1/6 region on chromosome 4.

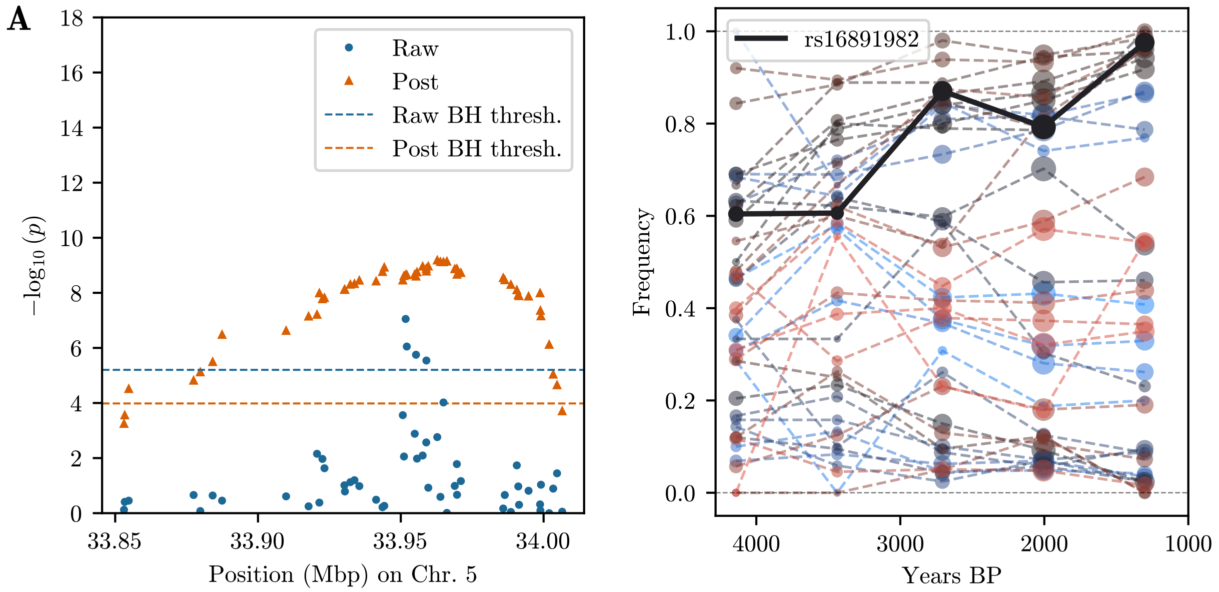


**Figure S.33:** *Additive* Manhattan plot and binned allele frequency plot for the SLC45A2 region on chromosome 5.

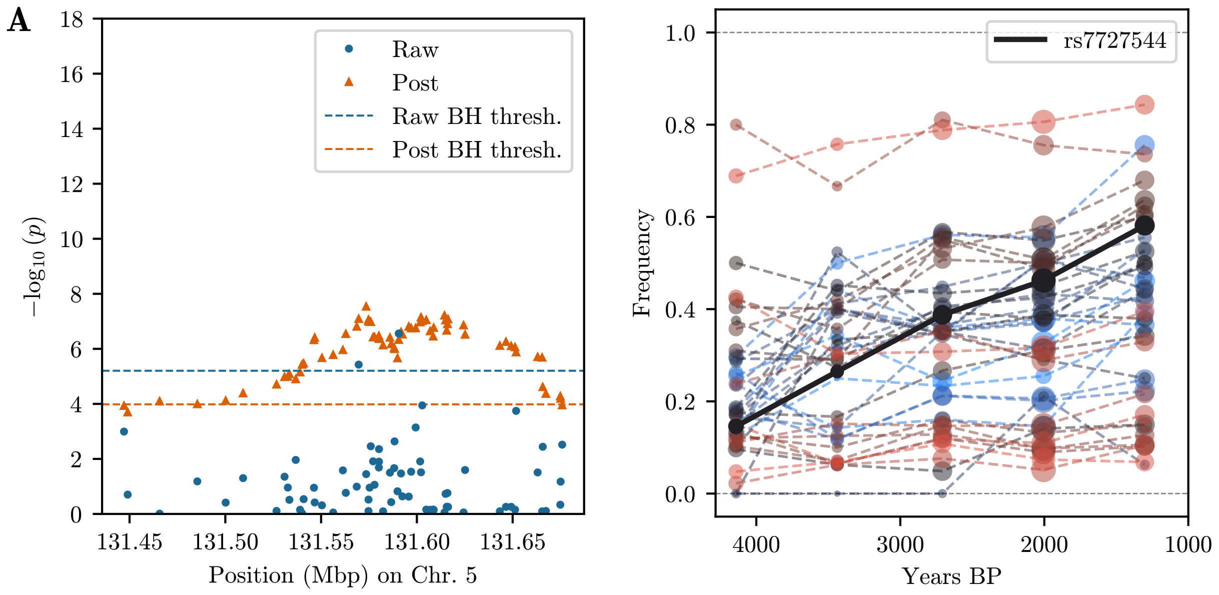


**Figure S.34:** *Additive* Manhattan plot and binned allele frequency plot for the SLC22A4 region on chromosome 5.

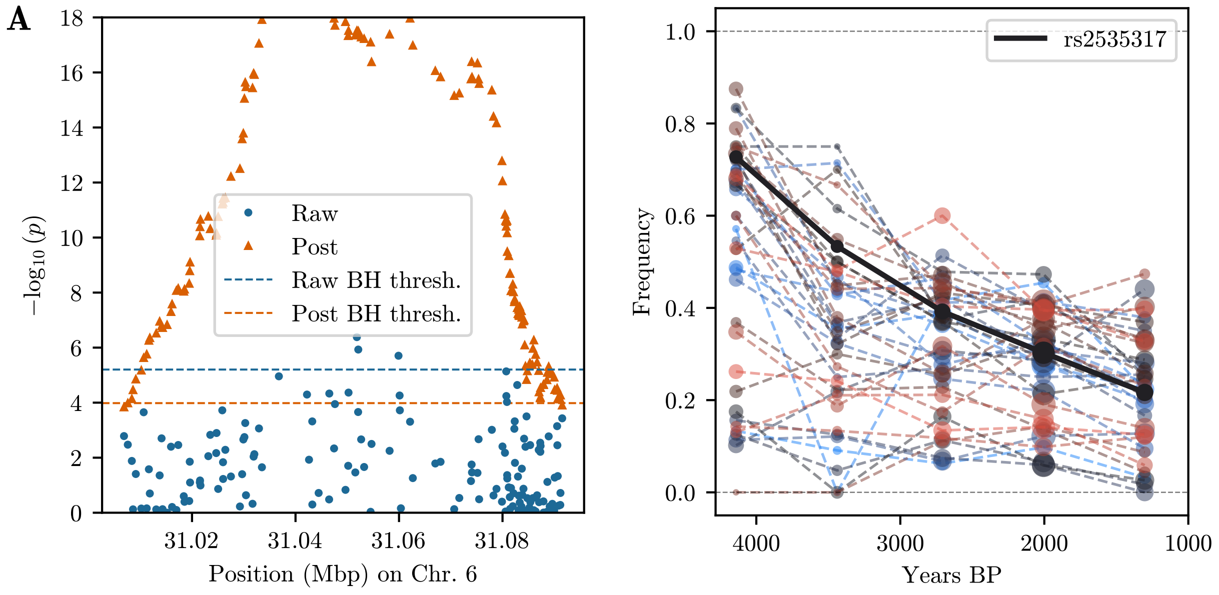


**Figure S.35:** *Additive* Manhattan plot and binned allele frequency plot for the HLA region on chromosome 6.

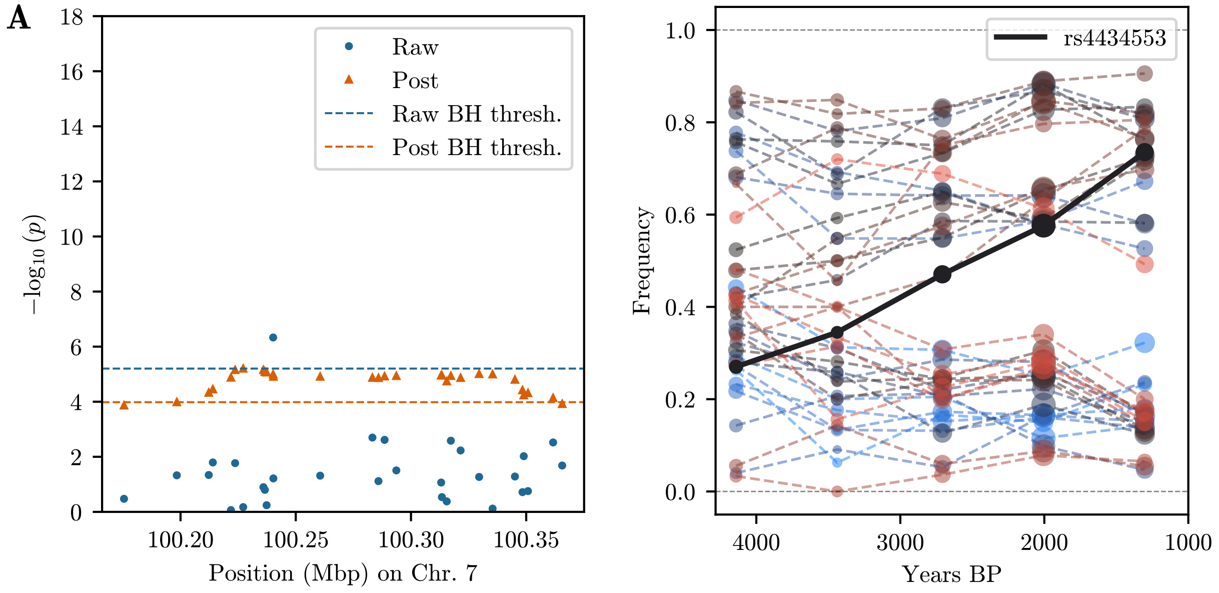


**Figure S.36:** *Additive* Manhattan plot and binned allele frequency plot for the TFR2 region on chromosome 7.

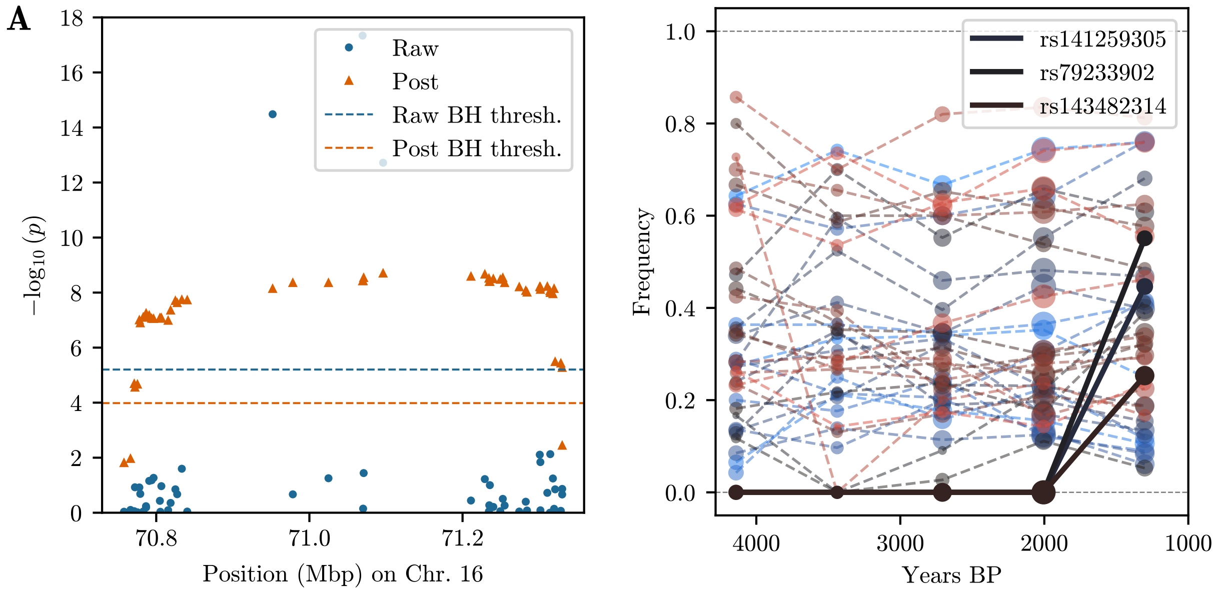


**Figure S.37:** *Additive* Manhattan plot and binned allele frequency plot for the HYDIN region on chromosome 16. All three SNPs with spuriously high p-values are highlighted.

#### S.12 Comparison to Mathieson and Terhorst (2022)

In this section, we present Manhattan plots of p-values under the *additive* mode and plots of binned allele frequency trajectories in the regions that Mathieson and Terhorst (2022) identify as significant, but were not identified in our study: Figure S.38 shows the region around DHCR7 on chromosome 11 and Figure S.39 shows the region around HERC2 on chromosome 15. Our p-values in both regions are low, but the raw p-values do not exceed the BH threshold. Colors in the allele frequency trajectory plots are as in the previous section.

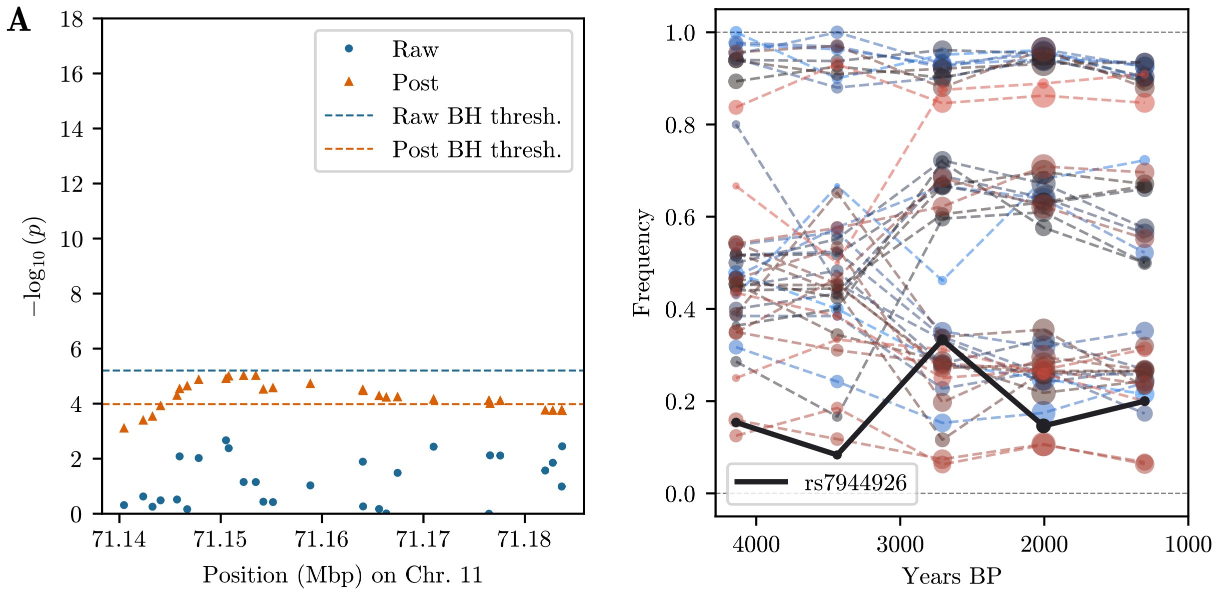


**Figure S.38:** *Additive* Manhattan plot and binned allele frequency plot for the DHCR7 region on chromosome 11.

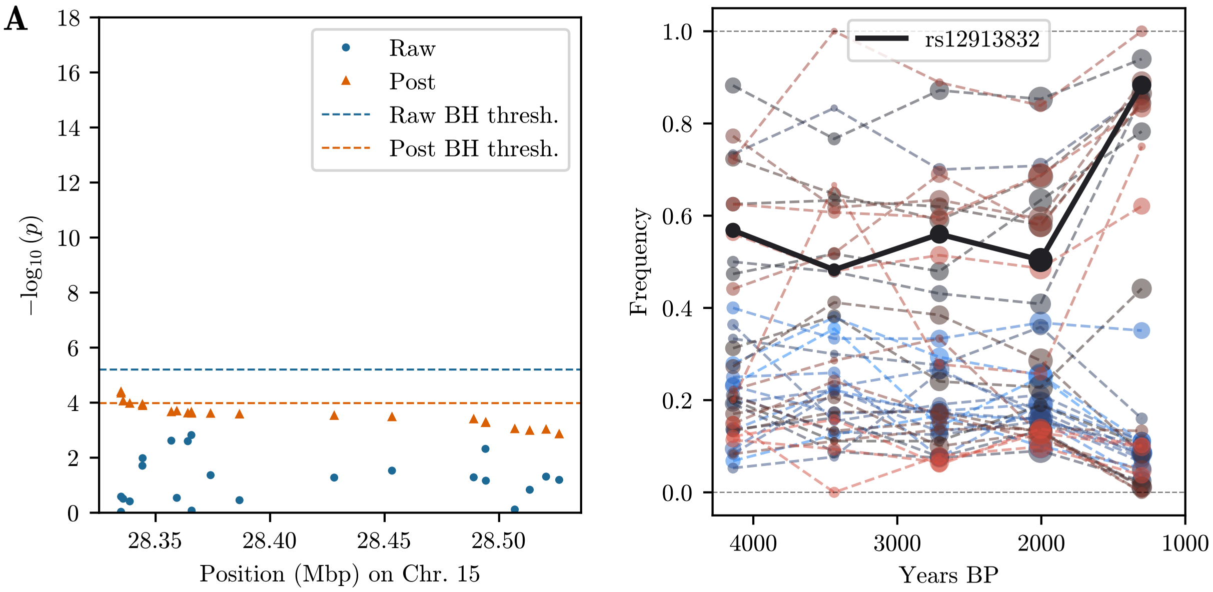


**Figure S.39:** *Additive* Manhattan plot and binned allele frequency plot for the HERC2 region on chromosome 15.

#### S.13 Raw ASIP allele counts

Table S.5 provides the raw data used for the analyses in Section 3.3 of the main text. Original data from diploid samples first genotyped in Ludwig et al. (2009); table directly based on the reproduction in Steinrücken et al. (2014).

| Sampling time (KYA) | Total num. samples | ASIP # derived alleles |
|---|---|---|
| 22.0 | 10 | 0 |
| 15.1 | 22 | 1 |
| 5.7 | 20 | 15 |
| 4.8 | 20 | 12 |
| 3.1 | 36 | 15 |
| 2.5 | 38 | 18 |

**Table S.5:** Table of allele counts for the ASIP locus in domesticated horses. Data originally from Ludwig et al. (2009).

### References


Auton, Adam et al. (2015). "A global reference for human genetic variation". *Nature* 526(7571), pp. 68–74.

Browning, Sharon R. and Brian L. Browning (2015). "Accurate Non-parametric Estimation of Recent Effective Population Size from Segments of Identity by Descent". *American Journal of Human Genetics* 97(3), pp. 404–418.

Gretzinger, Joscha et al. (2022). "The Anglo-Saxon Migration and the Formation of the Early English Gene Pool". *Nature* 610, pp. 112–119.

Ludwig, Arne, Melanie Pruvost, Monika Reissmann, Norbert Benecke, Gudrun A. Brockmann, Pedro Castaños, Michael Cieslak, Sebastian Lippold, Laura Llorente, Anna-Sapfo Malaspinas, Montgomery Slatkin, and Michael Hofreiter (2009). "Coat Color Variation at the Beginning of Horse Domestication". *Science* 324(5926), p. 485.

Mathieson, Iain and Gil McVean (2013). "Estimating Selection Coefficients in Spatially Structured Populations from Time Series Data of Allele Frequencies". *Genetics* 193(3), pp. 973–984.

Mathieson, Iain and Jonathan Terhorst (2022). "Direct Detection of Natural Selection in Bronze Age Britain". *Genome Research*, gr.276862.122.

Olalde, Iñigo et al. (2018). "The Beaker Phenomenon and the Genomic Transformation of Northwest Europe". *Nature* 555, pp. 190–196.

Patterson, Nick et al. (2022). "Large-Scale Migration into Britain during the Middle to Late Bronze Age". *Nature* 601, pp. 588–594.

Steinrücken, Matthias, Anand Bhaskar, and Yun S. Song (2014). "A Novel Spectral Method for Inferring General Diploid Selection from Time Series Genetic Data". *The Annals of Applied Statistics* 8(4), pp. 2203–2222.

Watterson, G. A. (1982). "Testing Selection at a Single Locus". *Biometrics* 38(2), p. 323.
